# Navigational strategy dictates hippocampal representation of space in an everyday memory task

**DOI:** 10.1101/2025.05.10.653115

**Authors:** Francesco Gobbo, Rufus Mitchell-Heggs, Adrian J Duszkiewicz, Elena Faillace, Dorothy Tse, Nuria Garcia-Font, Omer Hazon, Kathrine Clarke, Patrick A Spooner, Adana Keshishian, Mark Schnitzer, Simon R Schultz, Richard GM Morris

## Abstract

Hippocampal activity provides a representation of the world around us, as well as our position within that world. However, it is not known if and to what extent the chosen navigational reference frame can influence hippocampal representations during memory-based tasks, including those focused on future activity. Here, we develop and employ two naturalistic, carefully controlled variants of the everyday memory task to model the use of egocentric and allocentric coordinates in the same arena. By recording hippocampal neural activity through miniature microscopes in male rats performing each of the two tasks, we uncover differences in the representation of space, and in the features of memory-based action planning. By also deploying optogenetic inactivation during navigational decision making, we find that hippocampal representations observed during the planning phase are necessary for solving the allocentric, but not the egocentric version of the task. Overall, our findings reveal a functional link between non-local hippocampal representations and allocentric navigation.

## Introduction

A longstanding challenge in the neurobiology of spatial memory is understanding how neural representations of space contribute to goal identification and navigation. The hippocampus has long been known to contain cells that are selectively active when an animal occupies or passes through a specific location in space (O’Keefe & Dostrovsky, 1971). Place cell firing contributes to an allocentric (world-centered) representation of the immediate environment, with the contribution of other cell types including head-direction cells (Taube *et al*., 1990) and grid cells (Fyhn *et al*., 2004; Hafting *et al*., 2005). A fundamental logical problem, however, is how an animal in one place (Place A), seeking to navigate to a distal Place B but not to Place C, can access information about these possible destinations (Morris, 1991; Knierim, 2009).

While the predominant firing of place cells generally occurs in their place fields, particularly during free exploration, out-of-field firing can occur during task performance, albeit at the cost of spatial ambiguity (Poucet & Hok, 2017). Using calcium (Ca^2+^) imaging in behaving animals performing a demanding everyday spatial memory task (Wang *et al*., 2010), we have recently shown that spatially responsive cells corresponding to distal target locations also fire at the outset of navigation when animals are deciding on the correct destination (Gobbo *et al*., 2022). A key question is then what information is activated in the hippocampus when animals are in the startbox making their decision? How are these nonlocal representations linked to the animal’s choices?

A further issue is whether the cognitive strategy employed in the task affects how neural representations are accessed and used. Navigation can be achieved in diverse ways – using allocentric, egocentric and/or path-integration representations. The hippocampus has access to afferents from the entorhinal cortex conveying both allocentric and egocentric information (Wang, Chen, *et al*., 2020). In many real-world situations, both representational systems are likely to be active and can operate synergistically such that their differential role in guiding behavior can be complex.

For example, using an exclusively allocentric strategy, animals may form a representation of a distal target location and then compute an appropriate trajectory to it. Alternatively, using a strictly egocentric strategy, they may learn to head in a particular direction, and have egocentric representations from previous successful trips. Third, the animal may keep track of its translational and rotational movements to compute a return vector to a starting location and utilize this path-integration mechanism to aid aspects of navigation.

To elucidate how the cognitive strategy being deployed affects neural representations, we developed two modifications of our everyday spatial memory task to dissociate the use of allocentric and egocentric navigational strategies. Using Ca^2+^ imaging with miniature microscopes (miniscopes), we monitored the neural activity of pyramidal cells in the CA1 subfield of the dorsal hippocampus at the outset of navigation. We found that animals using an egocentric strategy had a poorer representation of space compared to those performing allocentrically. Furthermore, we also observed that optogenetic inactivation of hippocampus dorsal CA1 before the onset of navigation caused animals to make navigational errors when performing an allocentric but not an egocentric task.

## Results

### Rats can use either egocentric or allocentric navigation to retrieve food rewards

The experiments to be described were conducted in two separate phases. First, we recorded the neural activity in the hippocampus of rats in an allocentric and, separately, an egocentric task, and observed if, and to what extent, the strategy affected the spatial representation of goals, particularly during planning. Then, in a second experiment, we examined the differential impact of optogenetic inactivation between the two navigational strategies.

We employed Ca^2+^ imaging to record the activity of dorsal CA1 neurons in freely moving rats performing behavioral tasks (see Methods). The tasks revolved around everyday memory in which information is encoded incidentally, and because of this, it is often rapidly forgotten (Morris, 2006; Gobbo *et al*., 2022; Tse *et al*., 2023). Each day, food-restricted rats were first allowed to explore a large square ‘event arena’.

The arena is so named because ‘events’ can happen in it, such as the act of finding a sandwell containing food reward on sample and choice trials (Fig. 1a). On sample trials, rats had the opportunity to learn which one of six possible sandwells contained a hidden food pellet; their memory of the correct sandwell was later tested in choice trials after 90 min. The behavioral procedure differed between allocentric and egocentric training protocols. In the allocentric case, trials started from one of three possible startboxes (named East, South, and West), and retrieved pellets had to be taken to the goal box (North box) where they could be eaten. In the egocentric protocol, all trials within a single session started from the same startbox (North, East, South, or West) and ended with the animal returning to this same starting location (Fig 1b, see Methods).

**Figure 1:**
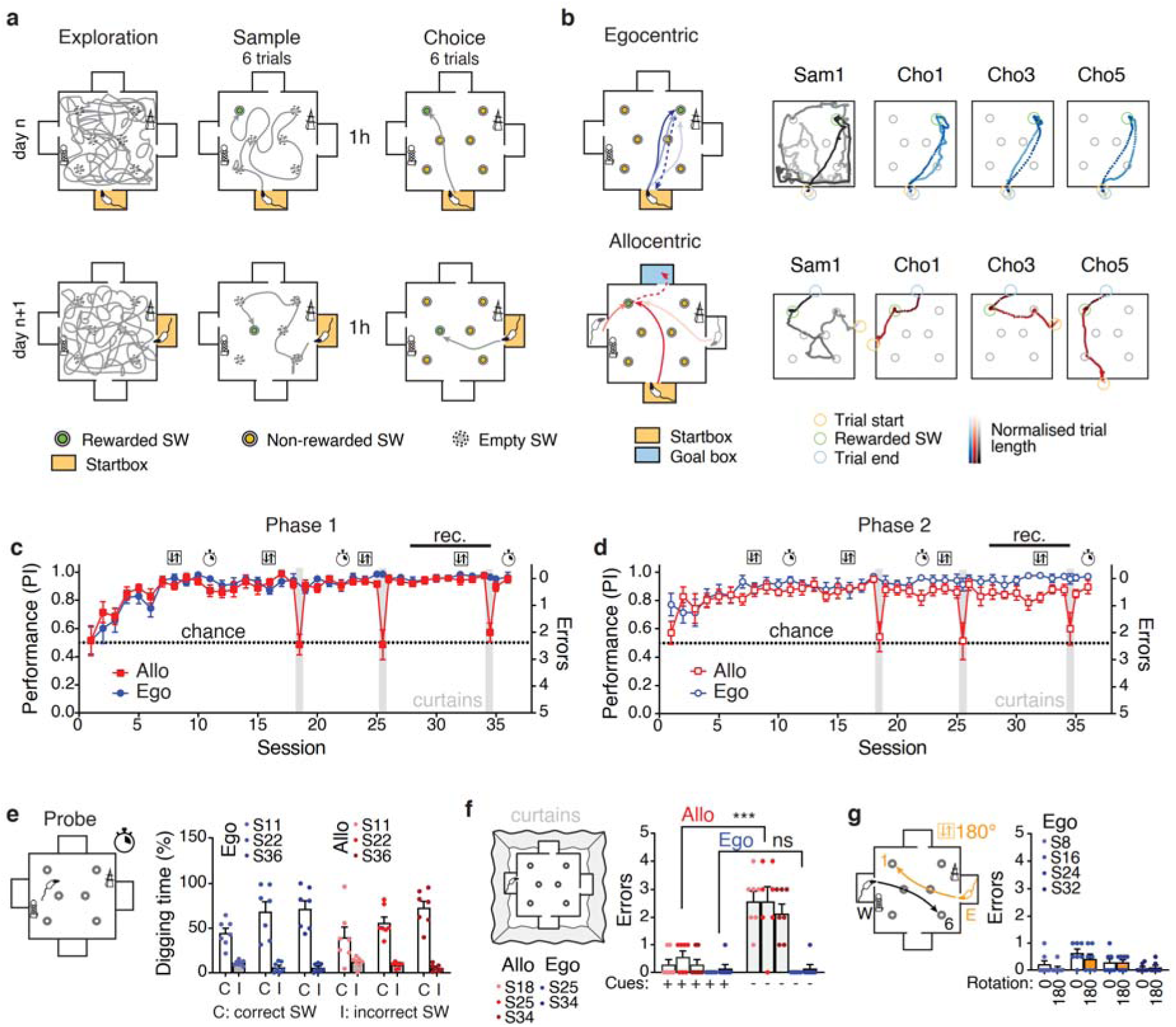
Establishing behavioral protocols that promote the use of allocentric or egocentric navigation in solving the everyday memory task. a) In the everyday memory task, animals were allowed to exit a startbox (orange) and then given 2 mins (extended to 20min in recorded sessions) to explore the arena. This served as daily familiarization and recording of the spatial fields of neurons. There were then 6 sample trials starting from either East (E), South (S) or West (W), whose aim was to allow the animal to locate and dig up food from a single sandwell (SW). These were followed by 6 choice trials in which both choice accuracy (Performance) and latency to find the reward were measured. Note that the correct sandwell changed across days (e.g. day n to day n+1). b) The protocols for egocentric (Ego)- and allocentric (Allo)-trained animals differed in the use of specific startboxes and cue availability in the arena (see Methods). Sample and choice trials in the Egocentric training involved leaving and returning to a single daily startbox; Allocentric training involved 3 blocks of trials starting from E, S, or W in a random combination (odd trials). The animal then carried the food to the goal box and then returned from there to the same sandwell for one further reward (even trials). Representative paths are shown for the first sample trial (Sam1) and 3 odd-number choice trials (Cho1, 3 and 5). c) Phase 1: The animals learned a new location each day in both the egocentric and allocentric protocols. Their performance over 36 daily sessions showed that the animals rapidly learned both tasks and assumed asymptotic levels of performance by Sessions 10-12 (average Performance Index in S28 to S34: Ego 0.96±0.01 and Allo 0.95±0.03 (mean±st.dev); n=7 animals per group). Curtains were pulled to surround the arena to occlude extra-arena cues on S18, S25 and S34. The Performance dropped to chance for the allocentric trained animals, reflecting their reliance on extra-arena cues. Here and in d), the clock symbol denotes sessions where a Probe test was run, whereas the boxed double arrows when a 180°rotation test was run. d) Phase 2: After the crossover, in which animals that had previously been allocentric-trained were switched to the egocentric protocol, and vice versa, performance was broadly equivalent (Performance Index from S28: 0.95±0.05 and 0.85±0.07 respectively). e) In Probe tests, no reward was available in the correct Sandwell to control for olfactory cues, and the fraction of time spent digging at the correct Sandwell is measured. In both groups, we observed focused and persistent digging at the correct Sandwell. f) The impact of occluding extra-arena cues was that allocentric-trained animals fell to chance (circa 2.5 errors), whereas egocentric-trained animals were unaffected. g) Comparison of error scores in egocentric-trained animals with a 180° rotation of the required trajectory. Sessions 8 and 16 were run with curtains, while Sessions 24 and 32 in presence of intra- and extra maze cues. The error rate remained at less than 1. Bars are Means±sem, points are animals. ***P<0.001 one-way ANOVA, Šídák’s multiple comparisons test.

Rats learned both strategies well and reached a steady Performance Index (PI, see Methods) in approximately 12 sessions (Fig. 1c). At asymptote, the PI ranged between 0.85 and 0.95, and median latency to find the correct sandwell on choice trials was 8.8s and 7.9s for egocentric- and allocentric-trained animals respectively (Supplementary Figure S1a). In Phase 1, rats were assigned to one of the two behavioral protocols and, after completion of this Phase, crossed over to the other behavioral protocol (Phase 2, see Methods).

Several tests were conducted to establish whether distinct navigational strategies were being used, as anticipated (Supplementary Figure S1b). First, probe trials were conducted to exclude the possible use of olfactory cues (Fig 1e). As expected, rats dug significantly longer at correct sandwells (C) than at the incorrect ones (I) (two-way repeated measures ANOVA; C/I factor F(1,12)=118.4 P<0.001 for the egocentric group, C/I factor F(1,12)=45.62 P<0.001 for the allocentric group).

Second, extra-arena and intra-maze cues were removed on selected choice trials (using masking curtains to obscure extra-arena cues; sessions 18, 25 and 34, see Methods). As expected, the performance of allocentric-trained animals declined to near random search, indicating their reliance on the use of extra-arena information (two-way repeated measures ANOVA Cues/Curtains factor F(1,12)=37.48 P=0.0003). In contrast, egocentric-trained animals performed equally well with and without cues (two-way repeated measures ANOVA; Cues/Curtains factor F(1,12)=0 P>0.99). Third, egocentric training required animals not to rely on available cues by means of 180° rotations of the start and goal positions with respect to the cues and the arena. For instance, animals trained to run from the West startbox to a sandwell positioned to the animal’s right at a distal point in the arena performed just as well when trials were started from the East to run to the symmetrically rotated location (Fig. 1g). Fourth, we tested if in choice trials, allocentric-trained animals could identify the correct sandwell when starting from a startbox not experienced in the prior sample stage. This was indeed the case (Supplementary Figure S1c), demonstrating their reliance on an allocentric map. Performance in probe trials, cue-masking and 180°-rotation tests were similar after the within-subjects crossover in phase 2 (Supplementary Figure S2).

### Miniscope recording highlights subtle differences in space representation

Using miniscopes, we then recorded the Ca^2+^ activity of CA1 neurons while the animals were performing one of the two behavioral variants of the event arena task during S28-34, and aligned the cell activity across these sessions (Fig. 2a,b and Supplementary Figures S3-S8). We observed similar properties of spatial representation when rats were trained in either behavioral variant, with place cells present in both groups (Figure 2c and Supplementary Figure S9). However, allocentric-trained animals displayed a higher proportion of place cells compared to egocentric-trained ones (Fig. 2d, Linear Mixed Model Phase z=-0.88, P=0.93, Task factor (Ego/Allo) z=-2.3 P=0.021, n=8 animals). Overall, we observed that cells from egocentric-trained animals showed a higher number of fields (Fig. 2e, Mann-Whitney test, P<0.001). This was confirmed by Hierarchical Bootstrap analysis (Saravanan *et al*., 2020) where we resampled these values for an even number of cells for each animal (Fig. 2d and 2e inset are means calculated by Bootstrap; Percentage of Place cells d=1.46, P=0.048, and Number of Place Fields d=-1.26, P=0.070). Statistical significance was found to be robust for the range of cells used in our analysis (Supplementary Figure S10). We observed no significant difference in the size of place fields (Supplementary Figure S10 and S11, Bootstrap analysis of means differences d=0.32, P=0.37) or dependency on the animals’ direction in the two groups (Fig. 2f and Supplementary Figure S12, Bootstrap analysis of means differences d=-0.55, P=0.288). Similarly, day-to-day stability (Fig. 2g, Linear Mixed Model Ego/Allo z=-1.025 P=0.305) as well as across days (Supplementary Figure S13) was not significantly different between the two groups.

**Figure 2:**
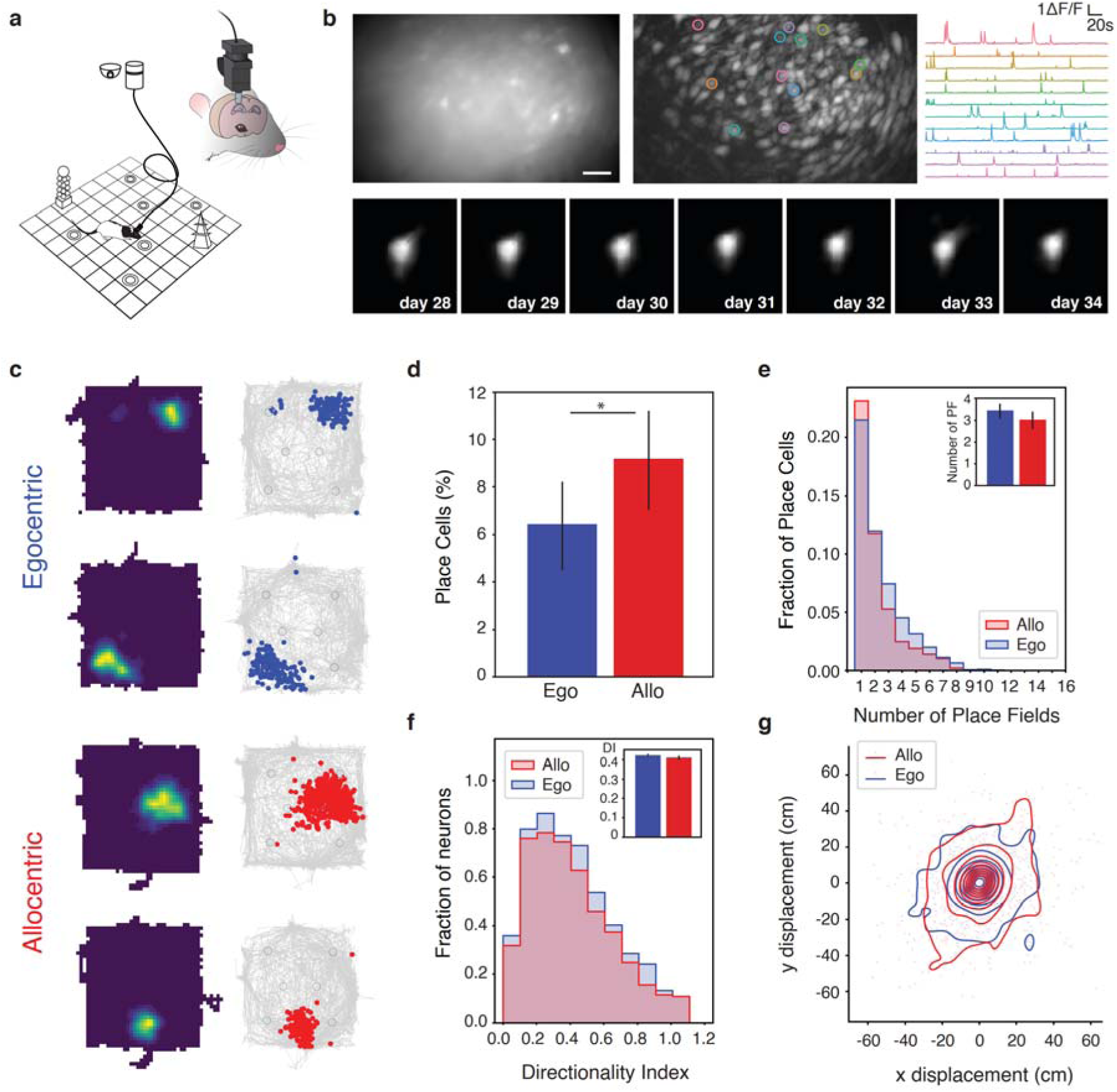
Miniscope Calcium imaging in the event arena and space representation. Schematic of the setup. Rats were able to move freely in the 1.6m-wide arena connected to the miniscope via a cable and a commutator to enable rotation. The rats’ behavioral activity was monitored with a high-resolution camera on the ceiling. b) Representative field of view of CA1 pyramidal cells through the miniscope lens and projected ΔF/F, scale bar 100 μm. On the left, representative ΔF/F traces from a number of cells indicated by circles of corresponding color in the middle panel. Below, a representative cell image identified across all recording sessions (CNMFe cell profile) demonstrating correct alignment of the cell set across days. c) Spatial fields of representative place cells identified during Exploration. In each pair, the left graph is the event density map, and the right graph the position of the events (dots) superimposed on the animal spatial occupancy over the exploration sessions (gray). Larger black circles represent the position of the sandwells. Maps are from two representative animals trained on the egocentric (top, blue events) and allocentric task (bottom, red events). d) Fraction of place cells identified in the two groups, showing a higher number in allocentric-trained animals. Bars are mean ± 25^th^-75^th^ distribution of resampled means, n=9 animals. Values are Bootstrapped to uniform animal contribution regardless of number of cells (see Methods) e) Number of place fields (PF) for place cells identified in the two experimental groups. Cells recorded from egocentric-trained animals had a higher number of PFs. Only PFs larger than 10 cm^2^ are considered. Inset, Mean ± 25^th^-75^th^ distribution of resampled means of average number of PFs per animal. f) Directionality index (DI, see Methods) for all neurons identified in the two experimental groups. Inset, mean ± 25^th^-75^th^ distribution of resampled means of average DI per animal. g) Remapping of place cells was minimal. The plot displays the x (North-South axis) and y (East- West axis) difference of the PF center position between session S28 and S34 for place cells in animals trained in the allocentric (red) or egocentric (blue) task.

### Analysis of neural Ca^2+^ activity in startboxes shows differences in in non-local representations between allocentric-trained and egocentric-trained rats

We then turned our focus to the neural activity occurring in the startbox prior to entry into the arena. Due to the task configurations, animals were expected to plan and decide their destination in the startboxes, and could only start a trial after the automatic opening of the door. The start of a trial was defined as the moment when animals started moving out of the startbox. We considered the 10 s preceding this time point and mapped the information content of the cells activated in this time interval (i.e. determined where these cells were active when the animal was in the arena).

We found that representations in the startbox differed between animals using an allocentric versus an egocentric strategy (Fig. 3). In both egocentric- and allocentric-trained animals, a significant fraction of this activity mapped onto one or more sandwells, with a higher fraction doing so in allocentric-trained animals (Figure 3a; Mixed Linear Model Regression ego/allo: z= -3.925 P<0.001). Importantly, we found that the majority of this activity in allocentric-trained animals represented a single sandwell, whereas they would often map to two, or more, sandwells in egocentric-trained animals (Fig 3b,c and Supplementary Figure S14,15; Mixed Linear Model Regression ego/allo: z= -15.487 P<0.001. We have interpreted nonlocal representations such as the ones observed here as instances of planning (Gobbo *et al*., 2022), and others have underlined a potential role of the hippocampus in playing out possible outcomes and destinations (Comrie *et al*., 2022). Consistent with this provisional hypothesis, we found that these measures were similar in “learning” (sample) and “recall” trials (choice); and in trials in which they would proceed to make a correct or incorrect choice (i.e. if the animal navigated directly to the correct sandwell, or instead dug first at one or more incorrect sandwells) (Supplementary Fig S14).

**Figure 3:**
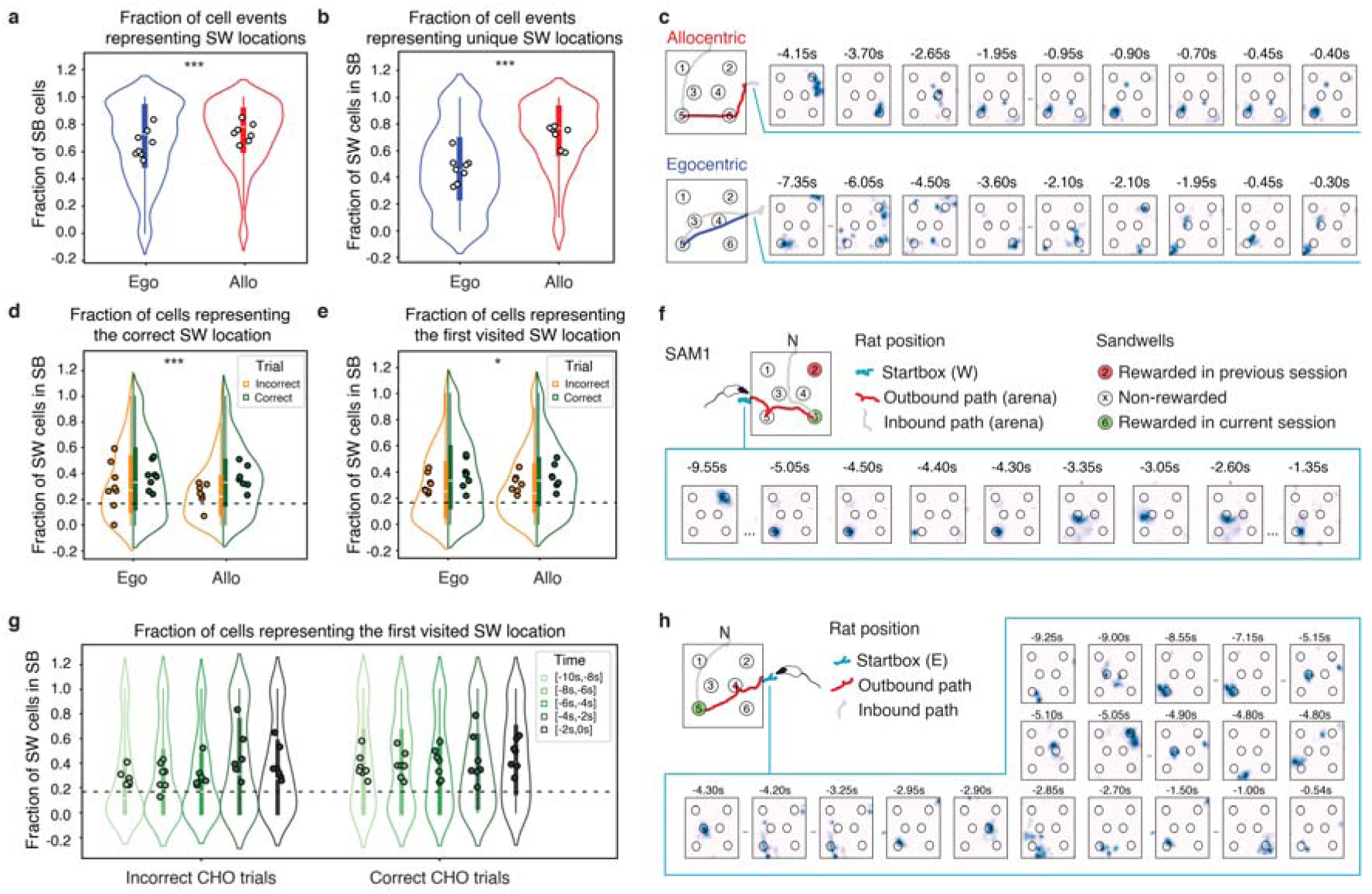
Reactivation of cells in the startbox. a) Fraction of all cells active in the startbox in the 10s preceding the trial start that represent one or more sandwells (SWs). b) Fraction of cells in (a) that represent a unique sandwell. c) Examples of nonlocal activity of cells in the startbox for the same rat trained in the allocentric protocol (above, red trace: Session A31) and in the egocentric one (below, blue trace, Session E32) with matching rewarded sandwells. On the left panel, the position of the rat in the trial is displayed in grey. The outbound path is highlighted in color (red or blue). The spatial fields of the cells active in the startbox in the 10s preceding the beginning of the trial are displayed as occupancy maps; the list is shortened for space purposes, and the corresponding events map and full list of cells are displayed in Supplementary Figure S15. Timestamps above cell maps indicate the time of cell activity from the beginning of the trial; negative times indicate events that happened before the trial start. When animals were trained in the allocentric protocol, most cells identified in the startbox mapped to a single position in space whereas in the egocentric case they would often map to multiple locations in space. d) Fraction of cell activations that correspond to the rewarded SW in allocentric- or egocentric-trained animals divided according if the animals would then navigate directly to the correct SW (correct trials) or would dig at other SWs first (incorrect trials). e) Fraction of cell activations that correspond to the first actually visited SW during the trial f) Example of startbox activity in the first learning trial (SAM1) where no prior information regarding the identity of the rewarded well is available to the rat. g) Fraction of activity events corresponding to the first visited SW in allocentric-trained animals calculated in intervals of 2s before the start of the trial, divided by correct and incorrect trials. h) An example of spatial fields of active cells before trial initiation. Timestamps indicate the time in seconds before the rat left the startbox and started the trial. Throughout the figure, violin plots with quartiles (boxplots) represent the distribution of individual trials, and large dots are the mean of individual animals. Egocentric data n=628 trials from 52 sessions/8 animals. Allocentric data n=590 trials from 46 sessions/7 animals.

Our analysis was confirmed by a Multinomial Logistic Regression (MLR) classifier trained on the neural activity in the Exploration stage and applied to the decision time window in the startbox preceding trials (see Methods). The decoder predicted a significant overrepresentation of sandwells compared to the expected rate (Supplementary Figure S16a,b), whereas the non-sandwell locations were significantly underrepresented in the startbox activity (Supplementary Figure S16c,d). Consistently with our analysis, multiple alternatives were decoded (Supplementary Figure S16d-f).

### Neuronal activity in allocentric-trained animals is linked to future behavior

We next sought to understand if the activation of neurons representing various locations in the arena was biased towards future destinations. We calculated how many of the instances of cells active in the startbox representing possible destinations mapped the correct sandwell (i.e. the rewarded one) – a location that, in this experiment, could be up to 1.5 m away from the animal position. Cells representing the correct destination were significantly more likely to be active in the 10 s preceding correct compared to incorrect trials (Fig. 3d, Mixed Linear Model Regression ego/allo z= 1.799 P=0.072, correct/incorrect z=5.267 P<0.001, ego/allo * correct/incorrect z=-1.519 P=0.129). We obtained similar results classifying trials by user annotation or with an automatic classification with an algorithm, confirming our trial classification. (Supplementary Figure S17a,b). Similarly, by automatically determining the cell’s field as the position of its maximum event rate in the arena, we showed that, in correct trials, a significantly higher fraction of cells active in the startbox mapped to a position along the trajectory leading to, or the destination of the following trial in the allocentric group than egocentric one (Supplementary Figure S17c). Conversely, the percentage of active cells representing the first visited sandwell was above chance on both correct and incorrect trials (Figure 3e and Supplementary Figures S18-20; Mixed Linear Model Regression ego/allo z= -0.088 P=0.375, correct/incorrect z=2.57 P=0.01, ego/allo * correct/incorrect z=1.037 P=0.3). Indeed, we found similar levels of activity even in the 10 s preceding the very first learning trial in which, by task design, there is no past knowledge about the reward status of the sandwells (Fig 3f and Supplementary Figure S21). While the representation of the correct sandwell was essentially at chance (as the animals had no prior knowledge of which sandwell would contain the food reward), a marked over-representation was apparent of the first sandwell the rat would visit, particularly in allocentric-trained animals (Supplementary Figure S22).

We then zeroed in on specific aspects of the startbox activity. Having shown that multiple possible destinations can be represented, we first asked if the percentage of cells mapping to the correct sandwell location increased as the start of a trial approached, i.e. if there was a “decision point” related to the start of the trial. We did not observe such an effect for the correct sandwell, possibly because, on incorrect trials, the frequency of cells representing the correct sandwell tended to decrease among cells active in the startbox as the trial start-point approached (Mixed Linear Model Regression correct/incorrect z=4.883 P<0.001, time z=-1.517 P=0.129, time*correct/incorrect z=1.917 P=0.55; Supplementary Figure S20a). However, when considering the first visited SW, we found a significant effect of time in allocentric-trained rats (Mixed Linear Model Regression correct/incorrect z=1.305 P=0.192, time z=2.588 P=0.011, time*correct/incorrect z=-0.932 P=0.351; Supplementary Figure S22b), particularly on choice trials (Mixed Linear Model Regression correct/incorrect z=0.493 P=0.622, time z=3.159 P=0.002, time*correct/incorrect z=-1.257 P=0.205; Fig 3g,h). No such effect was seen in egocentric-trained animals (Supplementary Figure S22c,d).

To corroborate our results, we trained a binary naive Bayes decoder on the rats’ spatial position on that day’s Exploration stage and then applied it to the 10s period prior to trial start on sample and choice stages. We then measured the distance between the predicted location and the goal location (see Methods). When we compared the goal location prediction errors in the startbox between trials from the allocentric and egocentric groups, we found that, for allocentric-trained animals, errors were significantly lower than for egocentric trials, consistently with our interpretation of representations being less precise in the egocentric group (Supplementary Figure S23,24; one-tailed Mann-Whitney U test). Furthermore, the median distance to the correct sandwell was lower in correct versus incorrect trials in the allocentric group, but not in the egocentric one (Supplementary Figure S23,24).

We measured the average event rate of cells mapping to sandwells, or other portions of the arena, during the Exploration stage and confirmed that our analysis was not biased by an intrinsic differential firing rate of cells corresponding to a sandwell or another location in the arena, to the rewarded sandwell, or the first visited sandwell compared to the others (Supplementary Figure S25a-e). Similarly, we found no evidence that the rewarded sandwell was represented by a higher number of cells with fields in that location (Supplementary Figure S25f). Inspection of neuronal Ca^2+^ traces confirmed that the events detected Startbox are due to reliable cell activity mapping to future destinations (cells mapping to visited or rewarded well) or alternative possibilities in recordings from animals performing in allocentric and egocentric tasks (Supplementary Figures S26 and S27 respectively). Last, we calculated the dispersion and inter-event time between neuron events in the startbox; we found that these occurred with higher temporal clustering than what would be observed by chance. Although one-photon Ca^2+^ imaging only allows to determine the activation of overlapping populations by flattening the temporal time scale – and therefore precludes the observation of fast, structured patterns and sequences with millisecond precision – this nevertheless suggests that startbox events display some degree of temporal clustering. This was true for all cells active in the startbox, as well as cells mapping to the rewarded well in the session, or the first visited sandwell (Supplementary Figure S28,29). Overall, our data suggest an increased focus on the chosen destination when animals identify their goal in allocentric terms.

We then asked whether the same cells representing the possible as well as the soon-to-be-chosen destination would be active across different trials in allocentric animals starting from East, South or West startboxes (and possibly from the North Goal Box). In allocentric trials, the rewarded sandwell represents a common destination, but the path to be taken differs considerably in direction and length.

Indeed, we found that more than half of the cells active in the startbox were reactivated in the 10 s before the trial start across trials in the same session. Perhaps unsurprisingly, we observed a trend whereby the percentage of common cells is higher if animals started from the same startbox. Overall, this consistency was greater in allocentric- than egocentric-trained animals (Figure 4a; Mixed Linear Model Regression Fraction common cells ∼ Group + (1|animal) Allo same SB vs Ego same SB z=-0.25 P=0.022; Allo different SB vs Allo same startbox (SB) z=0.63 P=0.139). The same trend was observed across sessions when comparing trials with the same rewarded well (Figure 4b; Mixed Linear Model Regression Fraction common cells ∼ Group +(1|animal) Allo same SB/same RW vs Ego same SB/same RW z=-12.53 P<0.001; Allo same SB/same RW vs Allo same SB/different RW z=- 7.12 P<0.001). Notably, the trials starting from North (the Goal Box) displayed a lower number of common cells, perhaps because even trials represent a “return” trial, whereby path integration can ease the hippocampal workload (Supplementary Figure S30). Importantly, a remarkable number of neurons active in the startbox represented the same destination regardless of the starting point (Fig 4c,d and Supplementary Figures S31,32).

**Figure 4:**
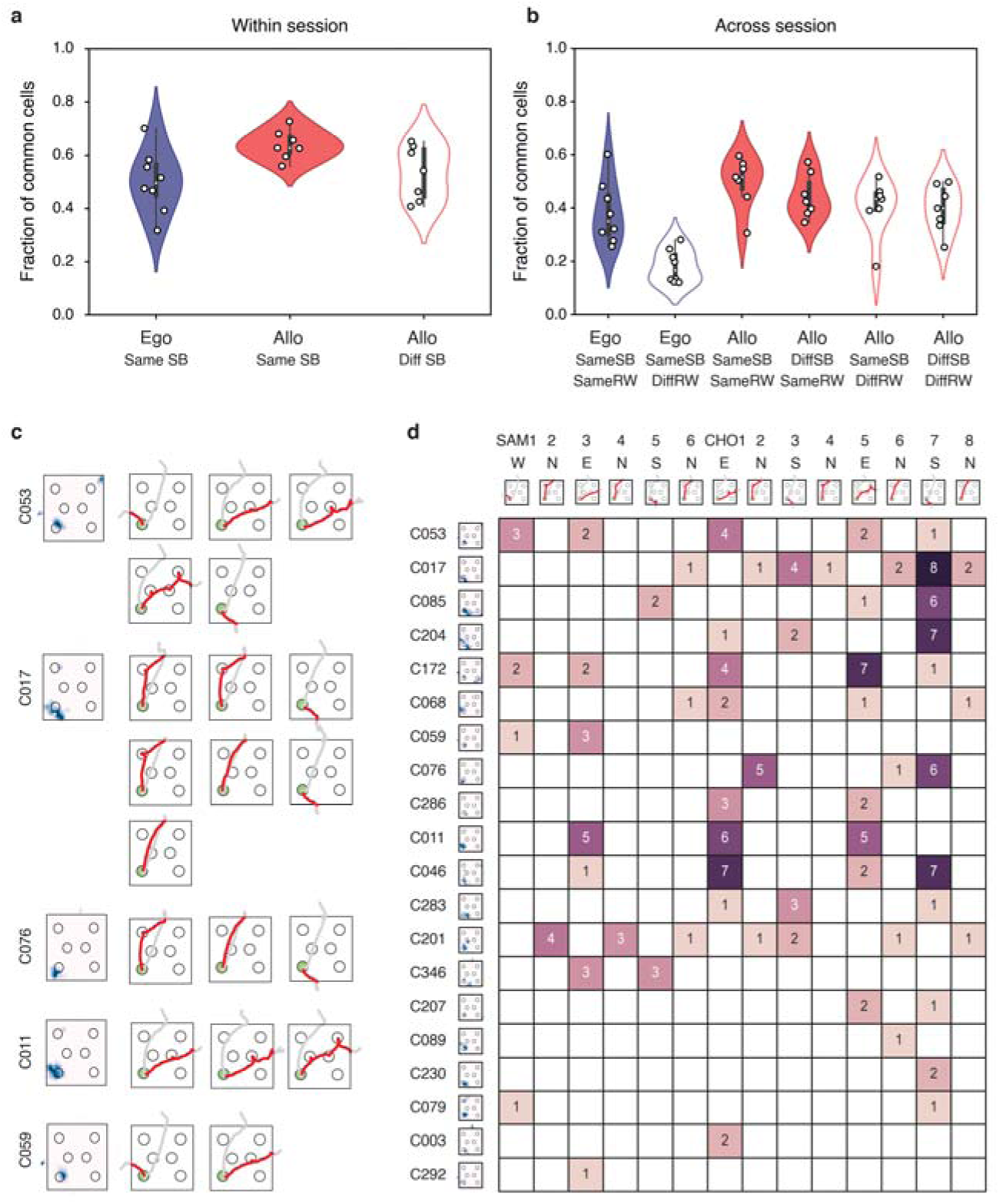
Invariance of destination representing cells from origin. a) Fraction of cells active in the startbox for trials in the same session for egocentric- or allocentric-trained animals. b) Fraction of common cells active in the startbox between sessions in the startbox pre-trial interval (SB). c) Examples of cells representing the current rewarded well (SW) and animal destination active while rats are in different startboxes. On the left, the rate map of the cell showing the field as event rate while the animal is in the arena. On the right, the path taken by the animal after that cell was active in the startbox. d) Summary of cells mapping onto the rat’s destination in the various trials of the session (same session as in c). Rows indicate the cells representing the future destination in various trials for a whole session. Next to them, their place fields are represented as rate maps. Columns represent the session trials (sample and choice trials) along with the path taken in that trial after exiting the startbox. The double entry matrix shows if and how many times each cell was active in the 10 s preceding the beginning of each trial while the animal was in the startbox. Numbers indicate the number of times each cell was active in the startbox during the 10s considered.

Together, these results indicate that representations by egocentric-trained animals are less precise, more ambiguous and less anchored to decision making compared to when rats are using allocentric coordinates. Consistent with this, in comparing the neural dynamics during trial execution (i.e. the time series of Ca^2+^ activity as the animals navigated between the starting box and the rewarded well), we found that allocentric trajectories had higher canonical correlations (i.e. higher trial to trial correlation of the neural dynamics after finding the linear transformation between trials that maximized correlation) than did egocentric trajectories (Supplementary Figure S33, see Methods). We chose CCA to enable the comparison between datasets with different, or partially overlapping, neuronal populations. In general, two trajectories are expected to have maximum CCA correlation if the profile of the neural vectors representing them overlaps completely (see also Supplementary Discussion). This was not due to spatial properties of the trajectories such as pairwise trajectory distance (Mixed Linear Model Regression z=-1.126 P=0.26), spatial correlation (Mixed Linear Model Regression z=0.736 P=0.561) or percentage of active neurons (Mixed Linear Model Regression z=0.581 P=0.462) (Supplementary Figure S33g-i). Rather, we hypothesized that this was due to a stronger structural similarity between the neural representation of individual allocentric trajectories in the neural space. Consistent with this interpretation, we also found higher CCA correlation between reference and symmetrical trajectories (i.e. identical upon 180° rotation of the arena; see Methods) in allocentric-trained animals, and a CCA decoder could predict the animal location along those trajectories better than chance, whereas a neural decoder did not. This was expected considering that symmetrical trajectories occupy distinct spatial positions, and recruit separate neuron populations (Supplementary Figure S33f,l).

### Neuronal activity during decision making is necessary for allocentric- but not egocentric task execution

The causal significance of these ostensibly predictive facets of hippocampal neural activity was explored using optogenetic inactivation of dorsal CA1 activity.

Inactivation was confined either to the startbox, or – in separate sessions – extended to include the period when the animals were navigating in the arena. To inhibit neural activity, we virally expressed the JAWS-GFP inhibitory opsin in CA1 pyramidal neurons (Chuong *et al*., 2014), aiming to encompass as much of the dorsal hippocampus bilaterally as possible (Supplementary Fig. S34). Using acute silicon probe recordings in vivo, we demonstrated robust and prolonged inhibition of cell-firing with red light illumination (Fig. 5a,b). Of a sample (n=48) of putative excitatory cells, the majority were inhibited (Fig 5b, Wilcoxon Signed Ranks Test, Z = 4.71 n = 48 cells, P < 0.0001), and for the full duration of the light-evoked inhibition (Fig. 5c). A further 20 were not significantly modulated, but still showed a trend towards inhibition (Supplementary Figure S35a). The occurrence of sharp-wave ripples (SWRs) was also inhibited (Fig. 5d,e; Wilcoxon Signed Rank Test,Z = 2.03, n = 5 recordings from 5 animals, P < 0.05). AAVs expressing either JAWS-GFP, or GFP as control, were then injected into dorsal CA1 in a new group of animals that would go on to be trained in the behavioral task with chronically implanted bilateral light cannulae directed at the same dorsal hippocampal region (Fig 5f and Supplementary Figure S36). Performance during training did not differ from that seen earlier during acquisition and was at asymptote with respect to choice performance, probe test accuracy, and the selective impact of intra- and extra-arena cue occlusion in allocentric-trained animals (Fig 5g and Supplementary Figure S37; 3-way repeated measures ANOVAs: Allocentric: Cues/Curtains F(1,22)=45.59, P<0.001, Virus F(1,22)=1.53 P=0.23, Session, F(1,22)=0.10 P=0.76; Egocentric: Cues/Curtains F(1,12)=2.94, P=0.11, Virus F(1,12)=3.05 P=0.11).

**Figure 5:**
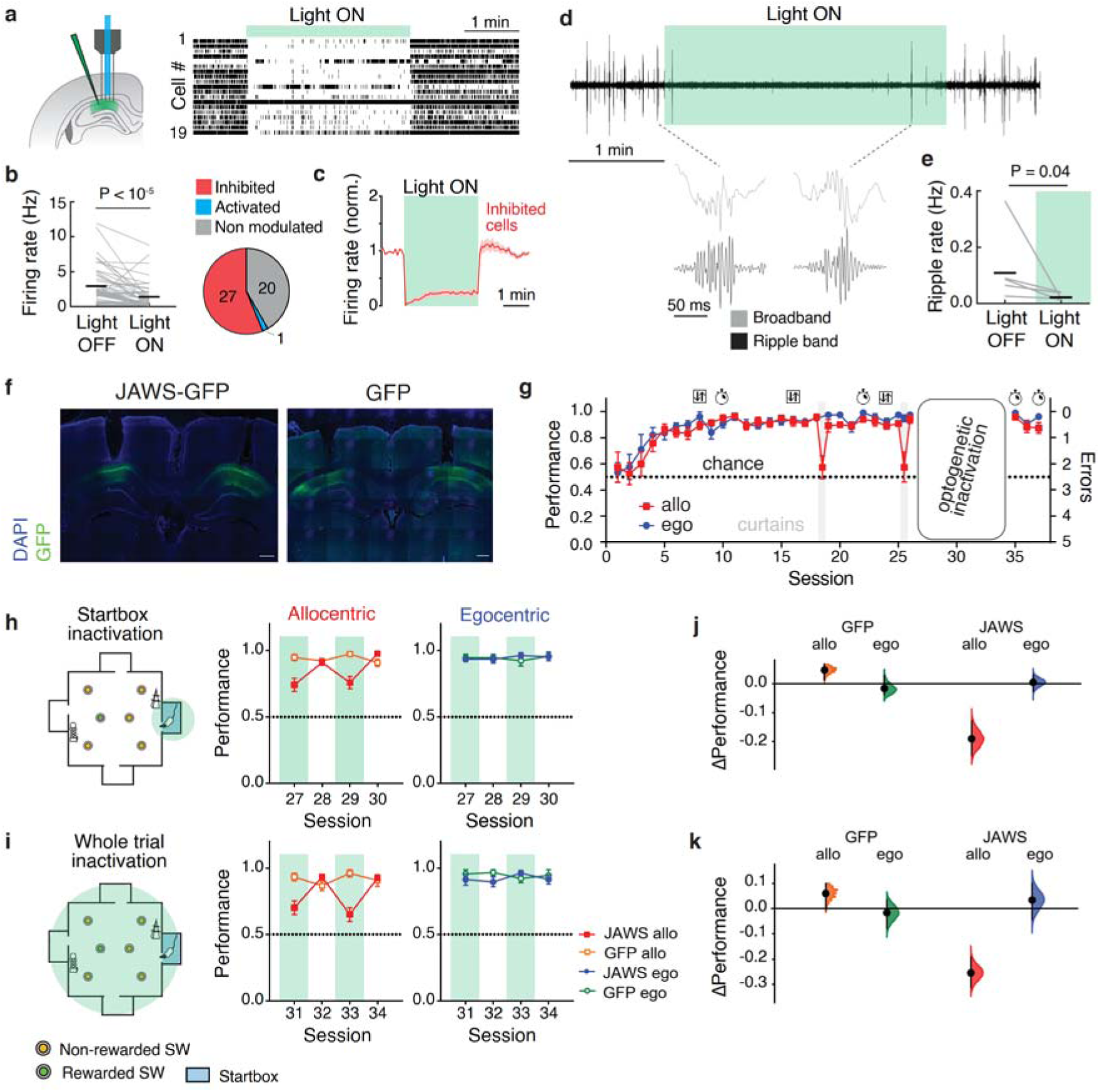
Optogenetic inhibition of CA1 cell activity in the startbox impairs allocentric navigation. a) Experimental setup. A light guide was directed at the dorsal CA1 region of hippocampus with concomitant silicon probe recording. Impact of 3 min light stimulation (light-ON) on 19 simultaneously recorded putative excitatory cells, showing reduced firing during light-ON the majority of cells. b) Left, mean firing rate of putative excitatory cells decreases during light-ON periods, n=5 rats. Right, over half of recorded putative excitatory cells are significantly inhibited during light-ON with a minority of cells not significantly affected. c) Firing rate changes of significantly inhibited cells before, during and after the light pulse, showing a near-complete inhibition of firing and a modest and short-lasting rebound effect. d) Example of a ripple-band filtered recording showing a decrease in occurrence of sharp wave-ripple (SWR) events. e) Occurrence of SWRs declined during the light-ON period in all 5 recorded animals. f) Examples of coronal sections of rats expressing JAWS-GFP or GFP showing transgene expression in CA1 and position of optic fiber cannulas. Scale bar 500 μm. g) The behavioral performance of this cohort of animals was broadly identical to what is shown in the Ca2+ imaging animals (see Figure 1). h,i) Startbox inactivation of cell activity (light green) caused a significant decrease in sandwell choice accuracy in the event arena (sessions 27-30) restricted to allocentric-trained animals. The decrease in performance was not observed in GFP virus controls and in egocentric-trained animals. j,k) Whole trial inactivation of cell firing caused a slightly larger decrease in choice performance, restricted to allocentric-trained animals and absent in GFP virus controls. Data points are mean±sem. (i,k) show effect size calculated with DABEST (Ho *et al*., 2019).

The key question was whether optogenetic inhibition of startbox activity would inhibit choice performance regardless of the strategy, or in a strategy-dependent manner. We therefore inhibited the dorsal CA1 while rats were in the startbox before the start of the trial (see Methods). The striking finding was that the inhibition of neural activity during this time frame significantly reduced choice performance in allocentric-trained animals without effect on egocentric-trained animals (Fig. 5h). Control sessions where no light was delivered show no such impairment, and the effect was specific to JAWS, but not the GFP group (repeated measures 3-way ANOVA Light x Virus x Behavioral protocol F(1,22)=32.96 P<0.001, post-hoc Šídák’s multiple comparisons test Light-ON:Allo:JAWS vs. Light-OFF:Allo:JAWS P<0.001) with the impact being on allocentric- but not egocentric-trained animals (Šídák’s multiple comparisons test Light-ON:Ego:JAWS vs. Light-OFF:Ego:JAWS P=0.99)). Control animals expressing GFP in dorsal CA1 showed no changes in behavior in the Light-ON vs Light-OFF conditions (Šídák’s multiple comparisons test, Light-ON:Allo:GFP vs. Light-OFF:Allo:GFP P= 0.62; Light-ON:Ego:GFP vs. Light-OFF:Ego:GFP P=0.99).

Quantification of the relative changes in performance resulting from startbox or whole arena inhibition is shown in Figs. 5j,k). Optogenetic inhibition throughout the event arena showed a trend towards a further lowering of choice accuracy in the allocentric-trained animals, but it was not significantly lower than that observed with only startbox inhibition (Fig. 5i; Mixed effect model, Tukey’s multiple comparisons test JAWS:startbox:Light-ON vs test JAWS:arena:Light-ON P=0.12). We also found that neuron silencing with JAWS increased the time spent between the door opening and the start of trial (“Decision time”) and the latency to reach the correct sandwell – particularly in sessions 31 and 33 – in Light-ON compared to Light-OFF sessions; this effect was only displayed by animals in the allocentric group, while illumination had no effect on the egocentric group (Supplementary Figure S38). Quantification of digging behavior during probe tests complemented the results seen in choice performance as it showed a failure to dig at the appropriate correct sandwell in the arena in Light-ON probe trials (Supplementary Figure S39).

## Discussion

The two key findings of this study are that (i) in startboxes at the edge of a training arena, hippocampal representations of remote locations, as captured in Ca^2+^ imaging, are different in allocentric-trained animals than in those egocentric-trained, and (ii) inhibition of this neural activity in the hippocampus can cause measurable and significant errors in the navigational choice performance of allocentric-trained animals. Our results show that the hippocampus plays a causal role in allocentric but not egocentric representations of spatial recency memory. Furthermore, we demonstrate that inhibiting hippocampal activity during decision-making prior to navigation can impact animals’ performance, but only when they need to make use of these allocentric spatial representations.

### Nature and role of nonlocal neural activity in the startbox

Here and in previous work (Gobbo *et al*., 2022) we observed the activity of neurons representing possible goal locations while rats were located in the startbox before trials start. Correct sandwell destinations were preferentially represented, particularly in trials where rats identified the rewarded sandwell as their first choice. Nevertheless, neurons mapping to other locations were also active (Figure 3a-e). Furthermore, nonlocal neuron activity in the startbox was also recorded without prior knowledge of the rewarded sandwell in the session in the first Sample trial, often correlating with the first inspected sandwell by the rat (Figure 3f). It is therefore tempting to interpret this activity as self-generated activity representing possible destinations. In our experiment, rats were well acquainted with the arena environment as well as the task structure. This is expected to have equipped the rats with a stable representation of the various arena locations, including salient ones like the sandwells (Ziv *et al*., 2013), as also shown by the stability of place fields across sessions (Figure 2).

Hence, it can be speculated that rats are using their existing knowledge to weigh out possible destinations. Without necessarily advocating for conscious activity, we think these are consistent a proposed role of the hippocampus in imagination and planning (Comrie *et al*., 2022). For instance, generative activity of the hippocampus at both single cell level and population level has been observed (Kay *et al*., 2020), and flexible route representations have been reported when rats have to navigate complex environments with barriers (Widloski & Foster, 2022). In human studies, extensive evidence has revealed a role for the hippocampus in imagination (Hassabis *et al*., 2007).

Here, we have adopted the terms “nonlocal activity” or “activation” to refer to neurons displaying calcium events in the startbox. One important caveat is that technical and analytical reasons, such as the lower temporal resolution of calcium imaging – which precludes some analytical methodologies (Tingley & Peyrache, 2020; van der Meer *et al*., 2020) – and the single neuron approach taken here, prevent us from making direct comparisons to experiments with electrophysiological recordings of replay and other forms of reactivations. This also prevents us from making considerations about our data in relation to other phenomena implicated in memory access such as organization around phases of theta frequencies and SWRs, or assembly patterns where cell ensembles display task-related co-activity (Dupret *et al*., 2010; Sugden *et al*., 2020; Gava *et al*., 2024).

Several reports have demonstrated the reactivation of remote goals and trajectories at decision making points (Gupta *et al*., 2010; Pfeiffer & Foster, 2013; Gillespie *et al*., 2021). Among these, some of these reactivations can occur in the form of replay, i.e. the rapid, sequential activations of cells in the order in which their place fields were previously traversed (Genzel *et al*., 2020). While still debated, growing evidence shows that neuron sequences representing known goals, trajectories and destinations can be replayed on a millisecond time scale, supporting decision making (Pfeiffer & Foster, 2013; Ólafsdóttir *et al*., 2015, 2018; Pfeiffer, 2017; Xu *et al*., 2019; Widloski & Foster, 2022). Increasing evidence also suggests that the temporal position of spiking with respect to theta phase is important for encoding current goals and future behavior (Wikenheiser & Redish, 2015; Wang, Foster, *et al*., 2020). Due to aforementioned limitations of our experimental and analytical approach, our results demonstrating the activity of individual place cells encoding nonlocal arena locations in the startbox cannot be extended to experiments addressing replay or phase-locked activity of cell assemblies. Future work will determine if, and to what extent, our observations relate to these reports.

### Spatial strategy shapes hippocampal representation

Despite its limitations in terms of temporal resolution, miniscope Ca^2+^ imaging allowed us to follow the dynamic of neurons across days and observe how they would be influenced by the spatial strategy. When comparing allocentric and egocentric groups, we observed that the quality and the precision of space representations are markedly affected by the strategy employed. In rats performing in the allocentric task, neurons representing individual destinations as well as intermediate locations are active in the startbox. While we observed such instances also in egocentric-trained animals, variability was higher, and a significant number of cells were active at multiple locations of the arena or had no obvious spatial correlation. In some instances, those could map broad features of the encoded sandwells (e.g. left side of the arena). One could speculate that they represent a less attentive representation of space: for instance, symmetrical sandwells near opposite corners were often co-represented, which might suggest some sort of perceived equivalence. Similarly, Moore *et al*. (2021) reported a deterioration of spatial accuracy when idiothetic cues prevail over allocentric ones. In either way, our data demonstrate that faithful reactivation is not necessary to solve egocentric planning.

Recent electrophysiology data have shown that deep and superficial layers in CA1 can project differentially to prefrontal and medial entorhinal cortical areas, and play a different role in the task by encoding future trajectories and choices (superficial CA1), or learned reward locations (deep CA1) (Harvey *et al*., 2023; Esparza *et al*., 2025). Ca^2+^ imaging data have shown that deep layer pyramidal cells are preferentially tuned to local cues, while superficial ones are oriented to global environment (Esparza *et al*., 2025). Furthermore, deep neurons were found to be modulated by goal-oriented learning to a larger extent than superficial ones (Danielson *et al*., 2016). While our recordings are likely to sample from a combination of cells across the layers (Supplementary Figure S4), this could potentially complicate our evaluation of the spatial properties in our tasks, since the two populations could differ in the processing of conflicting egocentric and allocentric cues (Sharif *et al*., 2021).

### Necessity of hippocampal activity in allocentric task

A crucial problem in linking neural activity with its scope is demonstrating if experimentally interfering with it has any effect on behavioral performance. This may also change depending on the brain area under observation. For instance, structured sequences to a salient location have been recorded from hippocampal neurons during homing behavior in a two-dimensional arena, yet the pharmacological inactivation of the hippocampus does not obviously affect the animal’s behavior (Pfeiffer & Foster, 2013; Duszkiewicz *et al*., 2023). Recent experiments showed that neuronal activity structured in replay sequences was necessary for the performance in a flexible navigation version of the cheeseboard maze; however the performance in a more elementary contextual association was not affected by a perturbation that eliminated theta sequences and temporal structure of replay, but preserved neuron activation (Liu *et al*., 2023).

Here, we found that suppressing hippocampal activity during decision making impairs the performance of rats in the allocentric version of the everyday memory task. The task, arguably more complex than the egocentric one, is dependent on intact hippocampal activity (Broadbent *et al*., 2020; Kanakis *et al*., 2025). This is consistent with existing literature showing that the use of allocentric coordinates relies on an active hippocampus, whereas using an egocentric framework does not (Packard & McGaugh, 1996). One possibility is that egocentric planning relies on striatal activity, as suggested by Packard and McGaugh (1996). Some level of representation may still take place at the hippocampus level (as also our data indicate) without being necessary to solve the task. Interestingly, in Packard & McGaugh (1996) the pharmacological inactivation of the striatum did not result in chance performance, but rather in the recovery of the – allocentric – place response, suggesting the removal of a striatal interference on the hippocampal processing. Although in our experiments allocentric-trained animals did not fall to chance performance levels, this could be due to several factors, including re-orientation in the arena and residual activity of uninhibited cells in the dorsal or, possibly, in the intermediate and ventral hippocampus (Jarzebowski *et al*., 2022).

The possibility of interfering with hippocampal activity was previously inquired in the context of alternating W-maze by suppressing SWRs. In this maze, the authors showed a selective impairment of outbound trials (from the central arm to the external arm not visited immediately before), but not of inbound ones (towards the central arm) (Jadhav *et al*., 2012); although a direct comparison is difficult, a re-interpretation of the data in light of our results suggests that rats in their outbound trials were using, at least in part, allocentric cues beyond working memory. In support of this interpretation, in early sessions the solution of the T-maze in Packard and McGaugh (1996) is also based on allocentric coordinates.

Different types of activity patterns will likely be affected in our experiments beyond the observed silencing of hippocampal neurons such as SWRs (Figure 5d) and theta oscillations. Hence, the optogenetic inactivation achieved in this experiment is to be regarded as a state-dependent perturbation of hippocampal activity during the planning window. From a technical standpoint, the choice of continuous illumination in our optogenetic inactivation experiments ensured that neural activity was suppressed regardless of the presence of SWRs. For instance, previous work has shown that reactivation events can occur both within and outside of SWRs (Widloski & Foster, 2022). Future work will aim to address their role individually with targeted silencing.

How performing a simpler task with less challenging requirements reduces the quality of spatial information remains to be determined. Habit learning seems to involve the suppression of hippocampal information processing by other brain areas (Packard & McGaugh, 1996), which could interfere with hippocampal activity. Otherwise, the use of an egocentric system (Wang *et al*., 2018) or local cues (Collett *et al*., 1986) could deteriorate allocentric fields. An open question is then whether information content in egocentric-trained animals simply reflects poorer space selectivity or the concomitant representations of alternative information. Last, if the hippocampus is involved in planning, at least in a fraction of the tasks, by playing out possible scenarios, what drives the decision point? Is it a “Eureka!” moment due to coincident confirmation from memory? Or, vice versa, is it due the accumulation of enough representations of a given destination resulting in a more secular “Oh well, that’ll do!” moment? Future research will shed light onto these fascinating aspects.

## Material and Methods

### Animal subjects

Male Lister-Hooded rats (Crl:LIS 603) were 2-3 months old at the start of the experiments. The animals were purchased from Charles River UK and group housed until surgery; after surgery, animals were housed in single cages. During the behavioral task, they were food-deprived to 85–90% of the free-feeding weight against a growth curve, with free access to water, 12:12 h light-dark cycle with training in the light phase. Care of the animals complied with the UK Animals (Scientific Procedures) Act conducted under a Project Licence (PPL P7AA53C3F).

### Immunohistochemistry

At the end of the experiments, the animals were anaesthetized with 200mg/ml pentobarbital and perfused transcardially with cold PBS (Phosphate Buffer Saline, P4417 Sigma Aldrich) and 4% formaldehyde in PBS (Sigma Aldrich 441244). Heads were postfixed for 24 h in formaldehyde, and then the brains extracted and postfixed overnight. These were transferred to 20% sucrose PBS and cut with a cryostat (Bright Instruments). After washing in PBS, slices were permeabilized in PBS 10% NDS (Normal donkey serum, Sigma Aldrich D9663) 0.1% Triton X-100 (Sigma Aldrich T8787) for 30 min, and then incubated in PBS 10% NDS 0.1% Triton X-100 with mouse Anti-GAD67 Antibody (Sigma Aldrich MAB5406 clone 1G10.2) 1:1000, guinea pig Anti-NeuN (SYSY 266 004) 1:500 overnight at room temperature. The next day, three washes with PBS 0.1% Triton X-100 were performed, and then incubated in PBS 0.1% Triton with 1:200 donkey anti-mouse Alexa 647 (Thermo Fisher A-31571) and 1:200 anti-guinea pig Alexa 555 (Thermo Fisher A-21435). After three washes in PBS, the slices were mounted in Fluoroshield with DAPI (Sigma Aldrich F6057).

### Microscopy

Multichannel images were acquired using a Nikon A1R Ti:E inverted confocal microscope with 1AU pinhole dimension, using a 40X Plan Apo/NA 1.25 OIL objective (histological staining) or 10X Plan Flour/ NA 0.3 (GCaMP6f/DAPI acquisition). Sequential acquisition of the four channels was performed with the following laser lines: DAPI 402nm, GCaMP6f 488nm, NeuN/Alexa555 562nm, GAD67/Alexa647 639nm, and 450/50, 525/50 and 595/50 filter cubes. Histological sections stained with DAPI were images with a Leica TCS SP8 on DM6B-Z-CS confocal microscope using objective HC PLAPO CS2 20x/0.75 DRY and laser lines DAPI 405nm, GCaMP6f 488nm. Serial histological analysis was performed with Zeiss Axio Imager Z2 and HXP 120 C Fluorescence Light Source using objective Plan-Apochromat 5x/0.16. Images were acquired with CoolSNAPHQ2 camera (Photometrics); acquisition parameters were maintained constant across animals.

### Surgical procedures

Surgical procedures were performed under sterile conditions and according to best practice as described in (Gobbo *et al*., 2022). Animals were maintained under 3-2% isoflurane (Covertus absis01) reversible anaesthesia and 0.5ml rimadyl (Zoetis) administered as an analgesic at the beginning of each surgery. In the first surgery, 1µl of AAV1.CaMKII.GCaMP6f.WPRE.SV40 (UPenn Vector Core, then Addgene 100834-AAV1) diluted with sterile PBS (Sigma-Aldrich) to a final concentration of 5.7 10^12^ vg/ml was injected in the CA1 region of each hemisphere (stereotaxic coordinates from bregma AP 3.6, ML ±2.2, DV 2.2 from dura) at 100nl/min using an automated injection pump and a Hamilton syringe equipped with a Nanofil needle (World Precision Instruments). The GRIN lens was implanted 7-10 days after the first surgery. In the second surgery, three surgical screws were implanted in the skull (Screws and More, Ennepetal (Germany), DIN 84 A2 M1X3). A circular craniotomy was performed with a trephine for microdrill (F.S.T. 18004-18) at AP 3.8, ML 2.4. A cylindrical volume of cortical tissue was then aspirated manually with a sterile blunt needle in the 27G-30G range. Constant irrigation with cold saline (Baxter) is performed during aspiration to prevent swelling and clean blood. A small portion of alveolar (anteroposterior) and callosal (coronal) fibers were then removed with the same blunt needle with mild aspiration to expose the outer surface of the hippocampus. The aspiration cavity was irrigated with cold saline and soaked sterile gelatine sponge (Gelita-Spon GS-110, Delta Surgical) until complete stop of the bleeding. A GRIN lens (1 mm-wide, 9-mm long, Inscopix) was lowered with a 5° angle in position AP 3.8, ML 2.4, DV 1.9 from dura (the dura is measured before the aspiration). Two stainless steel rods (0.09 mm diameter, CrazyWire UK) are attached to the side of the GRIN lens with superglue before implantation. A thin layer of surgical silicone (KWIK-SIL, World Precision Instruments) was applied to the sides of GRIN lens to prevent cement from entering contact with the brain tissue. The GRIN lens was cemented in place with Super Bond dental cement (PRESTIGE DENTAL, Bradford UK) to cover the skull and including the skull screws. After cement curing, the skin was sutured with Vicryl (W9500T, Ethicon).

Surgeries for animals in the optogenetics groups were performed similarly. pAAV-hsyn-Jaws-KGC-GFP-ER2 (Addgene 65014-AAV5) or pAAV-hSyn-EGFP (Addgene 50465-AAV5) were diluted to 2.4 10^12^ vg/ml in sterile saline and injected bilaterally at AP 3.6 ML ±2.2 DV 2.2, and AP 4.1 ML ±2.6 DV 2.2 (500nl per site at 100nl/min), where DV coordinates are measured from dura. The injection needle was maintained in place for 10 min after the end of injection before withdrawing it slowly, a small quantity was then injected in the air and aspirated to exclude blocking. In the same surgery, fiber optic cannulas (400µm diameter, 0.50NA, length 5mm, outer diameter 2.5 mm, ceramic Thorlabs CFMC54L05) were implanted bilaterally at AP 3.6 ML ±2.6 DV 1.8 at a 5 degrees angle away from the midline. After insertion, the cannula was maintained in place for 5 minutes, then the exposed surface of the brain was covered with KWIK-SIL or KWIK-CAST (World Precision Instruments) before cementing with a thin layer of Super Bond dental cement (PRESTIGE DENTAL, Bradford UK) followed by dental cement. Three screws were implanted in the rats’ skull before measuring coordinates and inject viruses, as described above. Animals used in acute electrophysiology experiments were injected with pAAV-hsyn-Jaws-KGC-GFP-ER2 similarly with the exception that no chronic cannula implant was performed.

### Calcium recordings

The Inscopix nVista system was used to perform calcium imaging experiments. At least 3 weeks after the GRIN lens implantations, animals were temporarily anaesthetised and the field of view checked with the Inscopix miniature microscope (hereafter, miniscope). The baseplate support was scored and then cemented in place with SuperBond using the pre-existing layer of cement as described in (Gobbo *et al*., 2022).

During recording, the miniscope was positioned in the baseplate and secured with a miniature screw attached to the baseplate. The miniscope cable was connected to a commutator on the ceiling which enables full animal rotation. The commutator was connected to the nVista^TM^ imaging system (v 3.0) that controls the miniscope functioning and stores the recorded data. Elastic wires were used to connect the two extremities of the cable to hold the weight of the cable without weighting in the miniscope. Ca^2+^ recordings were collected at 20 frames/sec using the Inscopix nVista^TM^ system imaging system (v3.0) and synchronized with the camera behavior via an electronic signal to GPIO channels timestamping the time of two separate LED lights in the camera field of view at the beginning and end of the recording and between trials. The camera pixel size is 0.82 μm, yielding a field of view (FOV) of 870x656 μm (1061 x 800-pixels).

### Behavioral Apparatus

All experiments were conducted using a modified 160 cm x 160 cm square ‘event arena’. The walls are transparent (40 cm high) and the floor is composed of a total of 64 removable white or light blue tiles (20cm x 20cm; for regular cleaning purposes). 6 plexiglas sandwells (6 cm diameter, 4 cm depth) that contained the hidden reward pellets were placed in one or a subset of the panels with holes. The position of the sandwells is depicted in Figure 1. To ensure navigation to a remembered location rather than guidance by smell, any perceptible smell of the correct sandwell was masked in three ways. First, each sandwell had a spherical plastic bowl separating compartments for reward pellets that were either accessible or inaccessible to the animals (BioServ F0171) (0.5 g). These plastic bowls had holes ensuring that both the rewarded and non-rewarded sandwells contain the same number of reward pellets at approximately the same depth and thus should smell similarly. Second, the sandwells were also filled with clean mouse bedding or bird sand with their own odors (Pettex Aviary Cage Proud Bird Sand, The Pet Express). Third, sandwell use was randomized and spatially counterbalanced to minimize olfactory artefacts: (a) the sandwells used in the sample trials were not used for the choice trial of the same session; (b) all sandwells were used as rewarded or non-rewarded sandwells across days; (c) the arena floor was regularly wiped with a 70% alcohol-impregnated towel between sessions, and before recall and probe trials; (d) tile positions were randomized regularly by moving them around and/or exchanged with extra tiles between sessions or trials.

There are four startboxes located on the four sides of the arena named according to cardinal points (N, S, E and W). Two intramaze cues (a miniature telephone box and a purple box) were positioned inside the arena on the W and E walls. Three large and prominent 3-dimensional extra-arena cues (>30cm) were present, and additional visual cues were present on the walls of the procedure room. Heavy white cotton curtains were hung from ceiling rails that could be pulled all around the arena to mask extra-arena cues in sessions without cues (marked as “curtain” in Figures); intramaze cues were also removed. The calcium recordings and optogenetic experiments were conducted in two separate arenas in adjacent rooms; necessarily, they used different animals subject to different surgical procedures prior to behavioral training. The number and position of cues, and sandwells, were identical between the two arenas.

### Habituation

All animals were handled for at least a week before the start of the experiment. Rats were further familiarized with the experimenters after recovery from the surgery and during daily handling. They were habituated to dig for food in sandwells placed inside their home cages over a series of sessions. In a first behavioral habituation session, they were put in a Startbox and given a 0.5 g food pellet to eat. When the pellet was eaten (typically around 30 s), the rats were allowed 10 min to explore the arena. In the second and third habituation session, one 0.5 g pellet was placed on top of a sandwell placed two tiles away from the startbox; the rats collected this pellet and took it back to the startbox; they were then allowed to retrieve up to 6 food pellets buried close to the surface of the Sandwell for 10 min. If the animals had difficulties in retrieving the hidden food pellets, they were helped by exposing one further pellet in front of them (this happened rarely and was soon unnecessary).

### Behavior

Two behavioral protocols were employed based on the overall structure of the event arena task. The procedure of this task is that the experimental animals learn to retrieve food pellets from one of the 6 sandwells in the arena, their location changing from one session to the next (sandwell 1-6 labeled in West to East, North to South order). The concept of the task is that digging in a sandwell is an ‘event” and such events happen in different places across days. The overall aim was to conduct calcium imaging of the patterns of cell firing in the startboxes and the whole arena of this ‘recency memory’ task. To this end, Sessions 28 to 34 were recorded with miniscopes. Animals were habituated to the presence of the camera and cables during exploration of the arena while the miniscope was plugged in during the week before the start of the recordings. Asymptotic behavioral performance required training over many sessions. Three groups of animals were trained independently.

During the daily Exploration session, all possible sandwells were empty of both sand and food. The animals explored the arena for 2 blocks of 10 min. In the first of these, they explored without any reward present (the sandwells were empty of both food and sand). Only in the second block of 10 min was a small number of 45mg food pellets were scattered around to encourage sampling of the whole arena (F0021 Bioserv; distinct from the bigger pellets used in the main experiment). This phase was later used in the daily recording sessions to establish the location of numerous hippocampal CA1 cell spatial firing fields.

In the sample and choice trial phases, rats were initially disoriented by the experimenter and then put in one of four startboxes (orange in Figure 1) with their access to the arena controlled with an automated door. There were 3 pairs of trials (total = 6 trials), with the exception of sessions where recordings were performed, where 4 pairs of trials (total = 8 trials) were performed. In all trials, animals were given a cue pellet, then after 30s in the startbox, a 0.5s, 2300Hz tone was played; 5s later the door opened allowing the animals to enter the arena. As the animals have agency to exit the startbox, the trial start (0s) was set when animals started moving out of the door, and entered the arena (Gobbo *et al*., 2022).

In the allocentric protocol first developed by Broadbent *et al*., (2020), the combination of startboxes was randomized with the exception that combinations of three identical startboxes in a single phase were avoided (Gellerman rule (Gellermann, 1933)); see Supplementary Table). Animals started from one of three startboxes (East, South, West); after exiting from the designated startbox and retrieving one pellet from the rewarded startboxes, allocentric animals carried the 0.5 gm food pellet to the Goal Box (conventionally the North startbox) (end of trials 1,3,5), consume it, then return to the sandwell to retrieve a second pellet and return to North (end of trials 2,4,6). In the egocentric protocol, animals started from the same startbox in a given session, chosen randomly across days. All startboxes were used equivalently, with the exception of recorded sessions, where the North startbox was not used to ensure consistency with the allocentric-trained animals. On pre-determined sessions (S18,25,34), animals in both protocols were tested on their reliance on cues by performing a “curtain trial” added at the end of the choice phase. All intra-arena cues were removed, curtains pulled around the arena to obscure distal cues, and a single trial was run. To promote the use of an egocentric strategy, the first 18 sessions were performed in absence of cues with closed curtains to preclude sight of distal cues, as repeating trajectories and cue-poor environments have been demonstrated to facilitate the development of egocentric strategies (Scharlock, 1955). From session 19 onwards, the arena looked identical for the animals in the allocentric and egocentric protocols. Appropriate controls were run to ensure the use of an egocentric strategy. First, “curtain trials” were performed after the choice phase on sessions 25 and 34: these were identical to choice trials but without cues. Second, “180° rotation trials” were performed on sessions 16, 24, and 32 at the end of the choice phase; in these trials the starting position of the rat and the position of the rewarded well were rotated by 180 degrees with respect to the centre of the arena. An egocentric strategy should not be disrupted by such a manipulation.

In both protocols, the artifactual use of olfactory cues rather than spatial recency memory was tested by performing “probe” trials 90 min after the sample phase as described earlier (Broadbent *et al*., 2020; Gobbo *et al*., 2022). Briefly, all sandwells were arranged to only contain pellets in the inaccessible area and covered with sand. The total amount of time spent by animals over 120s digging at correct and incorrect sandwells was counted.

At the end of each protocol (phase 1), animals were given 2 weeks rest and then switched to the opposite paradigm (phase 2) in a counterbalanced crossover design. In this way, we secured within-subject comparisons of the use of allocentric and egocentric strategies within animals. In the sample phase, only one of the six possible sandwells contained the food reward while the other 5 remained empty. In the choice phase, all 6 sandwells looked identical and were filled with sand. An accessible food pellet was present only in the designated (“correct”) Sandwell, that was rewarded in the sample phase. The remaining sandwells had an equivalent number of food pellets but were not accessible to the rats. In the choice phase, the rats were tested for their memory of the correct location that day, measuring the number of errors in getting there and the time to find the correct sandwell.

Sessions 28 to 34 were recorded with miniscopes (see above). Animals were habituated to the procedure and to explore the arena while the miniscope was plugged in during the week before the start of the recordings.

### Electrophysiology

Electrophysiological recordings with optogenetic stimulation were performed under non-recovery urethane anesthesia (1.3 g/kg body weight), at least four weeks post-injection. A craniotomy was made above the left hippocampus and an optic fiber (0.5 NA, core 400 µm SMA (FP400ERT, Thorlabs)) connected to a 625 nm fiber-coupled LED (M625F2, Thorlabs) was inserted above dorsal CA1 (AP 3.5, ML -2.6, DV -1.7). Next, a 32-channel 4-shank silicon probe (Buzsaki32, NeuroNexus) was inserted immediately posterior to the optic fiber, with shanks aligned with the proximo-distal axis of CA1. The ground and reference wires were connected to a skull screw above the cerebellum. Broadband neurophysiological signals were acquired at 30 kHz on an acquisition board (OpenEphys) with a 32-channel RHD headstage (Intan Technologies). The probe was lowered in 50 µm increments over several minutes until unit spiking activity characteristic of CA1 pyramidal cell layer was observed. The probe was left in place for at least 1 hour before data collection and probe location in CA1 pyramidal layer was further confirmed by presence of SWR events. Optogenetic stimulation protocol consisted of 20 min of baseline recording followed by 10 cycles of square pulse stimulation (180 s pulse length, 360 s duty cycle). Animals were perfused transcardially as described above immediately after the experiment and anatomical locations of the fiber and probe tracts were further confirmed histologically.

### Optogenetics

In optogenetic experiments, animals were trained for 26 sessions with the egocentric or allocentric protocol as described above. In the last week, they were habituated to being connected to optic fibers and move in the arena space while connected. The effect of CA1 optogenetic inactivation on performance was evaluated in sessions 27 to 34. During these sessions, 90 min after learning the position of the rewarded sandwell in the sample phase, the rats were connected to one 3.1m fiber optic patch cable 0.5 NA, core 400 µm SMA/FC2.5 (FT400EMT with FT020-BK tubing, Thorlabs) for each hemisphere via Ceramic Split Mating Sleeves (ADAF1, Thorlabs). Each fiber optic cable was connected to a 625nm fiber coupled LED (M625F2, Thorlabs) mounted on rotary joints. On/off illumination was controlled with a custom program. In sessions 27 and 29 LEDs were activated when animals were put in the designated startbox upon receiving the cue pellet, and switched off as entry to the event arena occurred as the rats left the startbox. In sessions 31 and 33, LEDs stayed on until the end of the trial (hence corresponding to the time spent in the startbox plus the time in the arena representing the choice execution. Sessions 28,30,32,34 served as light OFF sessions where the procedure was identical, except LEDs remained switched off (hence alternating light ON and light OFF sessions). choice trials in Sessions 35-37 were run as usual. In sessions 35 and 37 probe trials were run as described above after connecting animals to optic fiber cables; in session 37 LEDs were switched on for the whole duration of the probe trial (starting in the startbox), while session 35 was the corresponding light OFF control. Light power was measured at the beginning and at the end of each session and was 17.20±0.69W (mean±st.dev) at the end of each fiber. Two groups of animals were trained independently.

### Data Analysis

#### Behavior performance

For each trial, latency and number of errors were quantified. Latency is the time occurred between the moment the animal leaves the startbox to when it starts digging at the correct sandwell. The number of errors is the number of incorrect sandwells the animal dug at before reaching the correct sandwell, and in this experiment can assume values 0 (the correct sandwell is the first choice), to 5 (if the animal dug at all sandwells before reaching the correct one). The Performance Index (PI) was defined as

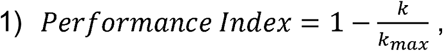

where is *k* is the number of errors. Hence if the animal performs at chance between *n*=6 sandwells

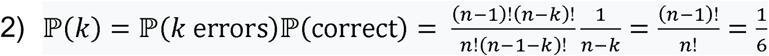

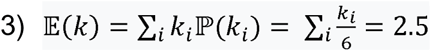

which is equivalent to a Performance Index of 0.5. In probe trials, the amount of time spent digging at any Sandwell was measured and expressed as a fraction of the total digging time.

#### Single-Photon Calcium Imaging

Inscopix data processing software (IDPS; Inscopix v1.6) IDPS software environment was used to process calcium imaging recordings and identify cells as described in Gobbo *et al*. (2022). Briefly, videos were downsampled spatially 2x2, background subtracted (spatial bandpass σ _low_ = 0.0005, σ _high_ = 0.5) and motion corrected. Cell regions of interest were detected with CaImAn constrained non-negative-matrix factorization-extended (CNMF-E) python API. Cell traces were expressed as ΔF/F = (F - F_0_)/F_0_, where F_0_ is the mean fluoresce over the trace. ROIs and their respective traces were manually inspected, with any duplicates or artefacts discarded. Neural events were then computed from the calcium traces using the OASIS (Online Active Set method to Infer Spikes) package (Friedrich *et al*., 2017; Giovannucci *et al*., 2019). OASIS parameter “s_min” was adjusted between 0.2 and 0.3 depending on the baseline noise of the trace. Longitudinal registration was performed with IDPS. Individual recordings from the same session were first aligned to each other generating one cell set for each session, then the resulting cell sets were aligned together. Correct alignment was verified by screening a random subset of individual cells from the correspondence table generated by the system.

#### Animal positional tracking

Behavioral recordings of the task are performed with a camera placed on the ceiling (20fps). Two separate input systems were used to align the camera video and the calcium recordings. In both systems, they synchronously switched on small LED lights within the camera field positioned outside the arena and an electrical signal timestamped into the nVista GPIO channels. Positional trajectories were tracked using the Python image recognition deep convolutional neural network DeepLabCut (DLC) (Mathis *et al*., 2018) as described previously (Gobbo *et al*., 2022). A second-order training of the DLC network from a dataset from various rats was performed with a total of 700,000 iterations. Results were also visually inspected and automatically corrected with a custom code to remove low-likelihood and incorrect points. The position of the animal is translated using one corner of the arena as reference origin.

#### Automated trial detection

To extract trial-wise behavioral trajectories, we implemented a computer vision pipeline that automatically identified key spatial features from each behavioral video. Using a custom OpenCV-based method, six circular sandwells (labelled 1-6) with a 3.85cm radius were detected via Hough Circle Transform and verified through manual inspection. The arena’s boundary was determined by localizing high-contrast corner features adjacent to the sandwells using cv2.goodFeaturesToTrack. DLC (x, y) positions of the rat were transformed into arena coordinates by computing a bounding box between identified corners. Startbox zones (N,E,S,W) were defined relative to arena boundaries. Trial epochs were extracted as continuous sequences of frames commencing from the moment the animal’s head exited the startbox and terminated when the animal re-entered the startbox (or entered another startbox if allocentric). Using the defined arena features, a behavioral vector was also created for each trial split by startbox (N,E,S,W), trajectory and RW.

#### Place cell identification

For each neuron, spatial information (SI) was calculated using the Kraskov algorithm to measure the mutual information between the neuron’s binarized neural event train and a vector of the animal’s linearized spatial positions. The arena was divided into spatial bins with a 10 cm bin size, and spatial occupancy and place fields were calculated by normalizing firing events with the animal’s occupancy in each bin. A neuron was classified as a place cell if it met the following criteria: (1) the neuron was active in at least three recording sessions, (2) the neuron exhibited more than three firing events during these sessions, (3) the animal traversed the bins in which the neuron was active at least three times, (4) a neural event occurred in at least 20% of the traversals, and (5) the neuron’s SI exceeded the 99th percentile of a distribution generated from 5000 shuffled neural event trains.

#### Directionality index

A neuron’s directional index was computed based on its firing rate over four cardinal directions. For each recorded frame, the animal’s head direction was calculated using x and y coordinates of the left and right ears (extracted using DLC, see earlier methods). The horizontal (*dx*) and vertical (*dy*) distances between the ear coordinates were used to define a vector representing the head direction. The angle between this vector and the positive x-axis was computed using the arctangent function, atan2(dy, dx), and converted to degrees. Angles were then categorized into cardinal directions as follows: *North*: 0° ≤ θ < 45° or 315° < θ < neuron, the firing rate, *r* was computed for each cardinal direction and the 360°, *East*: 45° ≤ θ < 135, *South*: 135° ≤θ< 225, *West*: 225° ≤ θ < 315. For each directionality index calculated according to the following formula, general for *n* directions (Rubin *et al*., 2014):

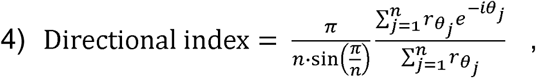

that for n=4 reduces to:

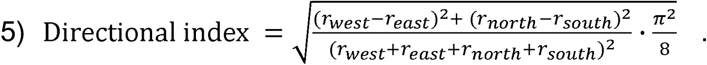

The resulting index ranged from 0 to 1, where values close to 0 indicate low directional selectivity (indicating uniform firing across directions) and values close to 1 indicate strong directional selectivity (indicating a preference for a specific axis).

#### Bootstrapping

Many cells were recorded simultaneously as each rat explored the environment. Because cells within the same recording or from the same rat may exhibit correlated responses, a hierarchical bootstrap resampling procedure was performed to account for this non-independence and the unequal number of cells recorded from each rat. Bootstrap resampling was performed at two levels: within recordings (cell level) and across rats (recording level). For each recorded population, cells were resampled with replacement to yield a standardized sample size equal to the median number of cells per recording. Recordings containing fewer than 20% of this median were excluded to prevent oversampling of small populations. Resampled allocentric and egocentric recordings were then paired and drawn with replacement across rats to yield datasets with equal numbers of rats (n = 9). The mean difference between allocentric and egocentric conditions was calculated from each resampled dataset. This procedure was repeated 10,000 times to generate a bootstrap distribution representing uncertainty due to hierarchical sampling structure. Two-tailed statistical tests were performed by calculating the proportion of bootstrap replicates where the allocentric-egocentric difference that was either greater or less than zero, whichever was smaller. This essentially provided a *p*-value of whether interval included zero.

#### Reactivation consistency

The reactivation consistency of neurons active in the startbox before a trial was computed for both navigational strategies within and across sessions. For each animal, correct trials were identified and the pre-trial startbox activity isolated, i,e the 10 second window before the animal entered the arena. For each trial, the startbox neurons were catalogued and their reactivation consistency compared across all other trials. The reactivation consistency was recorded as the number of common cells active in the startbox pre-trial 10s window. The analysis was conducted within sessions and across sessions. Within session reactivation compared activity from the same day’s recording session, whilst across sessions compared reactivation activity across all recording sessions.

#### Content of reactivation

For each reactivation instance, we plotted the event maps showing the activity of the cell in that session and, separately, in all sessions; only points where the animal was in the arena were considered. An experimenter blind to conditions annotated the information content. Cells with >3 event in that sessions within a 6 cm radius from the Sandwell centre were considered to represent that Sandwell. If more than one Sandwell met the criterion, that cell was considered to represent multiple sandwells. If no events mapped the sandwells, that cell was classified as ‘other’. The number of ‘other’ cells was not considered when expressing the percentage of unique cells (e.g. Fig 3b). Handling and statistics on annotated data were performed in Python. For display purposes, we display maps as rate maps rather than event maps in pink-to-blue lookup tables. Comparison with event maps is displayed in the Supplementary materials.

#### Reward and trajectory encoding

For each animal and experimental phase (allocentric or egocentric), during individual trials from SAM and CHO, we assessed spatial tuning of “startbox neurons” to goals and trajectories. Arena coordinates were extracted from representative behavioral videos, and the environment was discretized into 4 cm bins (5.2 pixels/cm) for spatial mapping. We selected neurons that had both startbox and arena activity and computed their place fields from binarized calcium events during the animal’s time in the arena. For each cell, we calculated the Euclidean distance from its PF centre to the reward zone and to the binned trajectory. Cells were classified as encoding the reward (RW), trajectory (Trajectory), or neither (Other) based on which feature the place field centre was closest to, and whether it was within a distance threshold of 20 cm (i.e. within 5 bins distance, the typical PF size in our experiment). Cells not meeting this criterion were assigned to the “Other” category. The classification was conducted for each correct trial trajectory and also for identical trial trajectories (trials with the same startbox and reward well).

#### Event frequency during Exploratory phase

For each animal and phase, we divided the arena in 10cm bins and calculated the place map by dividing the event rate map (events per bin) by the occupancy map (position per bin). Only moving points inside the arena were considered, and events were normalized to 0/1 before binning. Above-threshold activity bins were considered to estimate the place field with threshold=0.6 max value. The weighted centroid was calculated and defined as the PF centre for the cell. The distances to the sandwell centres in bin units were calculated and the closest sandwell was determined. If no sandwell centre lay within 20 cm in bin units, the cell was interpreted as not representing any sandwell and classified as “Arena”. For each cell, the Exploration event frequency per minute was calculated for moving points by diving the number of events by the duration of the interval. The resulting activity was compared between different classes of cells that were active in the 10s before the trial start. Startbox activity event frequency was calculated in an analogous way by counting the number of events in the 10 sec time window and expressed as events min^-1^.

#### Decoding sandwells from startbox activity

For each animal, we used as the training dataset the 20 minutes of free exploration of the arena at the beginning of the training session (stage “Exploration”). We labelled each time-point based on the position category the animal was in (possible positions: arena, SW1, …, SW6). Sandwell (SW) locations were considered as all points inside the circular area centred on the sandwell, with a diameter of 15cm. A decoder was trained from the event calcium activity of the animal to the labelled position: we used multinomial logistic regression classifiers (scikit-learn’s LogisticRegressionCV with 5-fold cross-validation) as our decoders, and the neural population activity was represented as firing rate vectors across all simultaneously recorded cells at each spatial location. Training was performed exclusively on arena locations (sandwells 1-6 and arena) with activity from non-arena periods excluded (Stratboxes). Moreover, we excluded all time-points where all the recorded cells did not have any calcium activity (all cells were zero), to avoid introducing a bias. The decoder was then applied to (test set) on the calcium activity from the startbox of the following stages (SAM and CHO), before the beginning of each trial (foraging to the correct rewarded well). The output of the decoder consisted of probability distributions over all locations, for each testing time-point. We chose this probabilistic approach as it allowed us to quantify the degree to which neural activity resembled different locations. The following parameters were calculated:

##### Sandwell overrepresentation

For each animal we calculated (i) training frequency: frequency of each location-class in the training set (Exploration stage), (ii) accuracy: percentage of correct predictions during the cross-validation on the training set, (iii) expected prediction: given the training frequency and accuracy, we can find the expected prediction for each location-class as:

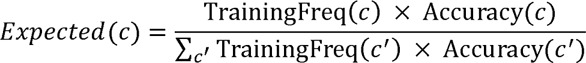

and (iv) Testing frequency: the frequency of each location-class in the test data (animal in startbox in SAM and CHO stages). Given these metrics we defined “overrepresentation” metric for each location-class as Overresentation_SW_ = Observed_SW_ – Expected_SW_, where “observed” is the testing frequency. This metric measures how much representation of each class is present in the neural activity from the testing set, compared to its frequency in the training set.

##### Uniformity of sandwells representation

To assess whether animals represented in the startbox specific sandwells more than others, we looked at two metrics: (i) Entropy: Shannon entropy of the normalised frequency distribution across the sandwells classes. From the testing predictions we removed the time-points that predicted the arena-class, and normalised the frequency of the sandwells-classes to obtain a probability distribution. Higher entropy indicates more uniform representation across sandwell classes.

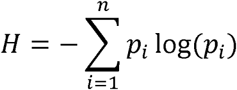

And (ii) Dominance of one class: we used the normalised frequency distribution calculated for the entropy analysis, and looked at the difference between the most frequent and second most frequent prediction. A highest number corresponds to a highest focus on one class above the others.

#### Binary naive Bayes decoder

To determine whether prospective goal location coding was stronger during the Allo or the Ego task, we trained spatial position decoders on the exploration sessions (Exploration stage) on each day and tested them on the task sessions (SAM and CHO stages) in the 10s period before trial start. The target variable was the goal location rather than real position. The decoders’ input was the neuronal data integrated for a given number of frames into the past and binarized. We treated the number of frames used for integration as a hyperparameter. The decoders’ output was a discretized position bin, converted into a position based on the position bin’s centroid. We treated the discretization lengthscale as another hyperparameter. The decoder itself was a binary naive Bayes model. For each exploration session, we used the first 70% of the session to fit decoders with various hyperparameter values, the next 15% as validation to select optimal hyperparameter values, and the last 15% as a test set for measuring the decoding performance. We assessed the performance by the root-mean-squared-error (RMSE) of the predicted bin centroids vs the continuous head position of the rat. The decoder RMSE was compared against a constant prediction of the center of the arena.

For decoding analysis, we included the four rats (H2224, H2225, H2226, and H2231) that exhibited a decoder test RMSE significantly different from the constant RMSE (See figure). Using the 10s period prior to trial start on the task sessions, combined with the spatial position decoder trained on that day’s exploration sessions, we measured the distance between the predicted location and the goal location (goal location prediction error). We present these distances as the RMS, mean, and minimum over the 10s period. We compared (i) the goal location prediction errors between correct and incorrect trials in Allo and Ego trials over the 4 rats with significant decoding performance, (ii) the errors during correct Allo trials and correct Ego trials.

#### Temporal clustering of events

We tested whether the temporal distribution of events for cells reactivated in the startbox was following a random Poisson-like distribution or instead showed increased clustering and synchronicity. For each trial, we calculated the event time of reactivations *N_T_* in the 10s interval before leaving the startbox and calculated the Fano factor:

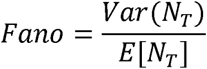

Where *Var(N_T_)* is the variance of the event times train density, and *E[N_T_]* is mean. Events were grouped in 0.5s intervals. A Fano factor F ≈ 1 implies a random Poisson distribution of events, while F > 1 implies clustering or synchrony. The inter spike interval (ISI) was calculated as the difference in seconds to the nearest event. The coefficient of variation was calculated for each trial as CV = σ(ISI)/µ(ISI), where σ is the standard deviation and µ is the mean of ISI times. For each trial, shuffle data was calculated by generating 1000 shuffled data distributions with a random uniform distribution of the event times matching the number of events of real data for the corresponding time interval (10s).

#### Neural correlation between trials

A trial was defined as the period from the animal leaving the startbox to reaching the correct reward-well, without pausing more than 0.5 seconds at any other well or taking longer than 5 seconds overall. Both training and memory recall sessions (Sampling and choice) were used to extract the trials. To compute the correlation between two trials we used Canonical Correlation Analysis (CCA), adapted in Python from MATLAB’s canoncorr function. This required pre-processing the data such that each trial pair had the same number of samples and features. Therefore, for each recording session we computed the PCA embedding of the events’ trains convoluted with a Gaussian kernel (σ=200 ms) that explained most of the variance (30 components, chosen empirically), and for each pair of trajectories we interpolated the PCA time-series to the average length of the two trials (using Numpy’s interp function).

We transformed the pre-processed trials with CCA (Canonical Correlation Analysis), finding a common space where the first components correspond to the directions of maximum correlation. We chose CCA because we aimed to compare not only repeated, identical trajectories, but also symmetrical trials (i.e. rotated by 180 degrees). We were initially agnostic regarding whether symmetric trajectories (especially in the egocentric case) would reactivate to any extent neurons representing the reference trajectory. Furthermore, even identical trajectories would not necessarily show 100% overlap between neurons, either due to detection limits, trial to trial variability in the pattern of neurons activated, or to representational drift, especially across days. CCA works by maximizing the correlations between two sets of signals through linear transformations, i.e. linear combinations of their neural dynamical activity vectors (Gallego *et al*., 2020; Safaie *et al*., 2023), yielding both the linear transformation, and the extent to which the signal sets can be mapped on to one another. We averaged the first 5 correlation coefficients, obtaining a quantitative measure of the correlation between the two neural trajectories. To validate the common CCA space, we decoded the position of the animal with a Gaussian Naïve Bayes decoder (from sklearn GaussianNB), cross-validated between pairs of trials. We repeated the analysis for each pair of trials in PCA (principal component analysis) space to compare the performance and confirm that spatial information was preserved in CCA space.

#### Electrophysiological data analysis

Spike sorting was performed semi-automatically using Kilosort 2.0 followed by manual curation of the waveform clusters using the software Klusters. At this stage, any cluster without a clear waveform and clear dip in the spike train auto-correlogram at the 0–1□ms time bin was classified as noise and cluster pairs with similar waveforms and a clear dip in their spike train cross-correlograms at the 0–1□ms time bin were merged.

Viable units were first identified as units that (1) had an average firing rate of at least 0.5□Hz during the baseline period and (2) had a waveform with negative deflection (criterion aiming to exclude spikes from fibers of passage). Next, putative excitatory cells and putative interneurons were classified on the basis of the through-to-peak duration of their average waveforms Putative interneurons were defined as cells with short trough-to-peak duration (< 0.5□ms) and, conversely, cells with long trough-to-peak duration (> 0.5□ms) were classified as putative excitatory cells. Statistical significance of modulation by light was assessed on a cell-by-cell basis by first calculating average firing rates during each light ON epoch and comparing them with average firing rates during each light OFF epoch with a two-tailed Wilcoxon signed rank test.

The wide-band signal was downsampled to 1.25□kHz and used as a local field potential (LFP) signal. For sharp wave-ripple detection, LFP signal was band-pass filtered at 150-300 Hz and the r.m.s. signal was smoothened with a 20 ms Gaussian window and z-scored. Sharp wave-ripple intervals were identified as intervals in which the resulting signal envelope exceeded 2 s.d. above the mean and the envelope peak was at least 5 s.d. above the mean. Events shorter than 25 ms and longer than 250 ms were discarded. The rate of sharp wave-ripple events was computed separately for each recording channel and the channel with the highest ripple rate was selected for further analysis. Ripple frequency was computed as the inverse of the average inter-peak interval within ripple events, and peak normalized ripple power was computed as the maximum value of the envelope signal within the ripple event.

#### Optogenetic illumination duration

For each video recording, a researcher blind to the experiment condition timestamped the beginning of each sample and choice by marking: (i) the moment the animal received the cue pellet in the startbox, and illumination was switched on in Light-ON trials, (ii) the moment the door od the startbox opened, (iii) the moment the animal exited the startbox and entered the arena – referred to as trial start in this work – (iv) any incorrect sandwell visit prior to (v) the moment the correct sandwell was reached, for each trial. Illumination time was calculated for each trial accordingly. “Decision time” was defined as the interval between the (i) cue-pellet reception and (iii) the trial start.

#### Blinding, Randomizing and Exclusion criteria

Where possible, data was analyzed with automatic algorithms that have no bias between groups. When users were involved in the quantification, they were blind to the Allo/Ego nature of the group. Users running behavioral experiments were blind to the virus group (Jaws/GFP) to which the various animals belonged. Behavioral data was quantified during the performance as it provides additional information than posthoc scoring from videos (e.g. the surface of sandwells can be checked to confirm digging) and were rescored by separate users blind to the group showing coherent values (Supplementary Figure S40) with live records. Extensive randomization was performed in the usage of sandwell and startboxes combinations (Supplementary Table S1). Physical sandwells and tiles were randomized as described in the Behavior section. Animals were purchased in groups and assigned randomly between groups. Rats in the calcium imaging group were initially screened for quality of field of view (visualization of active cells); rats with poor field of view were not recorded and the remaining ones assigned randomly between the egocentric or allocentric group. One animal (H2235) in the calcium imaging experiment broke the outer end of the GRIN lens before the recording of Phase 2 (See Supplementary Table S1). For one animal (H2222) the number of identified in Phase2 was deemed too low and excluded from analysis. For 3 animals in the allocentric-trained group (H2230, H2334, H2241) one of the sessions was not analyzed due to corrupted GPIO files, likely due to a technical problem on the day of recording. A limited number of trials where the beginning could not be automatically identified were excluded from the analysis. In the cohort used for optogenetics a small number of animals was terminated for health reasons unrelated to the experiment

#### Statistics

For behavioral analysis, two-way repeated measure ANOVA where applicable, with animals as within-subject factor. Kolmogorov-Smirnov test was used to test differences in distributions. When comparing data points within animals, linear mixed models were used to avoid pseudoreplication. E.g. we tested Percent_Cells ∼ Ego/Allo + Phase + (1|animal) (Fig.2c), Y ∼ Ego/Allo + (1|animal) (Fig. 2g, 3a,b), Y ∼ Ego/Allo + Correct/Incorrect + (1|Animal) (Fig. 3d,e), Y ∼ Correct/Incorrect + Time + Correct/Incorrect* Time + (1|animal) (Fig. 3g). Violin plots represent individual measures (e.g. trials), with dots representing animal averages. Boxplots inside violin plots reflect distribution quartiles using seaborn function violinplot. All data are biological replicate.

#### Software

Confocal and microscope images were open and processed with Fiji/ImageJ v2.1 (NIH); linear transformation of brightness and contrast were applied uniformly and equally to all compared images or channels. Camera videos were recorded with OBS Studio (https://obsproject.com/). Calcium videos processing was performed as described above using Inscopix Data processing Software v1.6 or v1.94 and publicly available or custom Python code. Python libraries Pandas, Numpy, Seaborn and OpenCV were used. Statistical analysis was performed with GraphPad Prism v9 or with Statsmodel.api library in Python (v 0.13.2). Electrophysiological recordings were processed and analyzed using MATLAB 2020b (Mathworks).

## Data and Code availability

Data will be made available at University of Edinburgh DataShare upon publication. Code is on Github at https://github.com/francesco-gobbo/Startbox-Activity-in-Egocentric-and-Allocentric-Tasks.

## Acknowledgements

We are grateful to Wellcome Trust for the financial support to the project, Grant n.207481/Z/17/Z “The retention of memory and creation of knowledge” (1 October 2017 - 32 May 2023) to RGMM. We wish to thank the members of the Schnitzer group for their support and constructive suggestions, and especially Thomas Rogerson and Jessica Rosa Maxey for their help and insight. We wish to thank the former and current members of the Inscopix team for their constant help and in-depth discussion to solve problems. We are particularly grateful to Diane Damez-Werno, Mark Turnbull and Fabrizio Stizia of Inscopix for their assistance and constant support. We wish to thank all the members of the Morris and Schultz lab and our colleagues at the University of Edinburgh for their support. Among them, Giuseppe Gava for help with code writing, and Emma Wood, Paul Dudchenko, Tara Spires-Jones and Nick Robinson for useful discussions. We wish to thank Hugo Spiers (University College London), Timothy Behrens, Francesca Pozzolo and the other members of the Behrens lab (University College London), Freyja Ólafsdóttir (Donders Institute, the Netherlands), Lisa Genzel (Donders Institute, the Netherlands), Sam McKenzie (University of New Mexico, USA) and Garrett Blair (New York University, USA) for their valuable insight and discussion. Last, we wish to thank the Bioveterinary Science Staff at the University of Edinburgh for their support and expertise, and looking after our animals’ welfare; we thank wholeheartedly Lynn Morrison, Caroline Quigley and Montse Rigol-Muxart.

