## Supplementary Material for "Navigational strategy dictates hippocampal representation of space in an everyday memory task"

for

### Supplementary Discussion

We wanted to understand if the there is a correlation in the neural activity between trials executed by animals navigating using the two different navigational strategies. Each trial was defined by the animal leaving the starting box and reaching for the rewarded well. Due to the nature of the analysis, we had to adopt inclusion criteria that led to include a fraction of Trials where a sufficient number of neurons were active and that were classified as "correct", i.e. the animal didn't stop more than 1s on any other well, and did not take more than 5s to reach the rewarded well. This was due to the need of trajectories to sample from a share spatial and neural space, whereby incorrect trials typically displayed a unique sequence of visited sandwells.

Given we imaged from different animals, we used CCA to find a common neural space where to compare the trajectories. We decided empirically that 5 components were enough to represent most of the neural activity. More components would have explained more variance, but would have led to the exclusion of a larger number of trials due to the low number of active neurons during each trial (a notorious characteristic of pyramidal cells in CA1). A low number of neurons lead to not fully ranked matrixes when building the CCA components, not allowing us to invert the transformation matrix, therefore preventing the projection of the neural activity from one space to another (the "alignment"). Given that CCA finds the components that maximise the correlation between two datasets, there is no guarantee that the firsts of these components preserve the information carried out by the full-higher dimensional neural activity. For this reason, we built a decoder for the position of the animal along the trial, and test that performance is similar when decoding from neural space or the first 5 components of the CCA space. We found that both spaces, neural and CCA components, carry a similar amount of spatial information, suggesting the "alignment" preserved meaningful components of the recorded neural activity (Fig:SD1.b,c,e,f). We also wondered if the difference in correlations between the trials from animals navigating allocentrically vs. egocentrically could be a consequence of biases in the included trials. We looked at the number of shared neurons IDs (Fig:SD1.i), and the similarity of the paths taken during the trials (Fig:SD1.g.h.m), without finding significant difference between the two groups. We also observed allocentric animals had a higher number of active neurons during the trials (Fig:SD1.l), and therefore a higher dimensional neural activity (less explained variance from the top 5 CCA components Fig:SD1.n). This could indicate that the lower correlations in the first 5 components of egocentric animals are a consequence of a "simpler", lower dimensional neural activity.

We then looked at the same results from animals in phase 2, the difference in number of neurons doesn't lead to the same strong difference in correlations, although the limited number of animals included in the CCA analysis for phase 2 prevents us from drawing firm conclusions. The limitation of this analysis lie in the lack of a large number of recorded active neurons during individual trials. Each trial is shorter than 5s and therefore only a few neurons are active during this time (Fig:SD1.l). Nerveless, we can consider Fig:SD1.a a preliminary result, with further experiments specifically designed to look at trajectory similarities and structure needed to confirm the initial findings reported here.

### Figure SD1

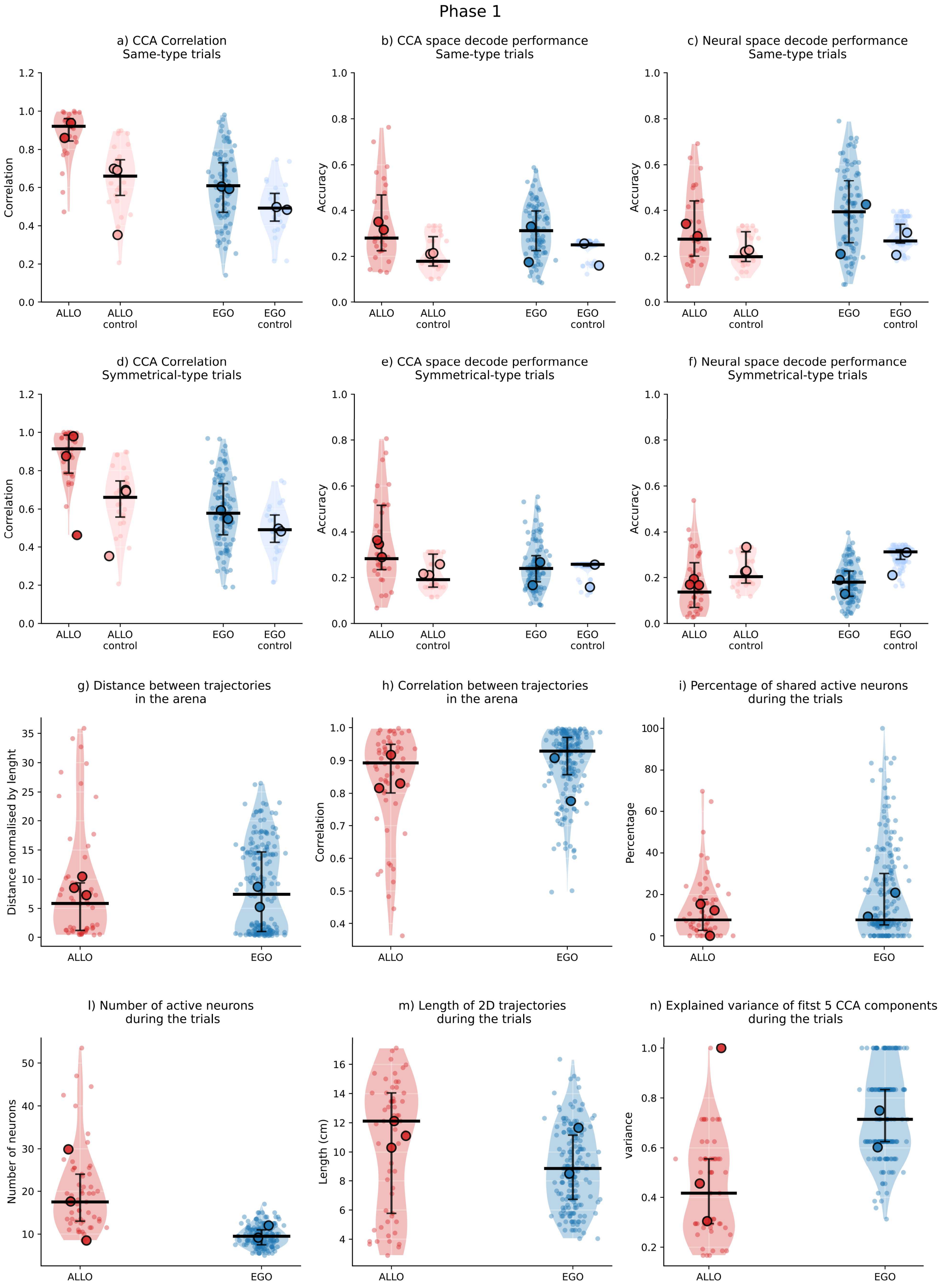
Phase 1
a) CCA Correlation
Same-type trials
b) CCA space decode performance
Same-type trials
c) Neural space decode performance
Same-type trials
d) CCA Correlation
Symmetrical-type trials
e) CCA space decode performance
Symmetrical-type trials
f) Neural space decode performance
Symmetrical-type trials
g) Distance between trajectories
in the arena
h) Correlation between trajectories
in the arena
i) Percentage of shared active neurons
during the trials
l) Number of active neurons
during the trials
m) Length of 2D trajectories
during the trials
n) Explained variance of fitst 5 CCA components
during the trials
Correlation
Accuracy
Distance normalised by lenght
Percentage
Number of neurons
Length (cm)
variance
ALLO
ALLO
control
EGO
EGO
control

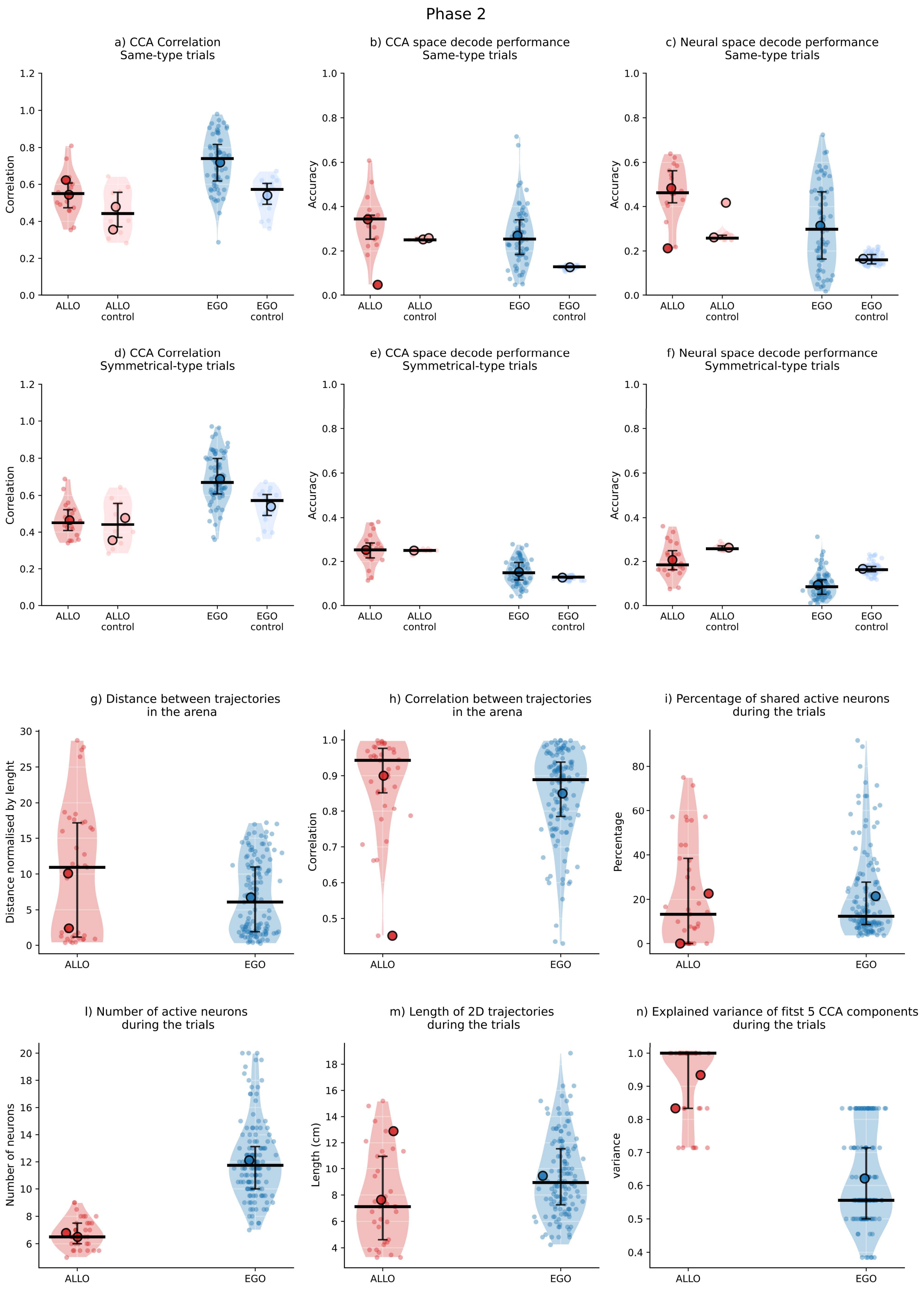
Phase 2
a) CCA Correlation
Same-type trials
b) CCA space decode performance
Same-type trials
c) Neural space decode performance
Same-type trials
d) CCA Correlation
Symmetrical-type trials
e) CCA space decode performance
Symmetrical-type trials
f) Neural space decode performance
Symmetrical-type trials
Correlation
Accuracy
ALLO
ALLO
control
EGO
EGO
control
g) Distance between trajectories
in the arena
h) Correlation between trajectories
in the arena
i) Percentage of shared active neurons
during the trials
Distance normalised by lenght
Percentage
l) Number of active neurons
during the trials
m) Length of 2D trajectories
during the trials
n) Explained variance of fitst 5 CCA components
during the trials
Number of neurons
Length (cm)
variance

**Legend for both Phases:**

- a & d: Correlation between trials along the first 5 CCA components.
- b & e: Train a decoder on one trial and test it on the other aligned trial. Decode location of the animal from the neural activity projected on the CCA space.
- c & f: Similar to b & e, but using the neural space to train and test the decoder, instead of the CCA space.
- g: Point by point, distance between the position of the animal in the two trials. Distance normalised by the average length of the two trials.
- h: Correlation along the x and y components of the position of the animal along the two trials.
- i: Given the neurons active during the trials, percentage of shared neurons between the two trials.
- l: Number of active neurons during the two trials.
- m: Length of the trials in centimetres.
- n: Fraction of explained variance along the first 5 CCA components.

### Supplementary Figures

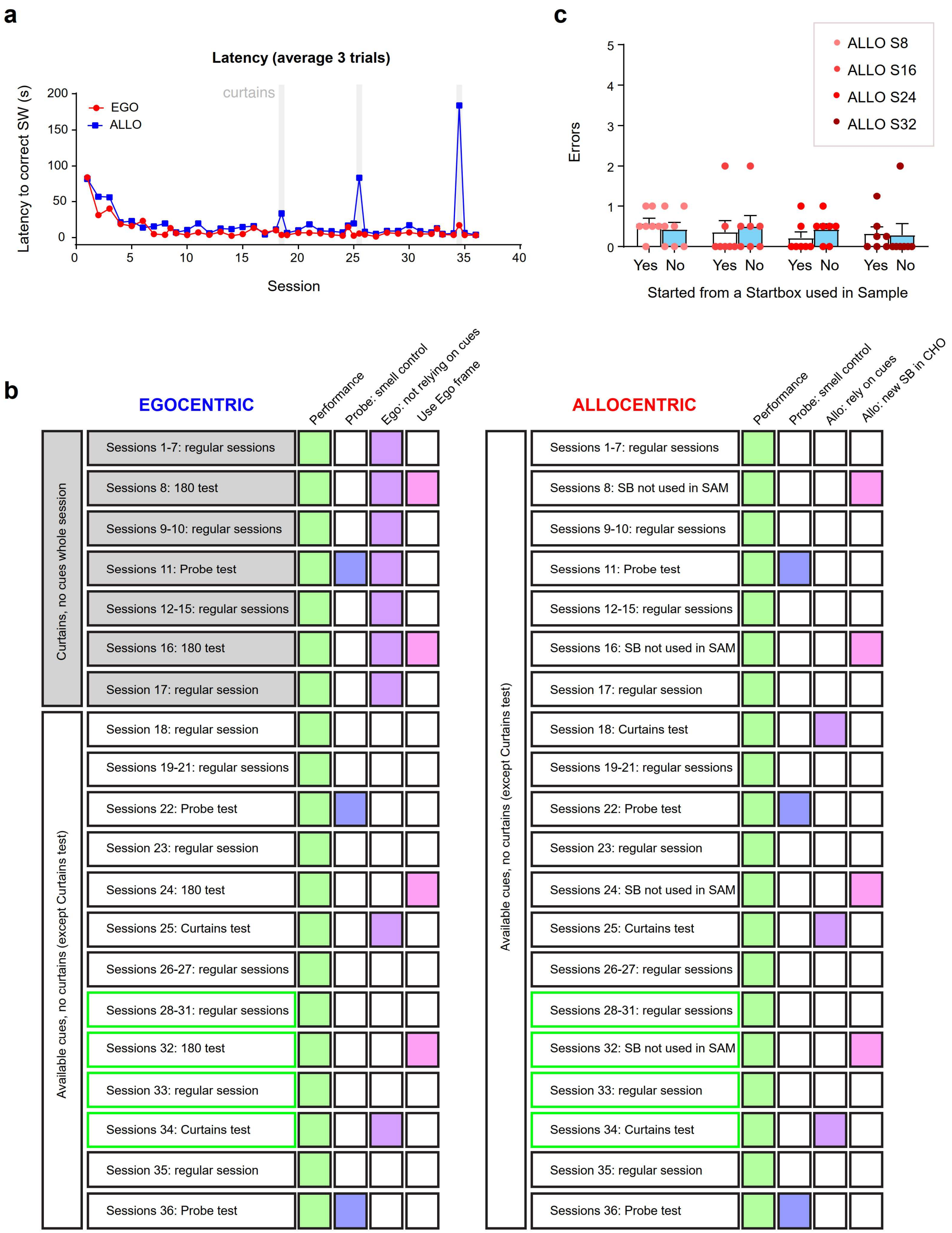

**Figure S1: Additional Behavior characterization**. a) Latency to reach the rewarded SW: Time between trial start as defined by the animal exiting from the Startbox (seconds) for Phase1 allocentric- (blue) and egocentric- (red) trained animals. Shaded areas indicate trials with curtains as in Figure 1. See also Methods. Data points are median values of animals displayed in Figure 1. b) Number of errors for animals in the allocentric group. In sessions 8, 16, 24, 32, all animals learned the position of the rewarded sandwell starting from only two possible startboxes in the Sample stage. They were later tested in Choice trials from the third, unvisited, startbox (blue bars), or from the other two (white). Within subject two-way ANOVA; startbox factor $F(1,6)=0.192$ $P=0.676$, session factor $F(3,18)=0.259$ $P=0.853$; session x startbox $F(3,18)=0.363$ $P=0.782$. c) Block diagram summarizing experimental details and what is tested at different sessions in Egocentric and Allocentric protocols. Sessions highlighted in green are sessions where miniscope recording was performed.

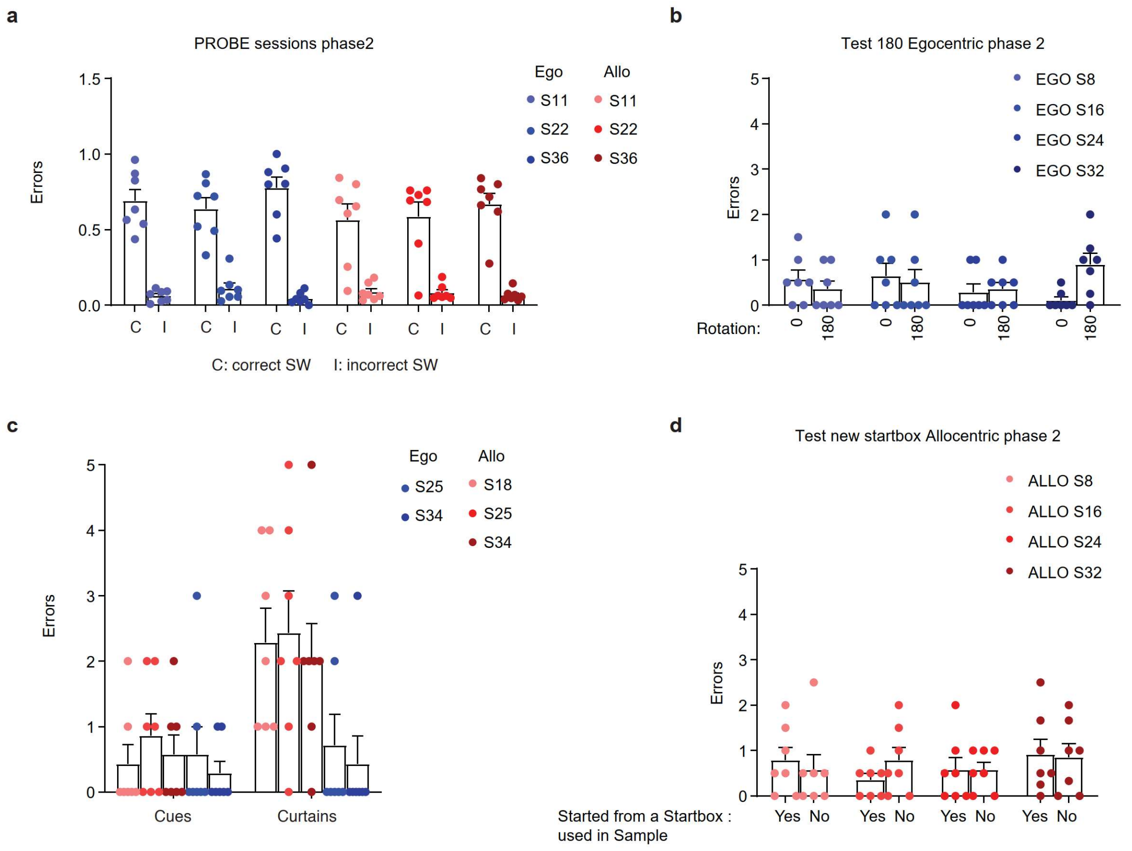

**Figure S2: Probe sessions and control trials for animals in Phase2.** a) Total dig time at correct (C) or incorrect (I) Sandwell in probe sessions for animals in Phase 2. Both groups dug significantly longer at the correct sandwell (two-way ANOVA: Ego $F(1,12) = 143.3$ $P<0.001$; Allo $F(1,12)=170.6$ $P<0.001$). b) 180 rotation test for egocentric-trained Phase2 animals. c) Number of errors with cues available or cues removed and masked with curtains for animals in Phase 2. Two-way ANOVA: Ego $F(1,12) = 0.091$ $P=0.768$; Allo $F(1,12)=14.95$ $p=0.0022$. d) Number of errors for animals in the allocentric group. In sessions 8, 16, 24, 32, all animals learned the position of the rewarded sandwell starting from only two possible startboxes in the Sample stage. They were later tested in Choice trials from the third, unvisited, startbox (No bars), or from the other two (Yes bars). Within subject two-way ANOVA; startbox factor $F(1,6)=0.025$ $P=0.880$, session factor $F(3,18)=0.692$ $P=0.568$; session x startbox $F(3,18)=0.657$ $P=0.589$.

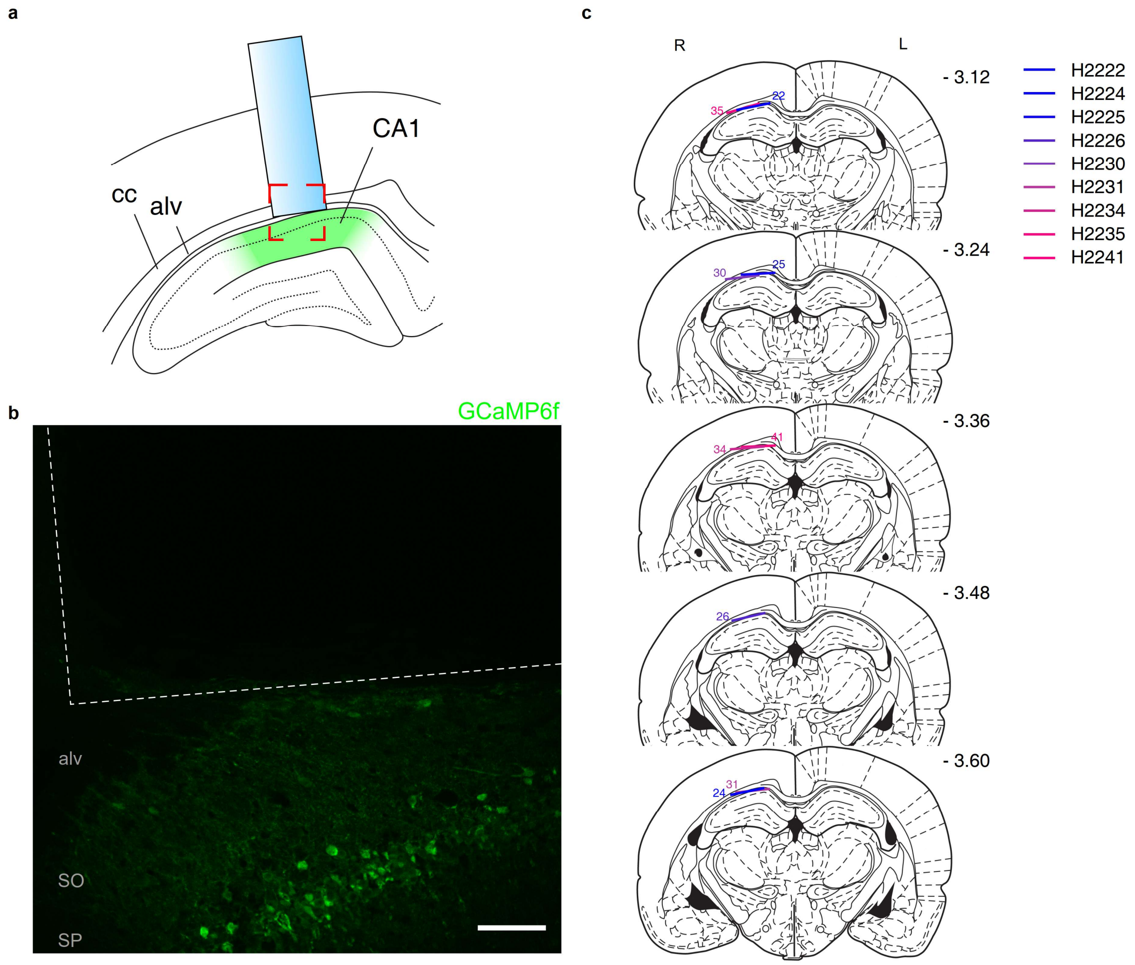

**Figure S3: Histology of rats implanted with GRIN lenses** a) Schematic of GRIN lens implant onto the outer surface of the hippocampus. CC: corpus callosum, alv: alveus fibers. b) Coronal section of recorded rat showing CA1 GCaMP6f fluorescence (green) and the lens tract above (dotted white line). Area is marked in red in panel (a). SO: stratum oriens, SP: stratum pyramidale. Scale bar, 100µm. c) Position of GRIN lenses for animals used in this study. Rat atlas coronal sections derived from (Paxinos & Watson, 2014).

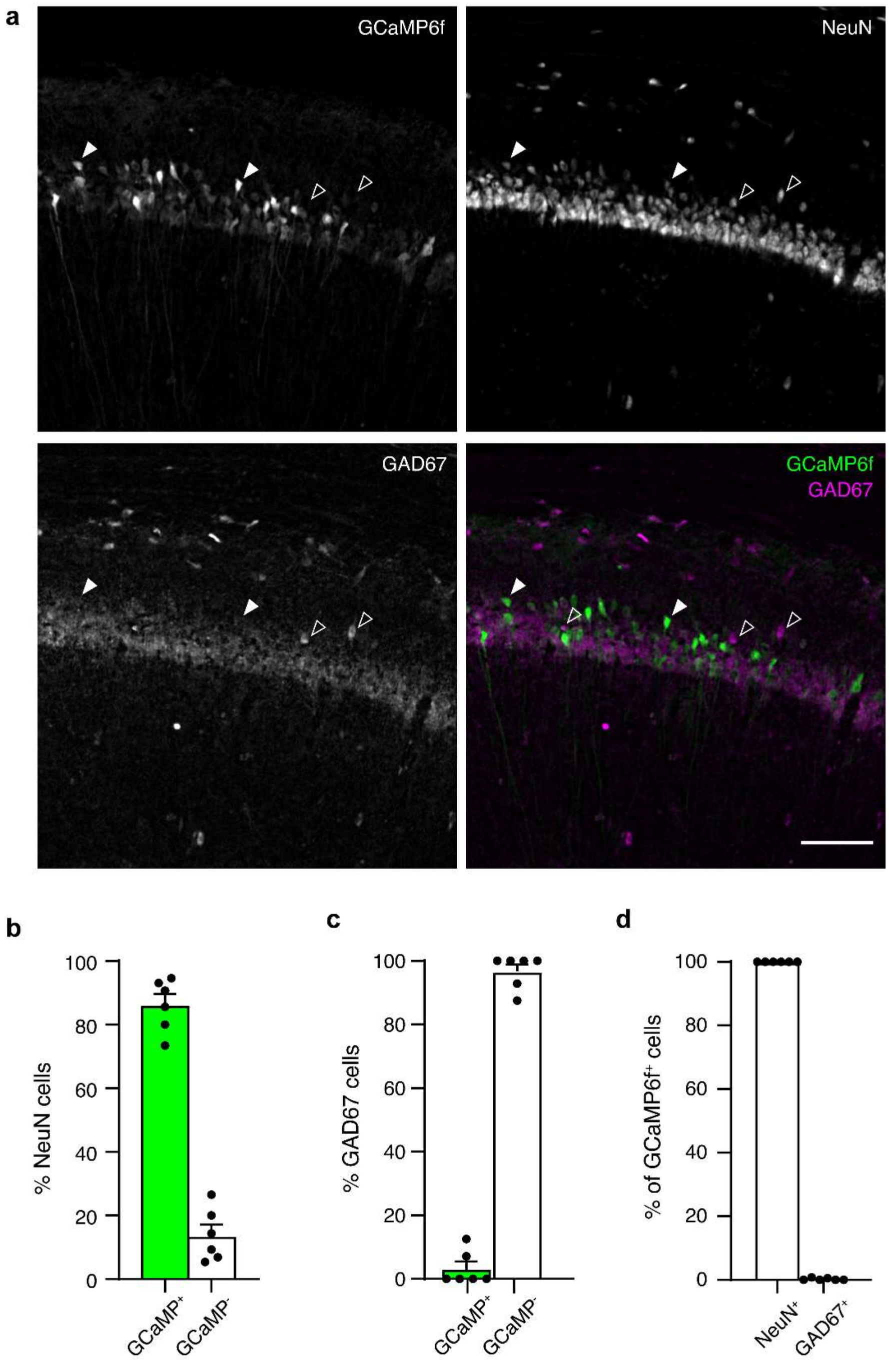

**Figure S4: Identity of recorded neurons** Expression in excitatory neurons in CA1 was confirmed histologically. a) Representative histology of a recorded animal showing GCaMP6f signal, NeuN and GAD67 immunostaining. Filled and empty arrowheads indicate example excitatory and inhibitory neurons across the different panels. Scale bar, 100µm. b) Percent of $NeuN^{+}$ neurons positive for GCaMP6f indicates the majority of neurons are infected. c) Only a minority of GAD67+ express GCaMP6f. d) Virtually all cells expressing GCaMP6f are excitatory neurons. N=6 animals, bars are mean±sem

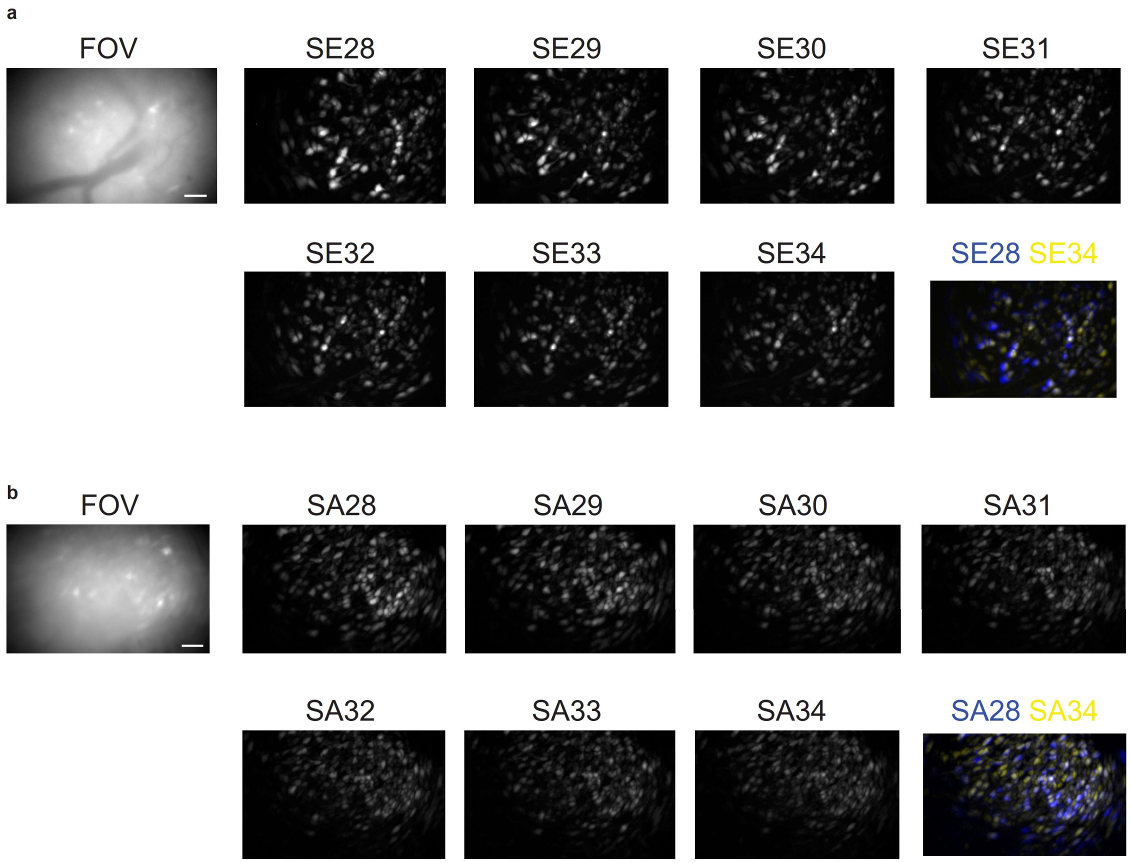

**Figure S5**: Field of view (FOV) and maximum projections of ΔF/F over consecutive recorded sessions 28-34 for an animal trained in the egocentric protocol in Phase1 (a) and one in the allocentric protocol (b). Scale bar, 100µm.

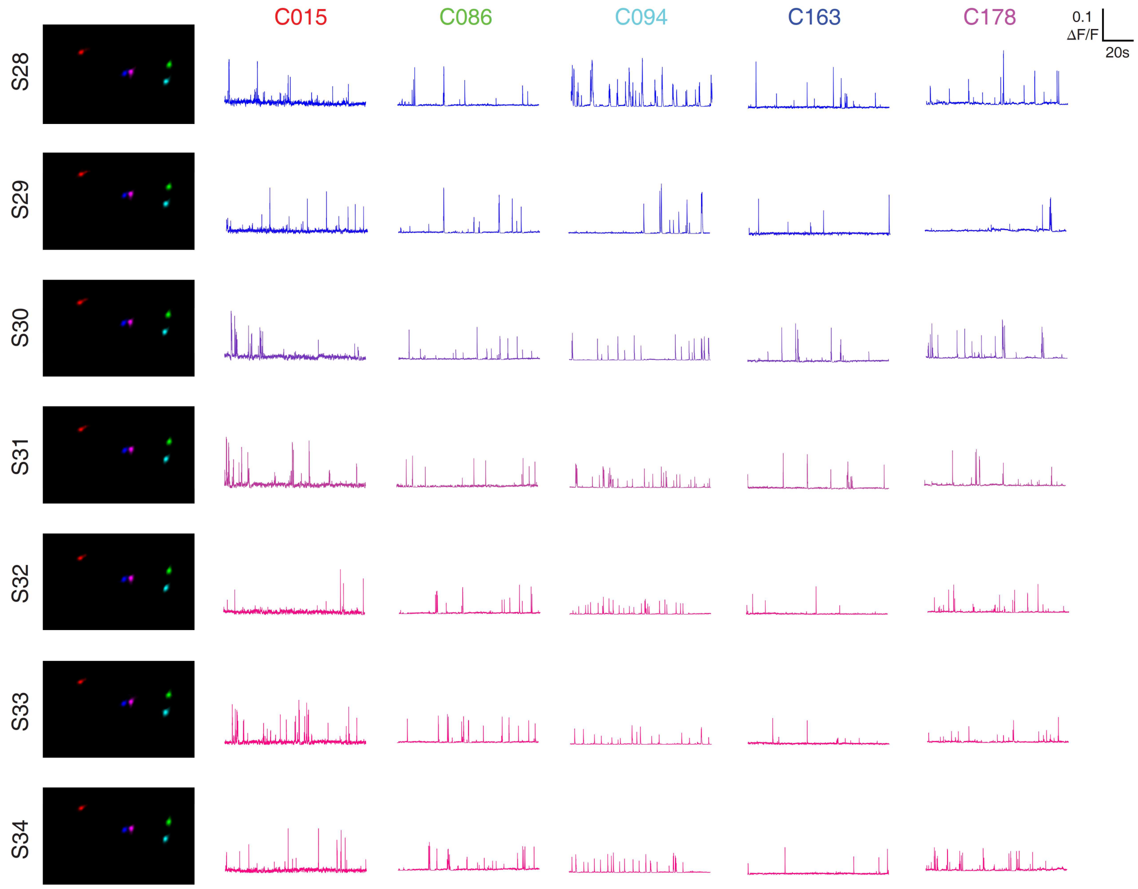

**Figure S6:** Traces and cell maps for representative cells confirming accurate alignment across recording sessions.

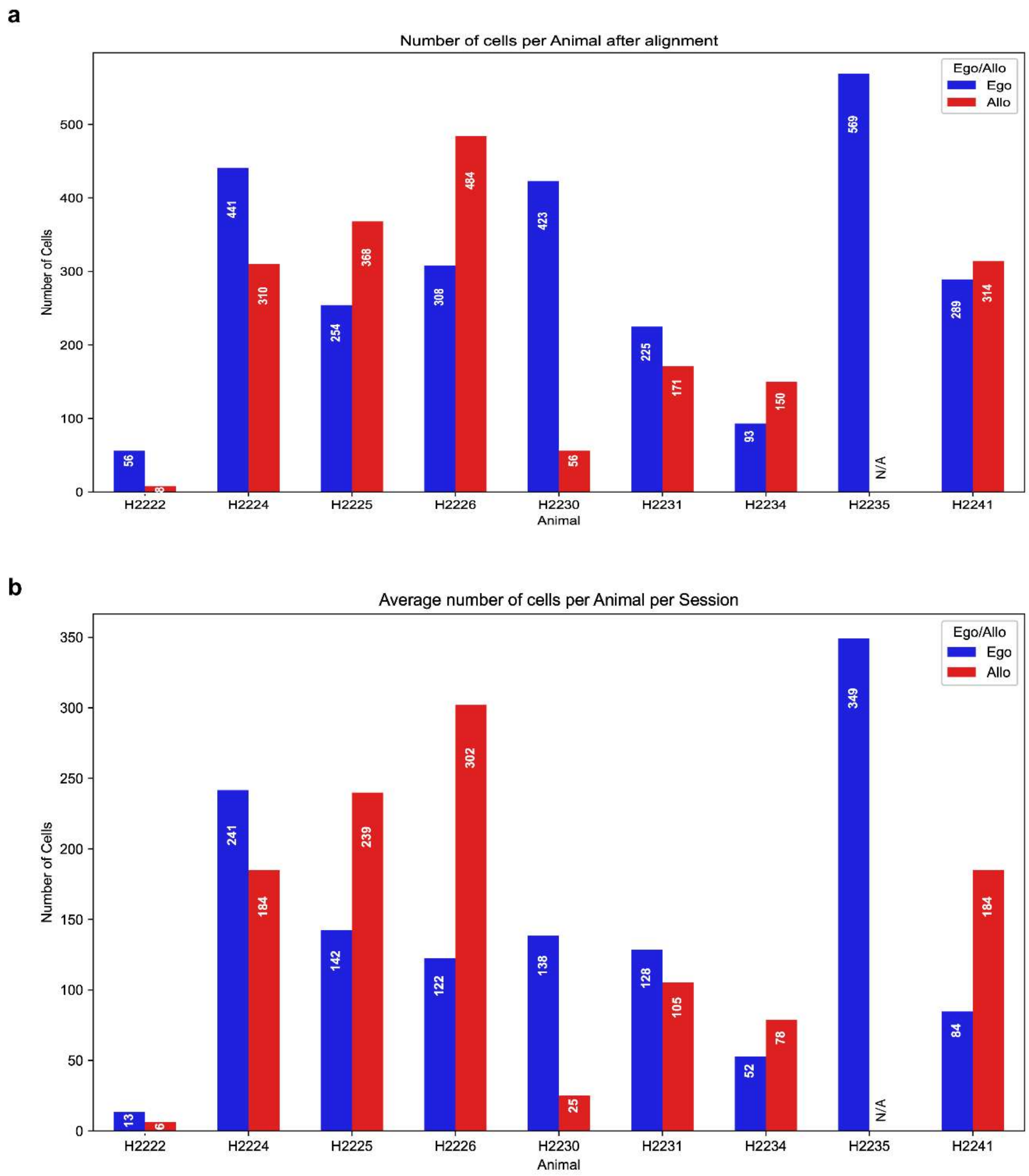

**Figure S7: Number of cells.** a,b) Number of identified neurons from animals included in the work. a) Number of individual neurons after longitudinal registration. b) Average number of neurons identified in each session.

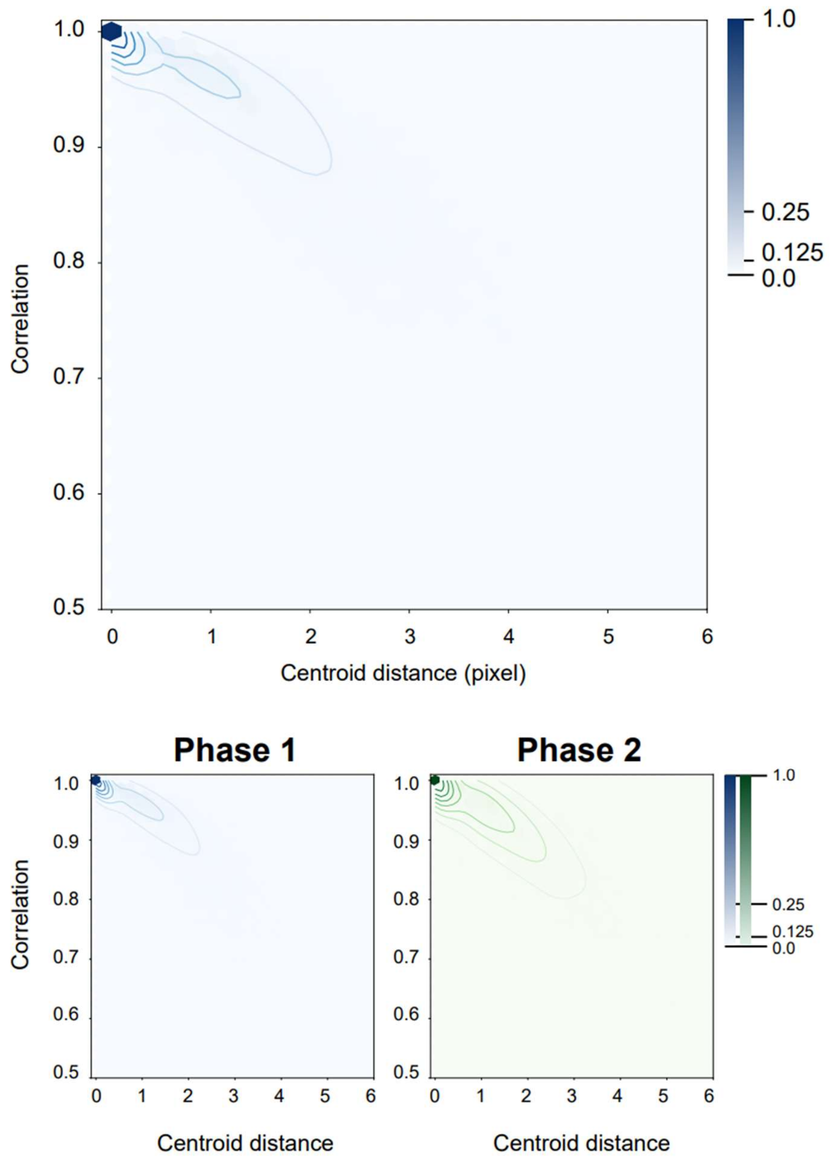

**Figure S8: Quality assessment of longitudinal registration of cell sets**. Top: pairwise Correlation/Centroid distance of region of interests (ROIs) aligned between cell sets. Bottom: Data in a) divided by alignments in Phase1 or Phase2 recordings showing no obvious difference between the two (see text). Hexbin density with superimposed level lines using seaborn library v 0.11.0.

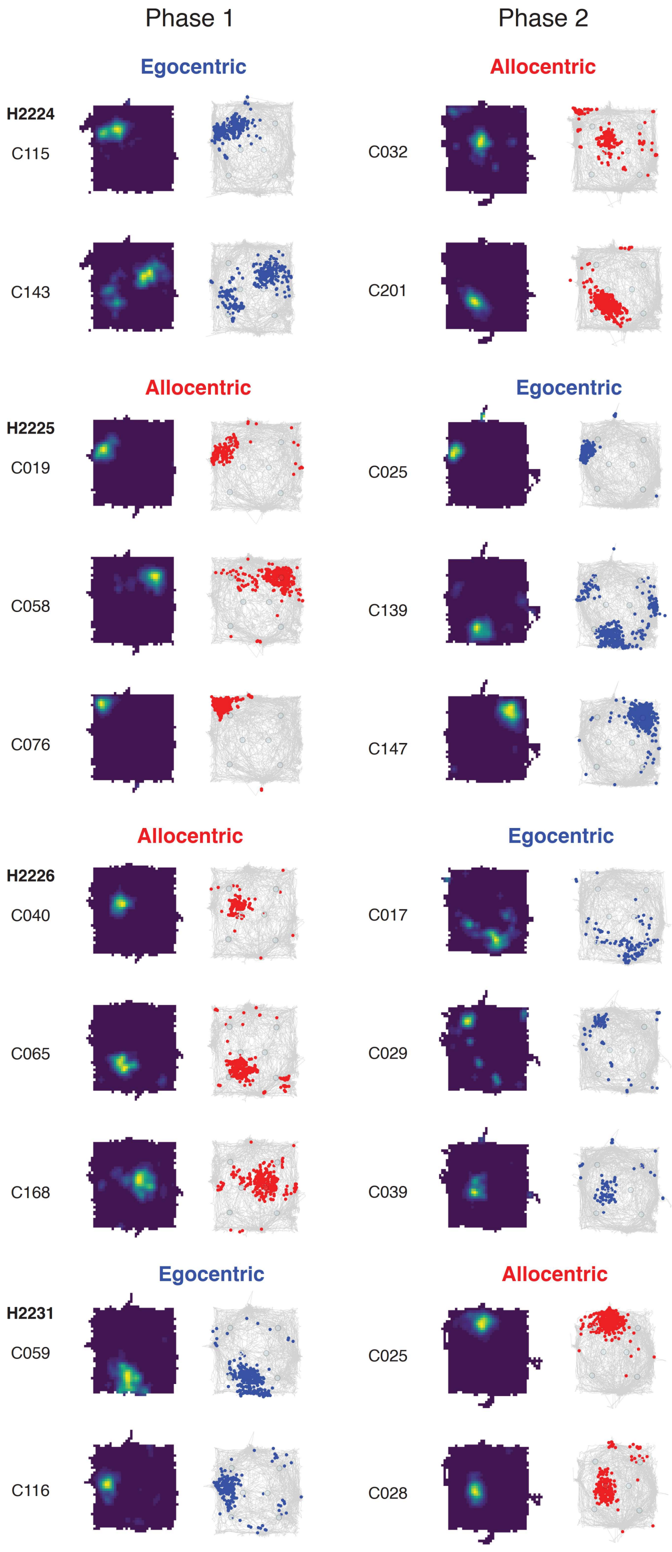

**Figure S9**: Representative place maps for cells recorded from animals trained in the egocentric and allocentric protocol. Allocentric-then-egocentric and egocentric-then-allocentric examples are shown. Red and blue dots are cell events in Allocentric and Egocentric groups. Rate maps represent event densities in blue-to-yellow.

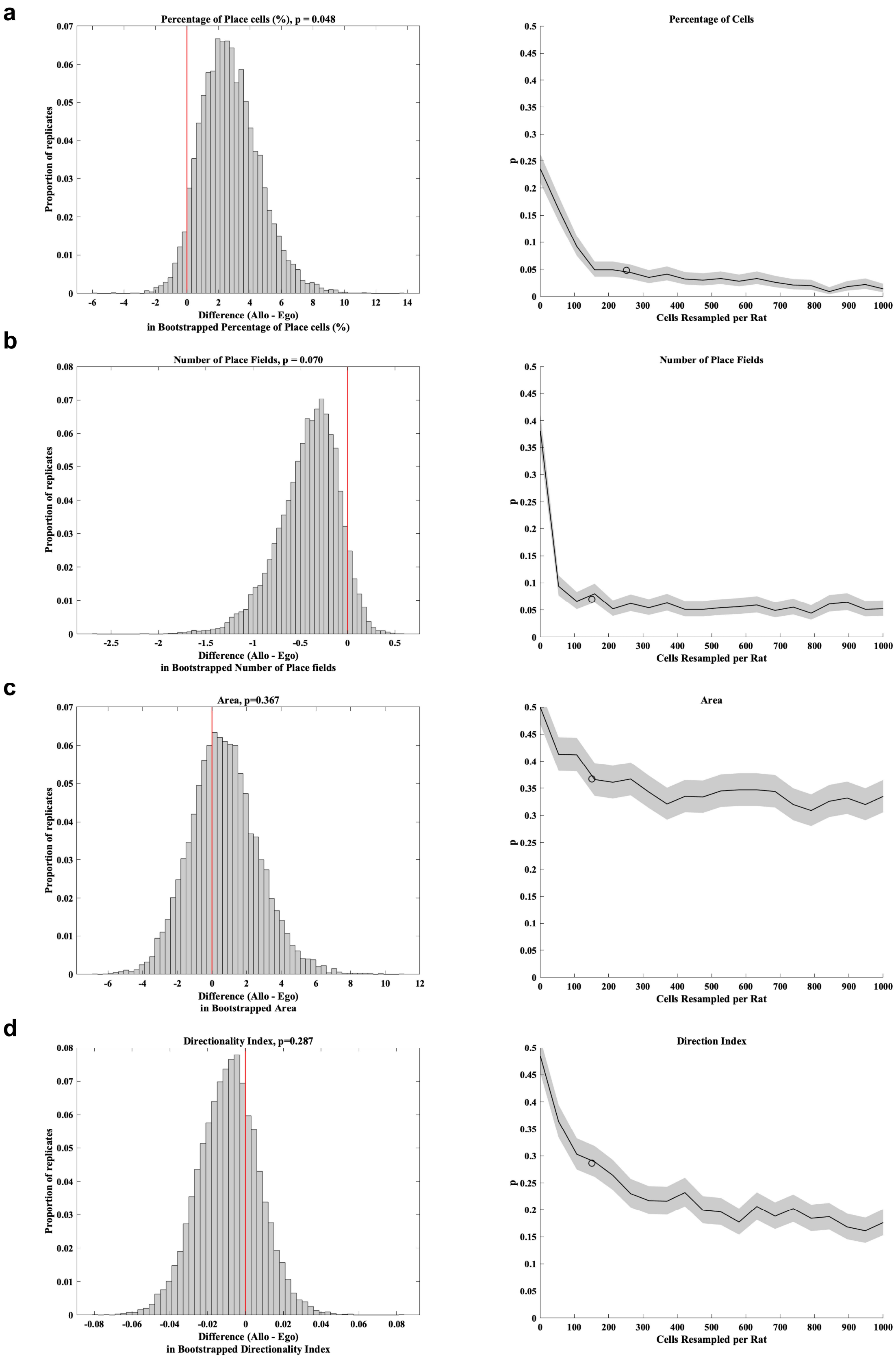
a
Percentage of Place cells (%), p = 0.048
Proportion of replicates
Difference (Allo - Ego)
in Bootstrapped Percentage of Place cells (%)
Percentage of Cells
p
Cells Resampled per Rat
b
Number of Place Fields, p = 0.070
Proportion of replicates
Difference (Allo - Ego)
in Bootstrapped Number of Place fields
Number of Place Fields
p
Cells Resampled per Rat
c
Area, p=0.367
Proportion of replicates
Difference (Allo - Ego)
in Bootstrapped Area
Area
p
Cells Resampled per Rat
d
Directionality Index, p=0.287
Proportion of replicates
Difference (Allo - Ego)
in Bootstrapped Directionality Index
Direction Index
p
Cells Resampled per Rat

**Figure S10 (previous page)**: Bootstrap analysis of Percentage of Place cells (a), Number of Place Fields (b), Place field Area (c) and Directionality Index (d). Left panels, Bootstrap distribution calculated for resampled means, and red line indicate real data. Right panels, p-value as a function of number of cells. Circles indicate average number of cells in our dataset.

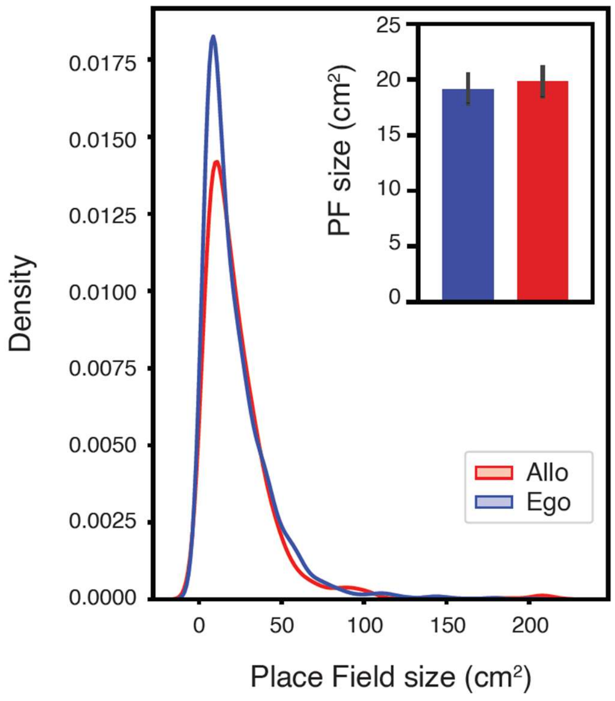

**Figure S11**: Distribution of Place field sizes recorded from animals trained in the allocentric or egocentric protocol. Inset, mean ± 25th-75th distribution of resampled means.

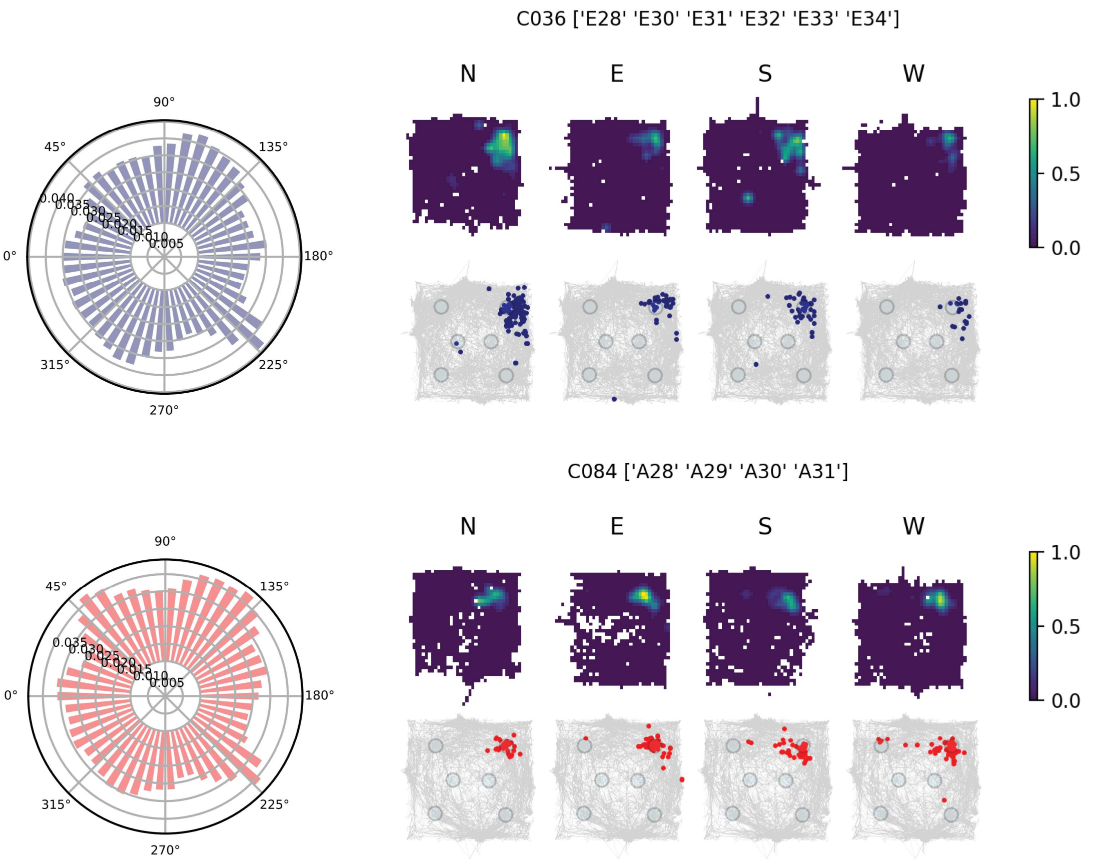

**Figure S12: Directionality of Place cells** Left panel, head direction distribution of animals in the Exploration of the arena in recorded sessions. Right panel, representative cells maps divided according to the head direction of the animal. Gray trace is the overall animal trajectory, and dots represent events divided according to animal's HD. Above, density maps of events normalized by animal's occupancy in each direction.

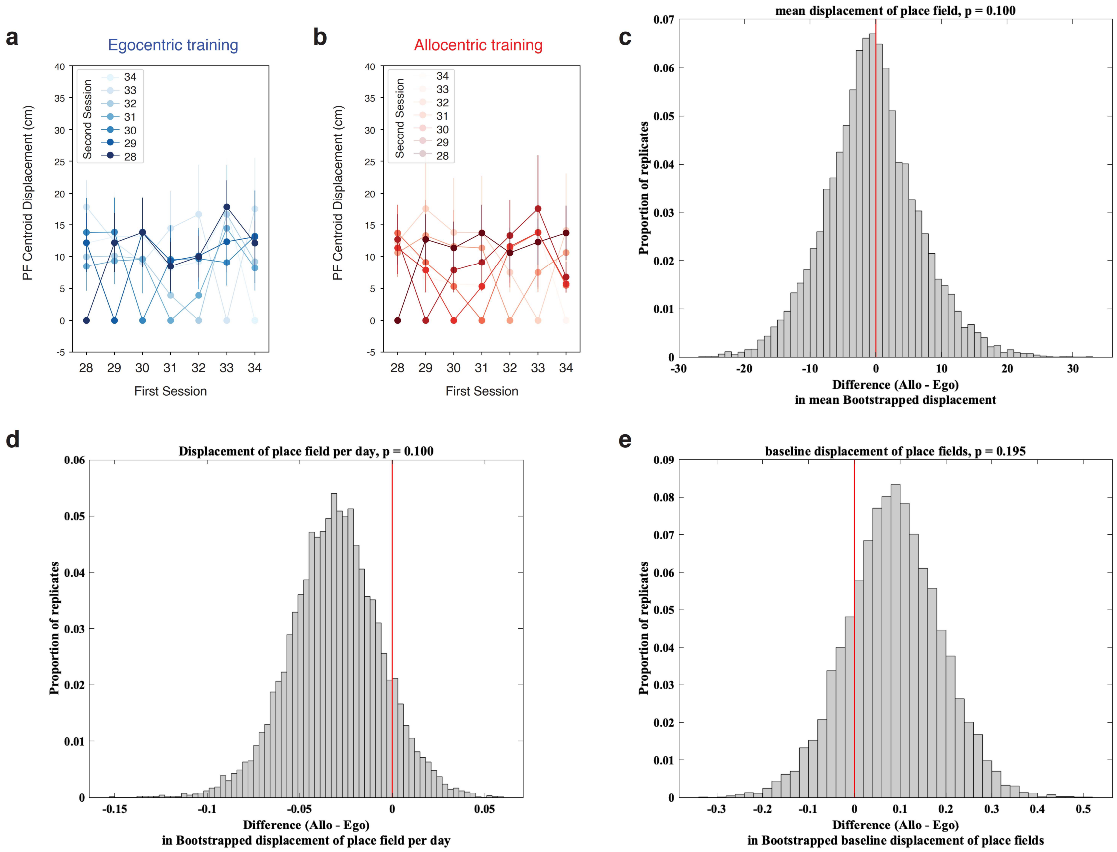

**Figure S13: Consistency of Place cells** Average pairwise session-to-session displacement of Place Field centers in cm for animals trained in the egocentric (a) and allocentric (b) protocols. Points are mean±sem. (c-e) Bootstrap analysis of displacement. In (c) we consider the option of random displacement as opposed to stability. We treated the choice of pair of days as an element of the hierarchy, so resampled which ones we were choosing along at a layer lower than rat (so we resample neurons, then day pairs, then rats), and look at average displacement. In (d,e), we considered the drift of place cells, or the accumulation of differences as time goes on. A linear model for each cell for displacement over time ($y = d\,x + e$) was fit. We called these d: instability, e: change rate and looked at whether they change as a function of time. We displayed instability and change rate in (d) and (e) respectively. No significant differences were found between the two groups.

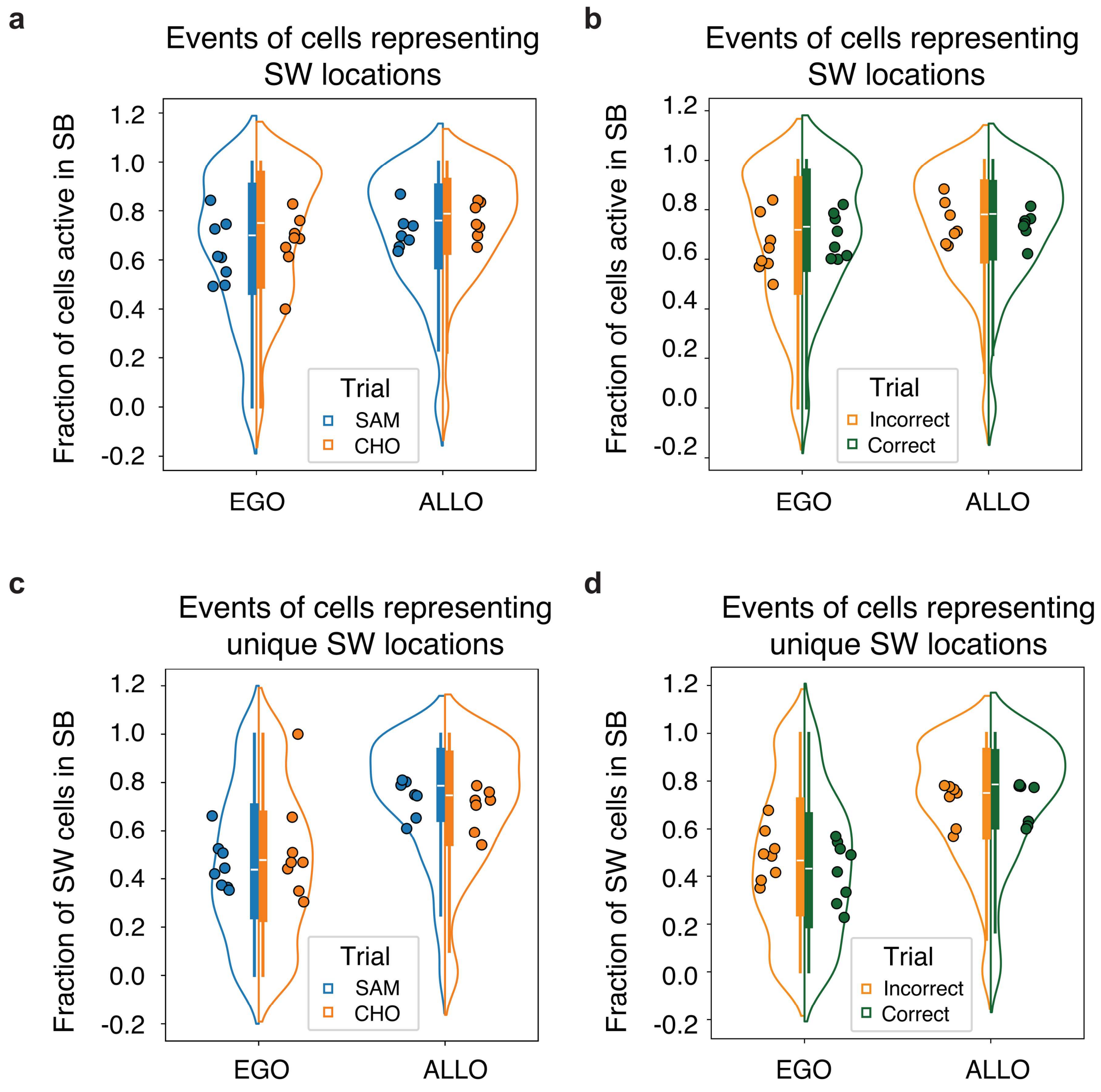

**Figure S14**: a) Nonlocal instances of cell activity mapping to one or more sandwell (SW) locations in egocentric and allocentric animals divided by sample (SAM) or choice (CHO) trials. Mixed Linear Model Regression: Ego/Allo $z=-3.971$ $P<0.001$; SAM/CHO $z=-3.268$ $p=0.001$. b) Same as (a) divided by correct and incorrect trials. Mixed Linear Model Regression: Ego/Allo $z=-3.712$ $P<0.001$; Correct/Incorrect $z=-0.751$ $p=0.453$. c) Nonlocal instances of cell activity mapping a single SW in egocentric and allocentric animals divided by sample (SAM) or choice (CHO) trials. Mixed Linear Model Regression: Ego/Allo $z=-15.473$ $P<0.001$; SAM/CHO $z=-1.430$ $p=0.153$. d) Same as (c) divided by correct and incorrect trials. Mixed Linear Model Regression: Ego/Allo $z=20.556$ $P<0.001$; Correct/Incorrect $z=0.819$ $p=0.413$.

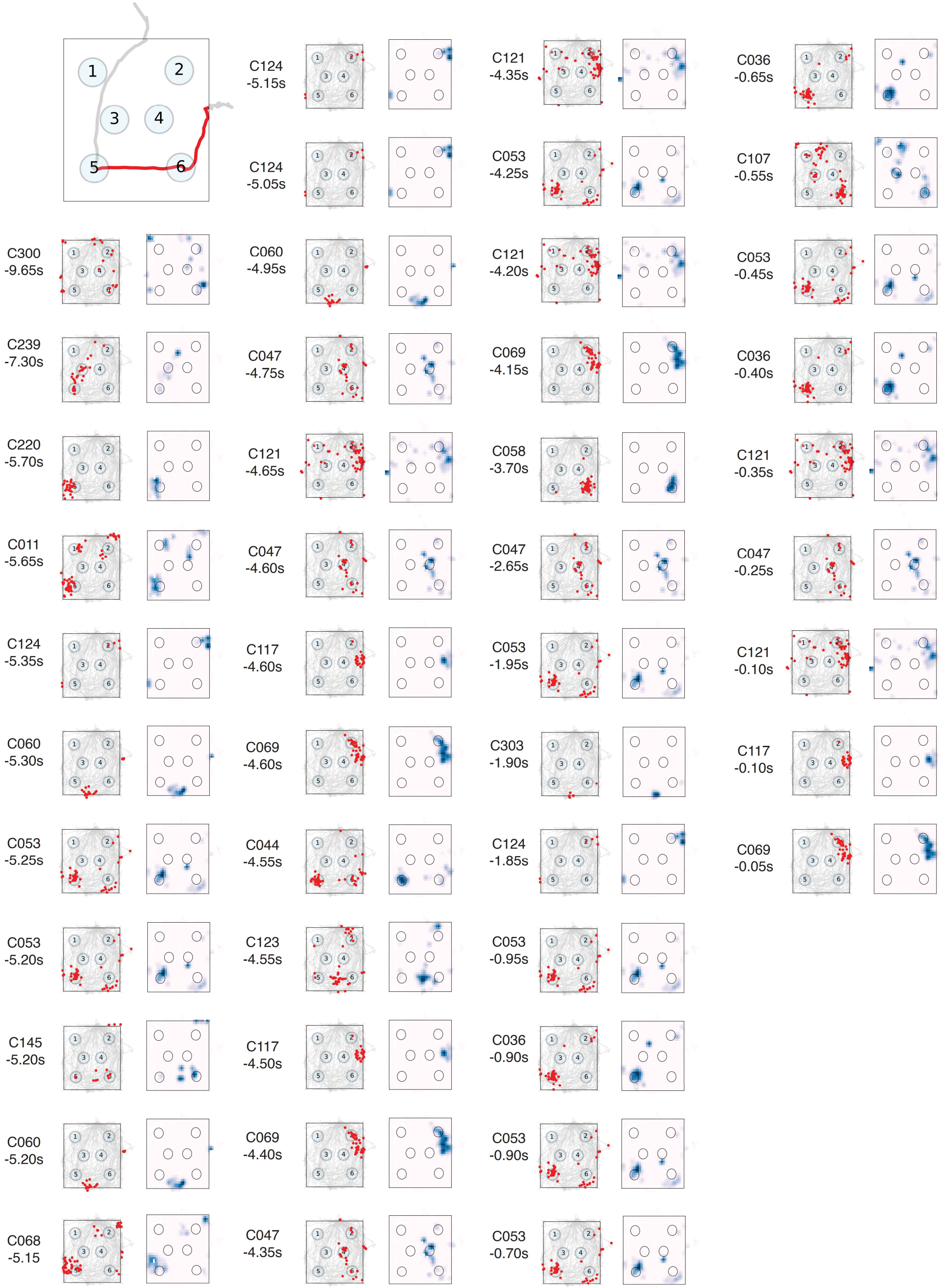

#### Figure S15 (part 1)

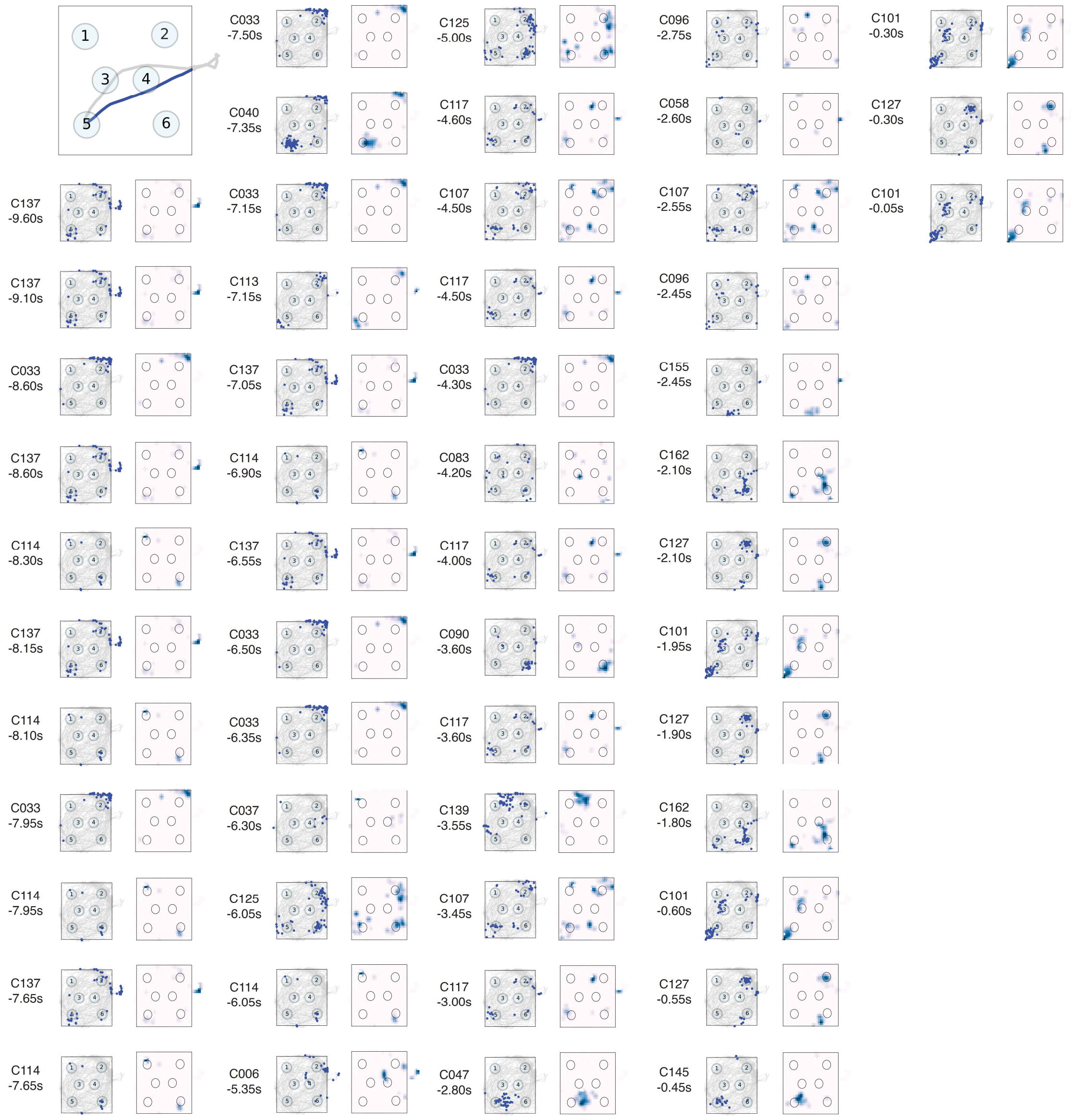

**Figure S15 (part 2)**. Full sequence of cells active in the startbox for trials displayed in Fig 3c. For each activity instance, we have displayed, from left to right, cell ID and time of calcium event onset (negative values indicate times before the trial start), Event map with events as dots (blue or red for egocentric or allocentric tasks, respectively) superimposed to the explored space in gray, and Density map in pink-to-blue. Events are locations where the cell is active while the animal is in the arena in the corresponding session. First panel, actual trajectory run by the animal in color.

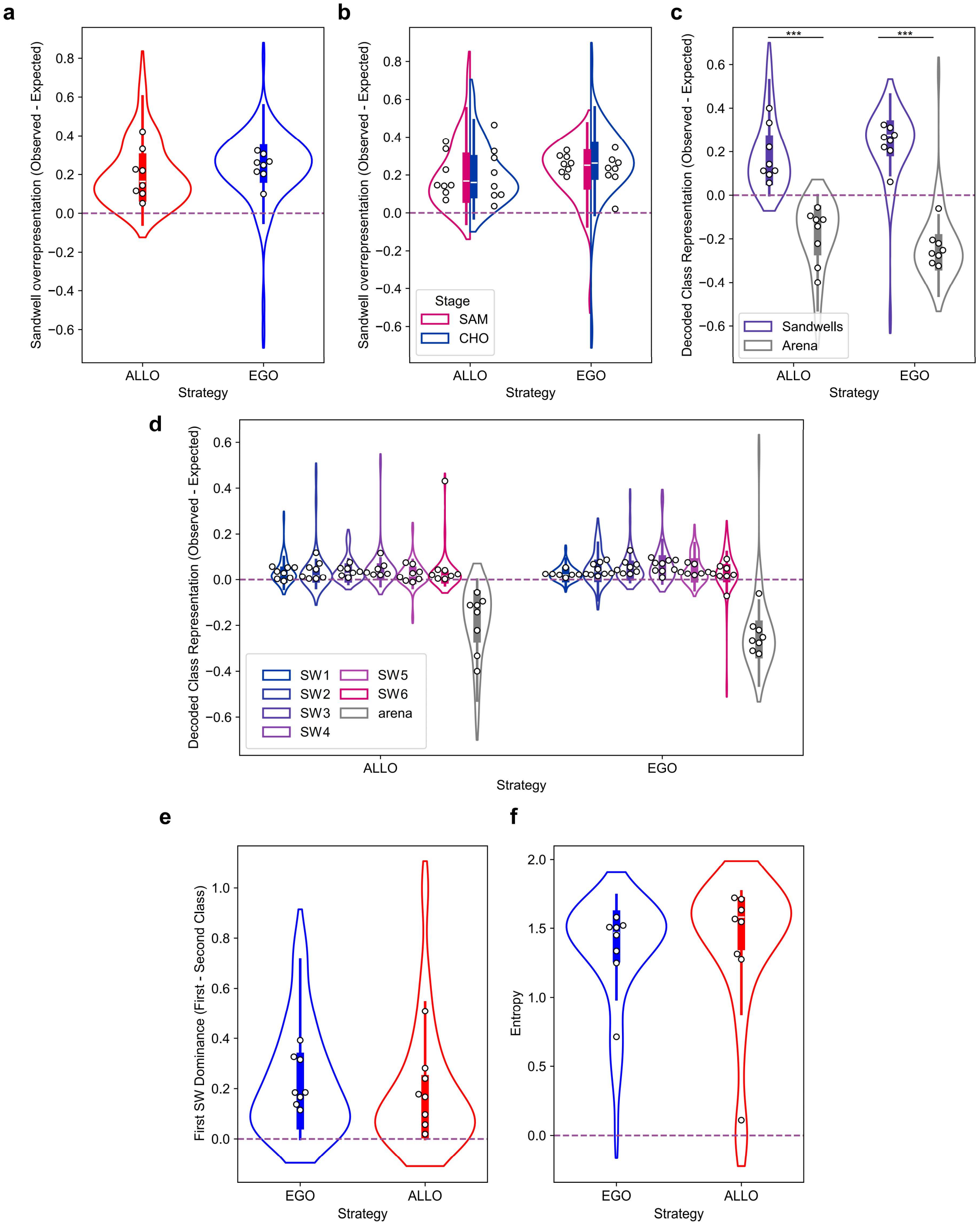

**Figure S16**: A Multinomial Logistic Regression Decoder was trained on the data from the Exploration phase and run on the 10s in the startbox before trial starts. a) Sandwell overrepresentation (difference between the observed decoded frequency of sandwells and the expected frequency) in the startbox intervals for animals trained in the egocentric (EGO) or allocentric (ALLO) task. b) same as divided by sample and choice trials. c) Sandwell locations were significantly more represented than other arena locations. LMER Decoded class representation ~ Strategy (Ego/Allo) +

Location + (1|animal), Ego/Allo z=0 P=0.99; Location z=19.56 P<0.0001; ***P<0.0001 Bonferroni-adjusted pairwise comparison. d) Decoded Class Representation (Observed - Expected) for all sandwells and non-sandwell locations. e) First sandwell dominance (first decoded – second decoded) for allocentric- trained and egocentric-trained animals. f) Entropy of decoded sandwells for allocentric- trained and egocentric-trained animals.

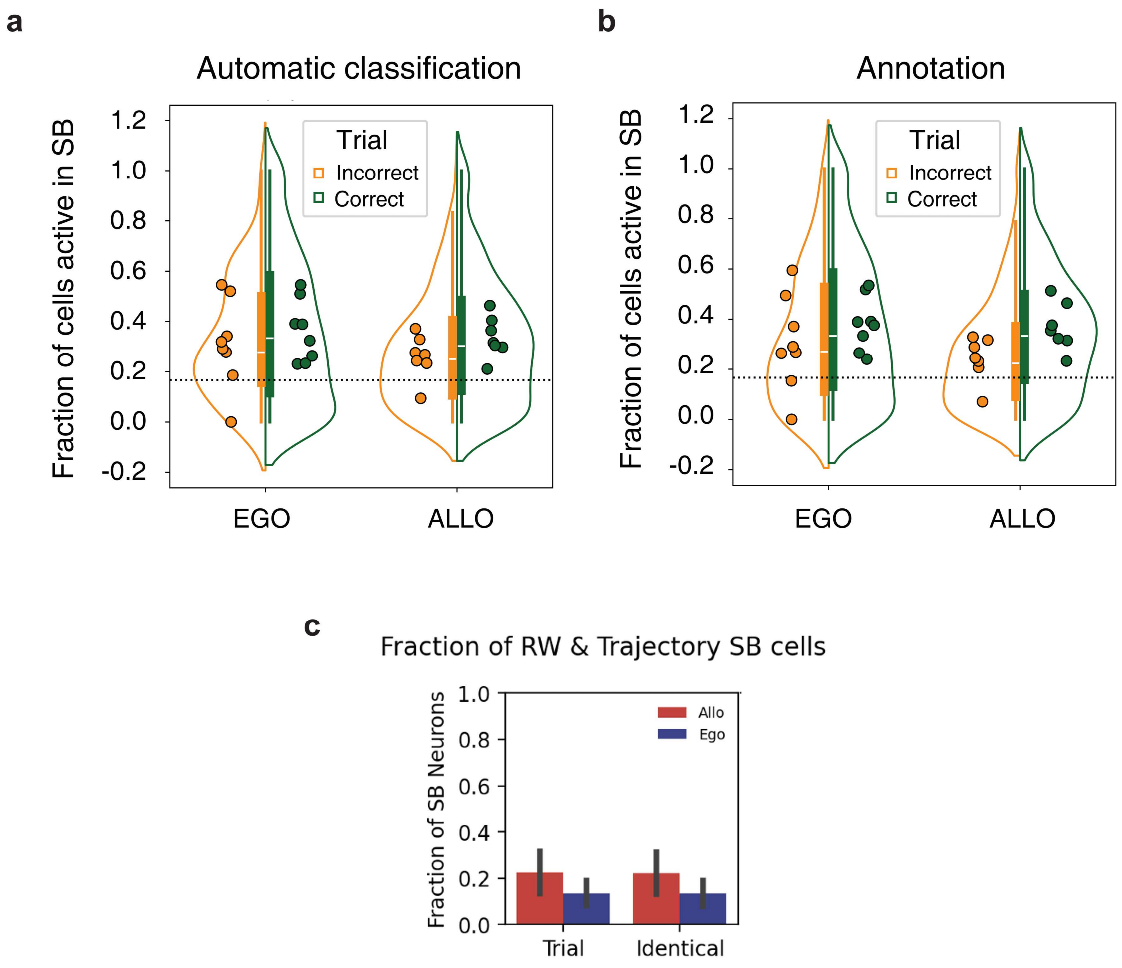

**Figure S17**: Fraction of cells active in the startbox mapping to the correct SW divided by trial number calculated with a fully automatic classification of trials between correct and incorrect (a) or manually annotated trials (b). (c) Fraction of startbox (SB) cells mapping to the correct destination or positions along the trajectory leading to it calculated with a fully automatic algorithm.

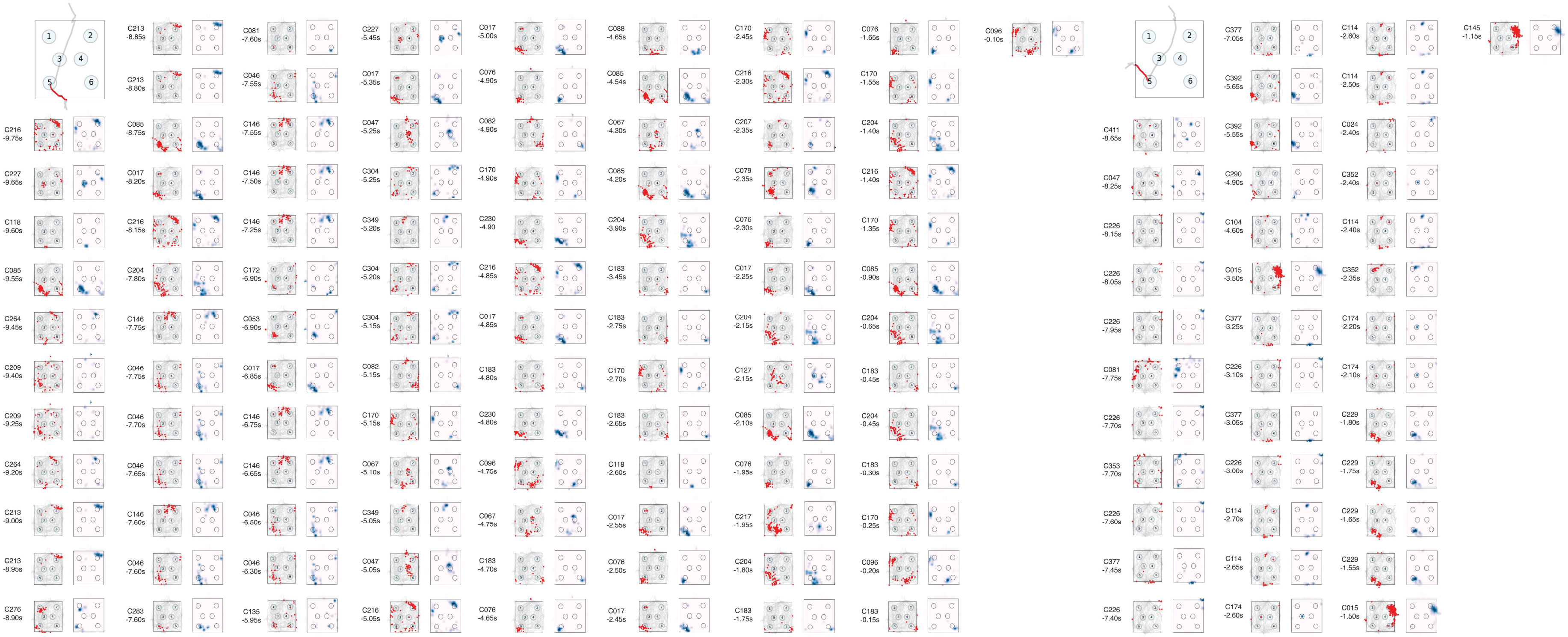

**Figure S18 (part 1)** Further examples of startbox activity from multiple animals before the beginning of the trial during the allocentric task. For each activation instance, we have displayed from left to right, cell ID and time of activation (negative values indicate times before the trial start), Event map with events as dots (red) superimposed to the explored space in gray, Density map in pink-to-blue. First panel, actual trajectory run by the animal in color.

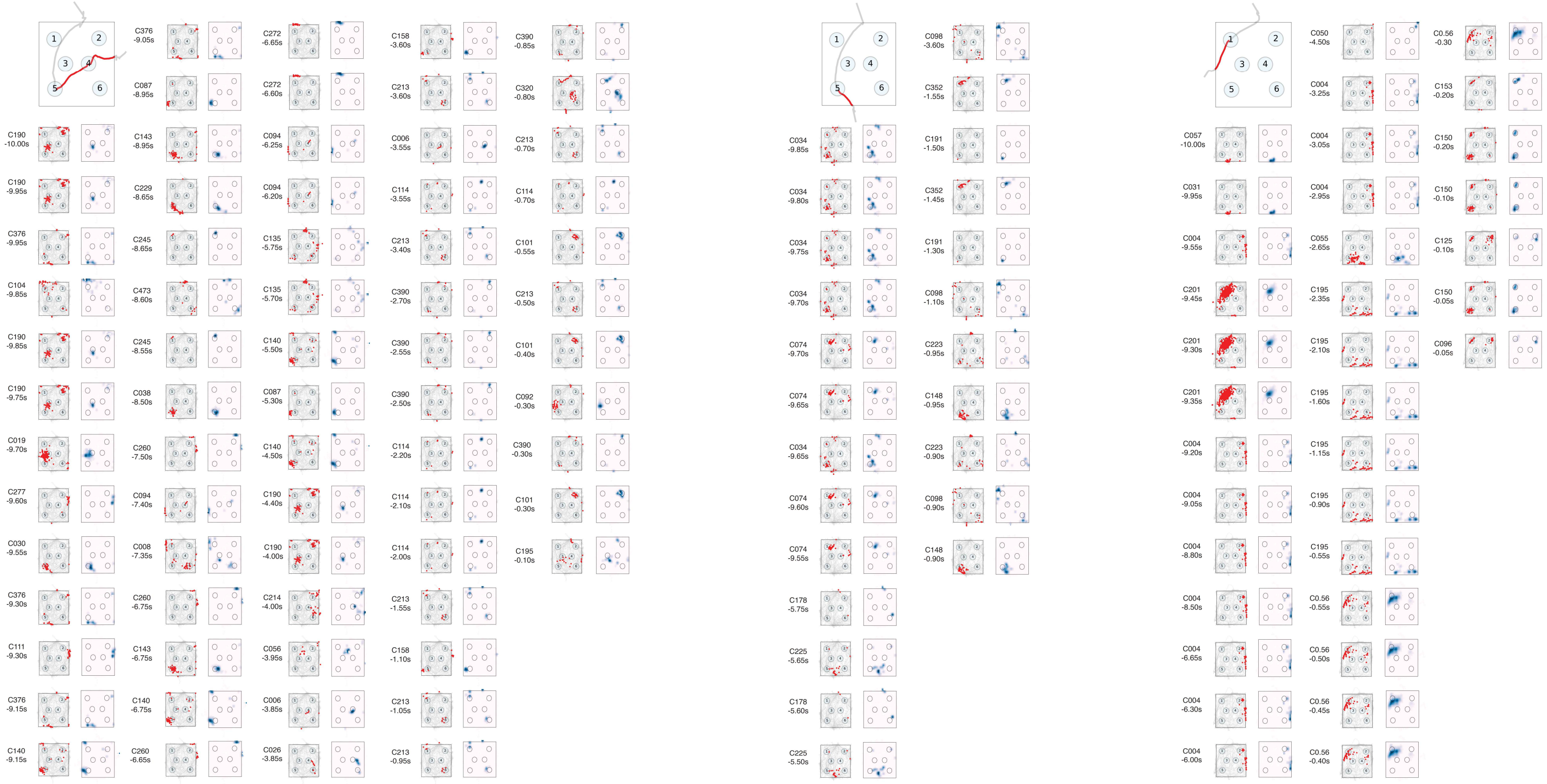

### Figure S18 (part 2)

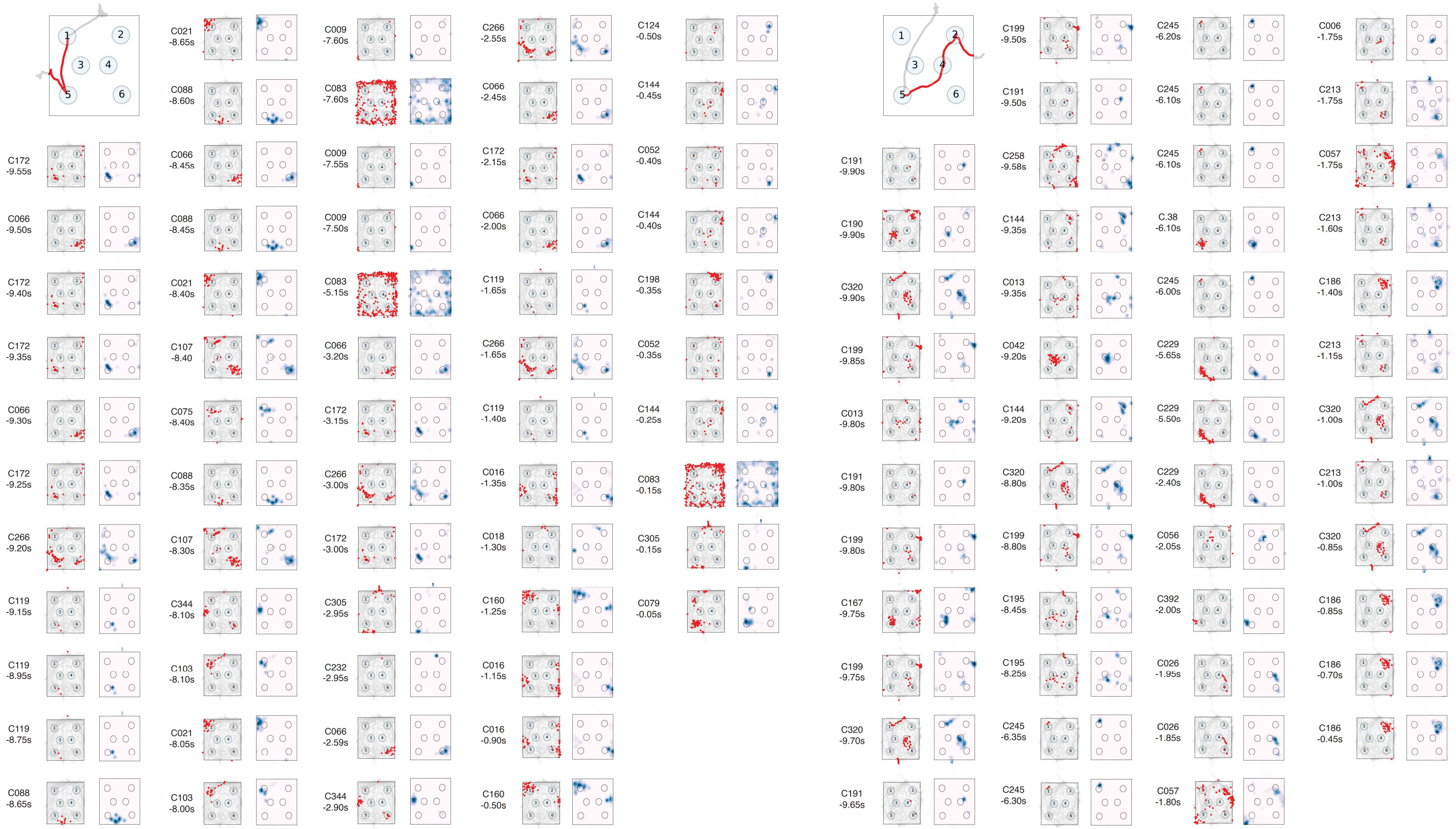

**Figure S19 (part 1)** Further examples of startbox activity from multiple animals before the beginning of the trial during the allocentric task in incorrect trials. For each activation instance, we have displayed from left to right, cell ID and time of activation (negative values indicate times before the trial start), Event map with events as dots (red) superimposed to the explored space in gray, Density map in pink-to-blue. First panel, actual trajectory run by the animal in color.

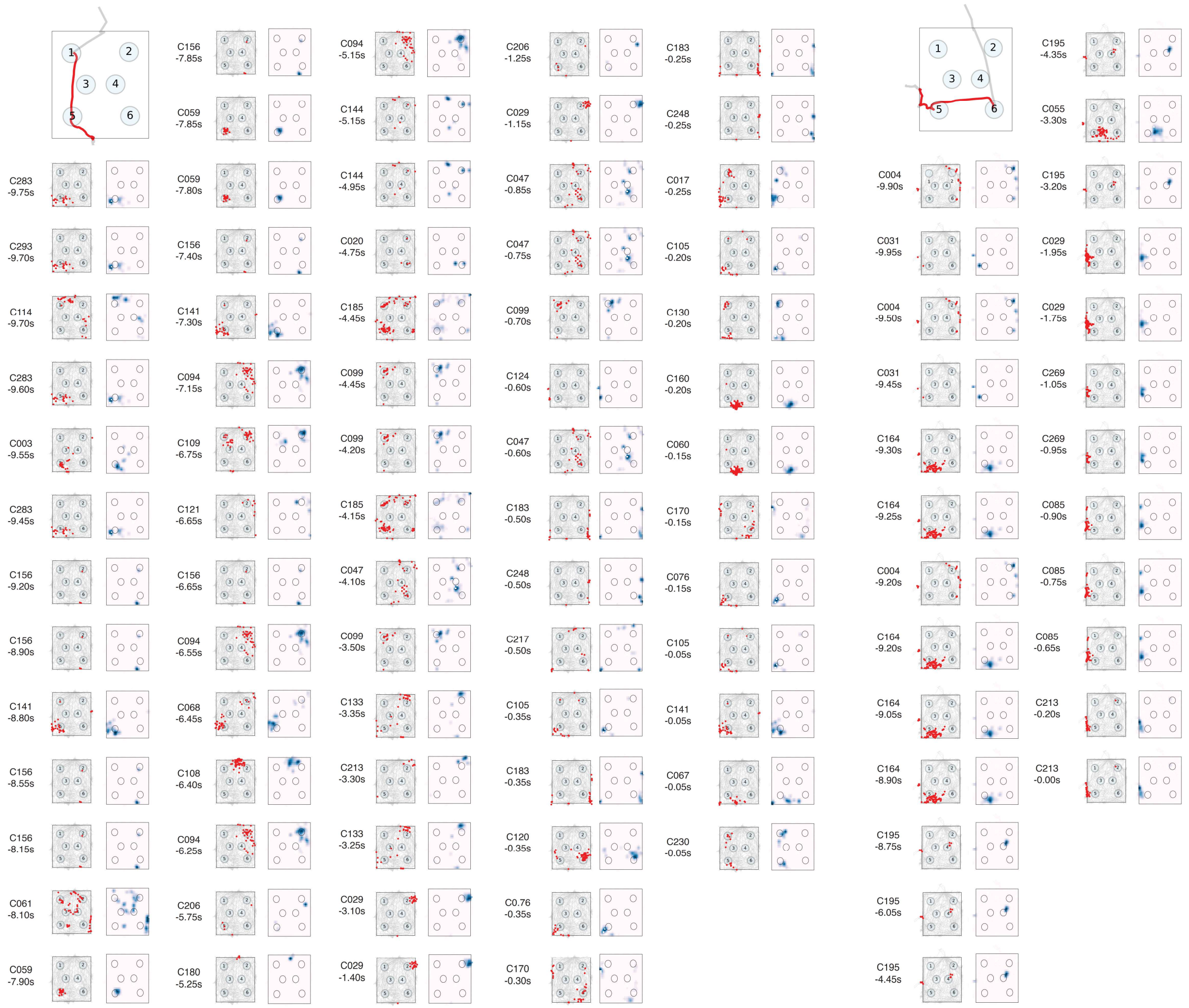

Figure S19 (part 2)

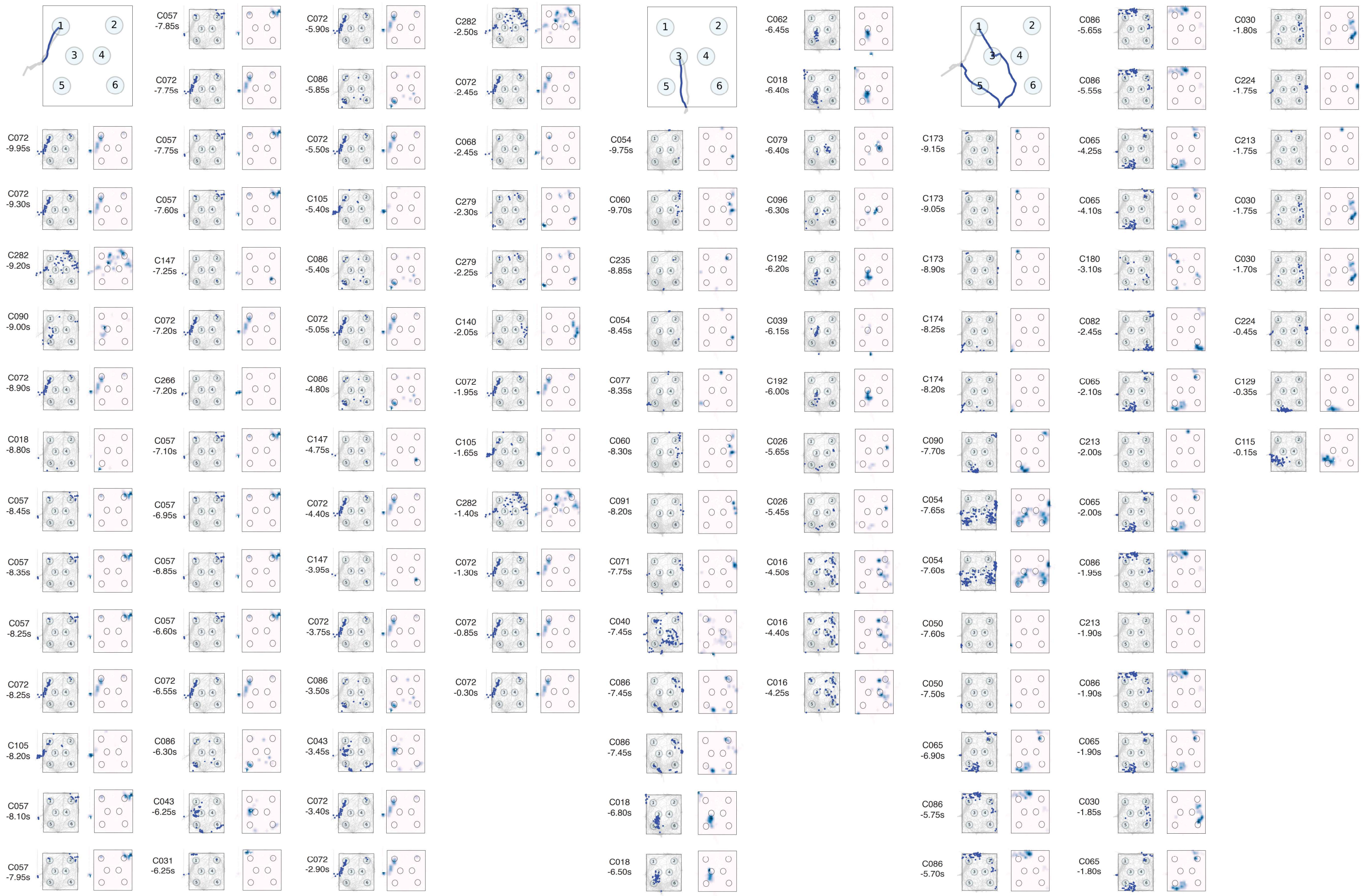

**Figure S20 (part 1)**

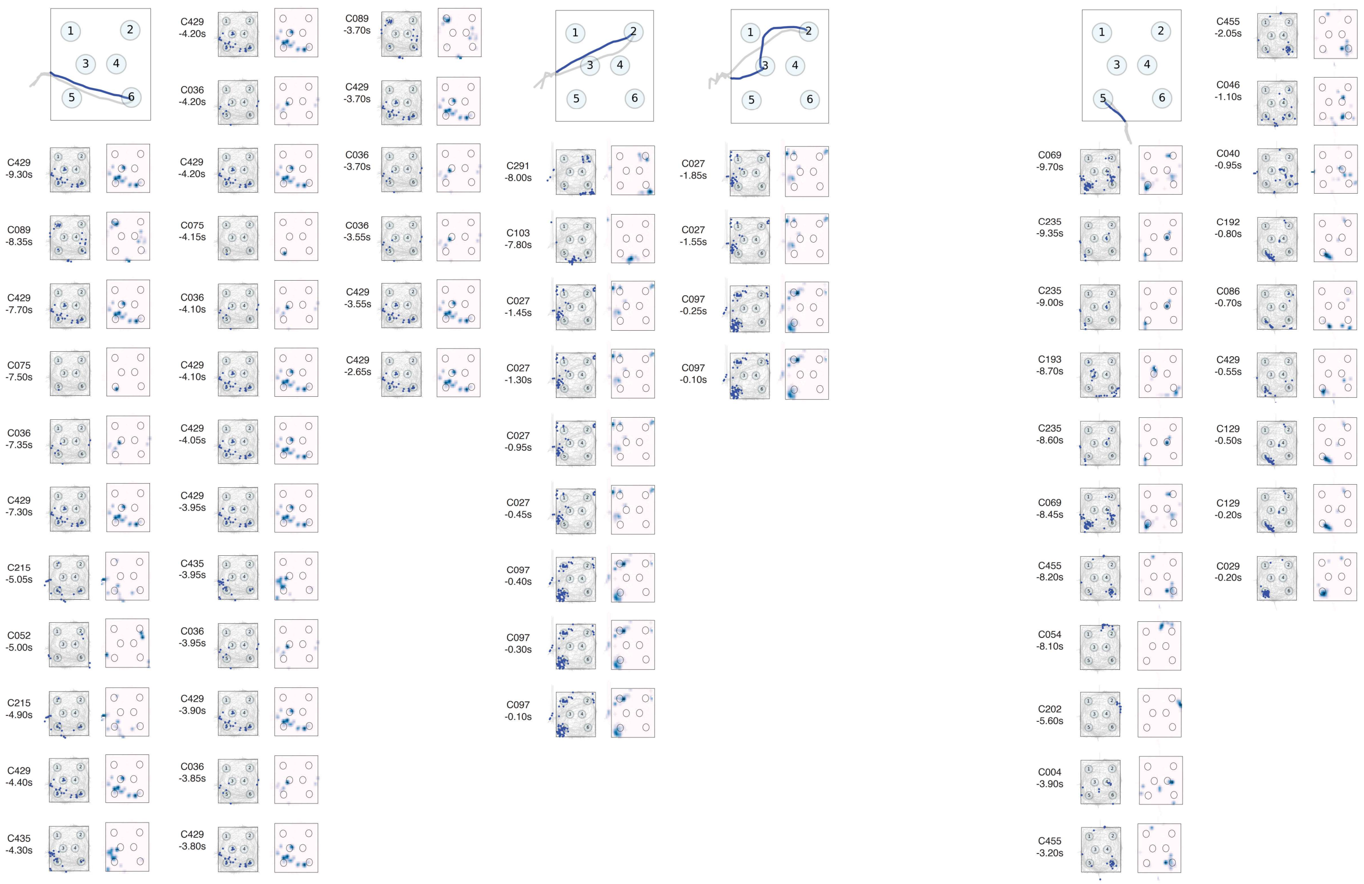

**Figure S20 (part 2)**: Further examples of startbox activity from multiple animals before the beginning of the trial during the egocentric task. For each activation instance, we have displayed from left to right, cell ID and time of activation (negative values indicate times before the trial start), Event map with events as dots (blue) superimposed to the explored space in gray, Density map in pink-to-blue. First panel, actual trajectory run by the animal in color.

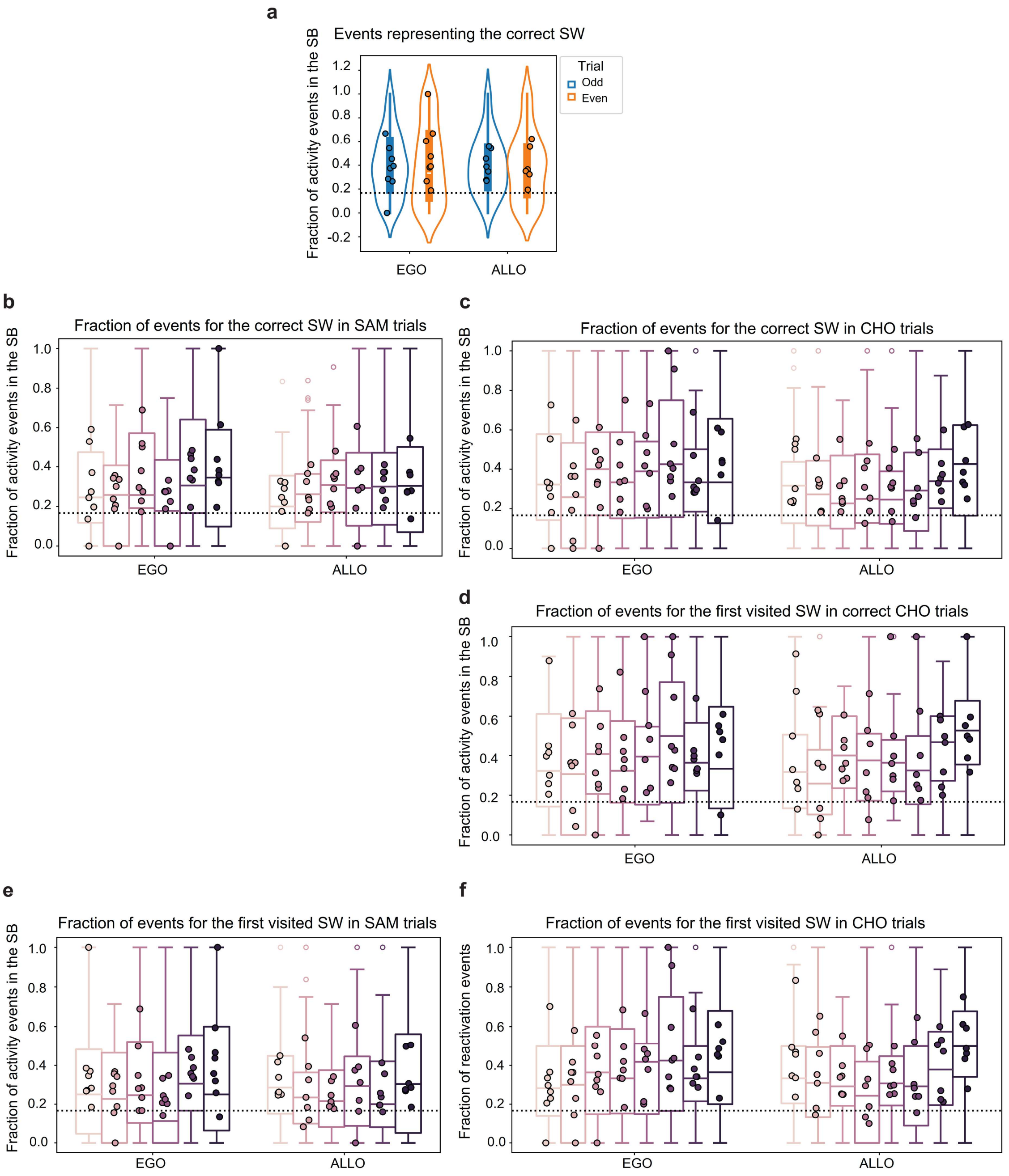

**Figure S21:** a) Fraction of active cell events mapping to the correct SW divided by odd (1,3,5,7) and even trials (2,4,6,8). b) Fraction of active cell events mapping to the correct SW divided by trial number in SAM (b) or CHO trials (c). d) Same as (c) but for correct trials only. (e,f) Fraction of active cell events mapping to the first visited SW divided by trial number in SAM (e) or CHO trials (f).

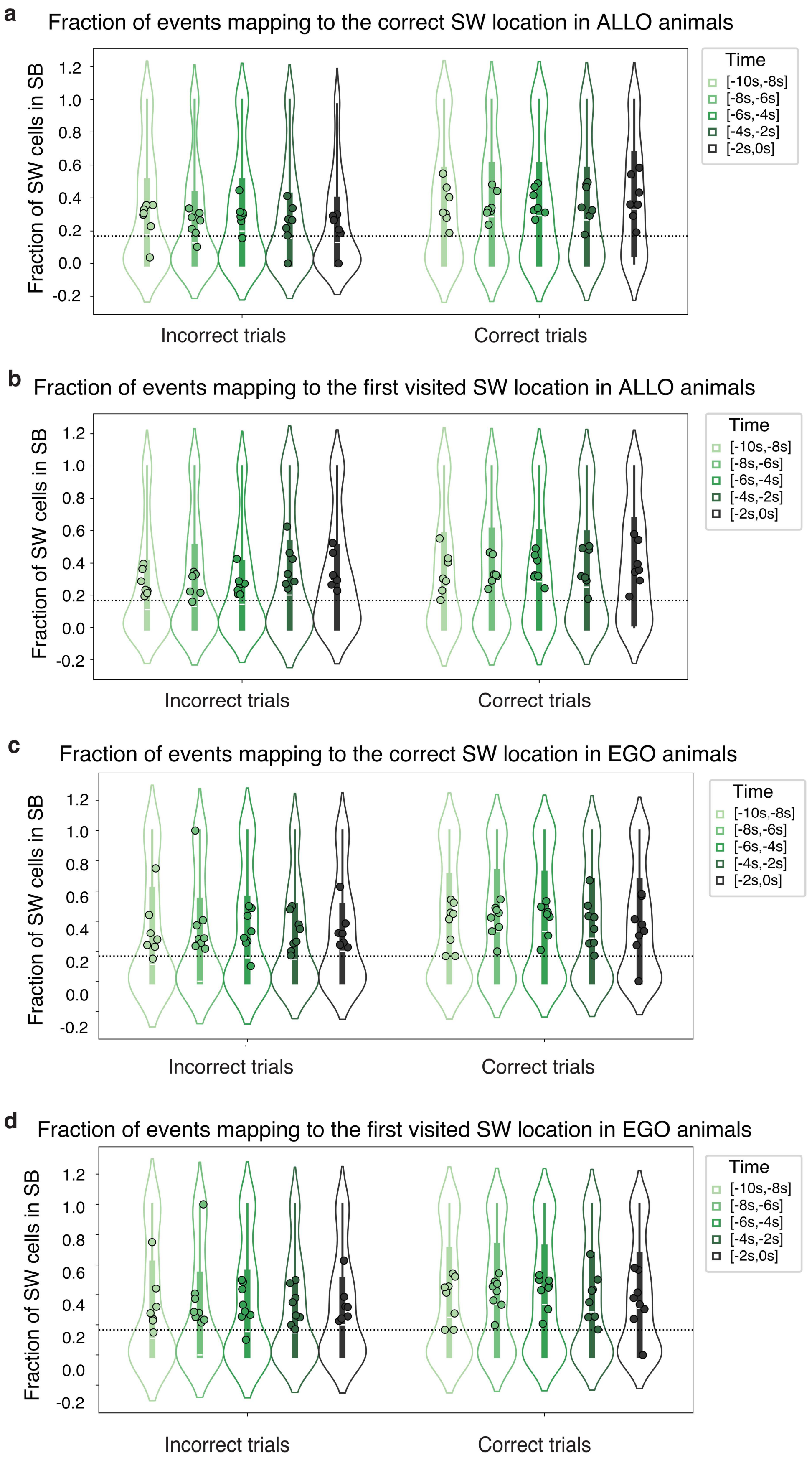
a
Fraction of events mapping to the correct SW location in ALLO animals
Fraction of SW cells in SB
1.2
1.0
0.8
0.6
0.4
0.2
0.0
-0.2
Incorrect trials
Correct trials
Time
[-10s,-8s]
[-8s,-6s]
[-6s,-4s]
[-4s,-2s]
[-2s,0s]
b
Fraction of events mapping to the first visited SW location in ALLO animals
Fraction of SW cells in SB
Incorrect trials
Correct trials
Time
[-10s,-8s]
[-8s,-6s]
[-6s,-4s]
[-4s,-2s]
[-2s,0s]
c
Fraction of events mapping to the correct SW location in EGO animals
Fraction of SW cells in SB
Incorrect trials
Correct trials
Time
[-10s,-8s]
[-8s,-6s]
[-6s,-4s]
[-4s,-2s]
[-2s,0s]
d
Fraction of events mapping to the first visited SW location in EGO animals
Fraction of SW cells in SB
Incorrect trials
Correct trials
Time
[-10s,-8s]
[-8s,-6s]
[-6s,-4s]
[-4s,-2s]
[-2s,0s]

**Figure S22 (previous page):** a) Fraction of active cell events mapping to the correct SW in allocentric animals in correct and incorrect trials as a function of time (Mixed Linear Model Regression correct/incorrect $z=4.883$ $P<0.001$, time $z=-1.517$ $P=0.129$, time*correct/incorrect $z=1.917$ $P=0.55$). b) Fraction of active cell events mapping to the first visited SW in allocentric animals in correct and incorrect trials as a function of time (Mixed Linear Model Regression correct/incorrect $z=1.305$ $P=0.192$, time $z=2.588$ $p=0.011$, time*correct/incorrect $z=-0.932$ $P=0.351$). c) Fraction of active cell events mapping to the correct SW in egocentric animals in correct and incorrect trials as a function of time (Mixed Linear Model Regression correct/incorrect $z=1.366$ $P=0.172$, time $z=-0.925$ $P=0.355$, time*correct/incorrect $z=0.163$ $P=0.87$). d) Fraction of active cell mapping to the first visited SW in egocentric animals in correct and incorrect trials as a function of time (Mixed Linear Model Regression correct/incorrect $z=2.076$ $P=0.038$, time $z=-0.078$ $P=0.938$, time*correct/incorrect $z=-0.271$ $P=0.786$).

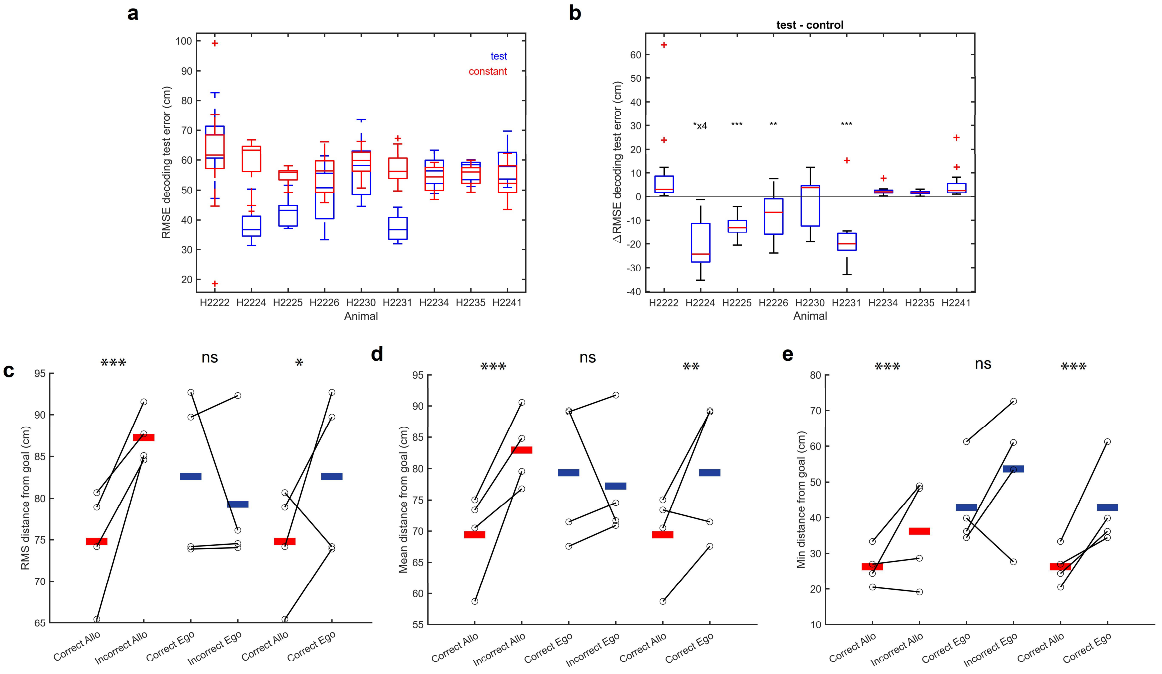

**Figure S23: Bayesian Decoder of Startbox activity.** a) Test vs. constant decoding error in decoders' evaluation (Exploration stage). The decoding error on the test set vs. the error of a constant prediction. For each rat, the overlayed boxplots show the root-mean-squared error (RMSE) of decoding position on the test set of all the sessions using the trained decoder (blue) and a constant prediction at the center of the arena (red). b) Test minus constant decoding error in decoders' evaluation (Exploration stage). The difference between the decoder's test set error and the constant prediction error of all sessions, grouped by rat. Four rats have difference values statistically significantly below 0 (one-tailed Wilcoxon signed-rank test). The other rats are excluded from further decoding analysis due to a lack of significant decoding ability. c-e) Summarized median error values across rats. The paired comparison from all four rats between the median errors over trials for correct vs. incorrect Allo trials (left), correct vs. incorrect Ego trials (middle), and correct Allo vs. correct Ego trials (right). The error measures are the RMS (c), mean (d), and minimum (e). The red and blue horizontal lines denote the mean over the rats color coded by strategy (blue: egocentric, red: allocentric). Statistical comparisons were performed with linear mixed models on individual trials D ~ Correct_Allo v Incorrect_Allo + (1|Animal) for D=RMS $z=3.55$ $P<0.001$; D=Mean $z=3.68$ $P<0.001$; D=Min $z=3.32$ $P<0.001$. D ~ Correct_Ego v Incorrect_Ego + (1|Animal) for D=RMS $z=1.58$ $P=0.13$; D=Mean $z=1.45$ $P=0.15$; D=Min $z=0.73$ $P=0.47$ . D ~ Correct_Allo v Correct_Ego + (1|Animal) for D=RMS $z=2.18$ $P=0.03$; D=Mean $z=3.15$ $P=0.002$; D=Min $z=7.46$ $P<0.001$.

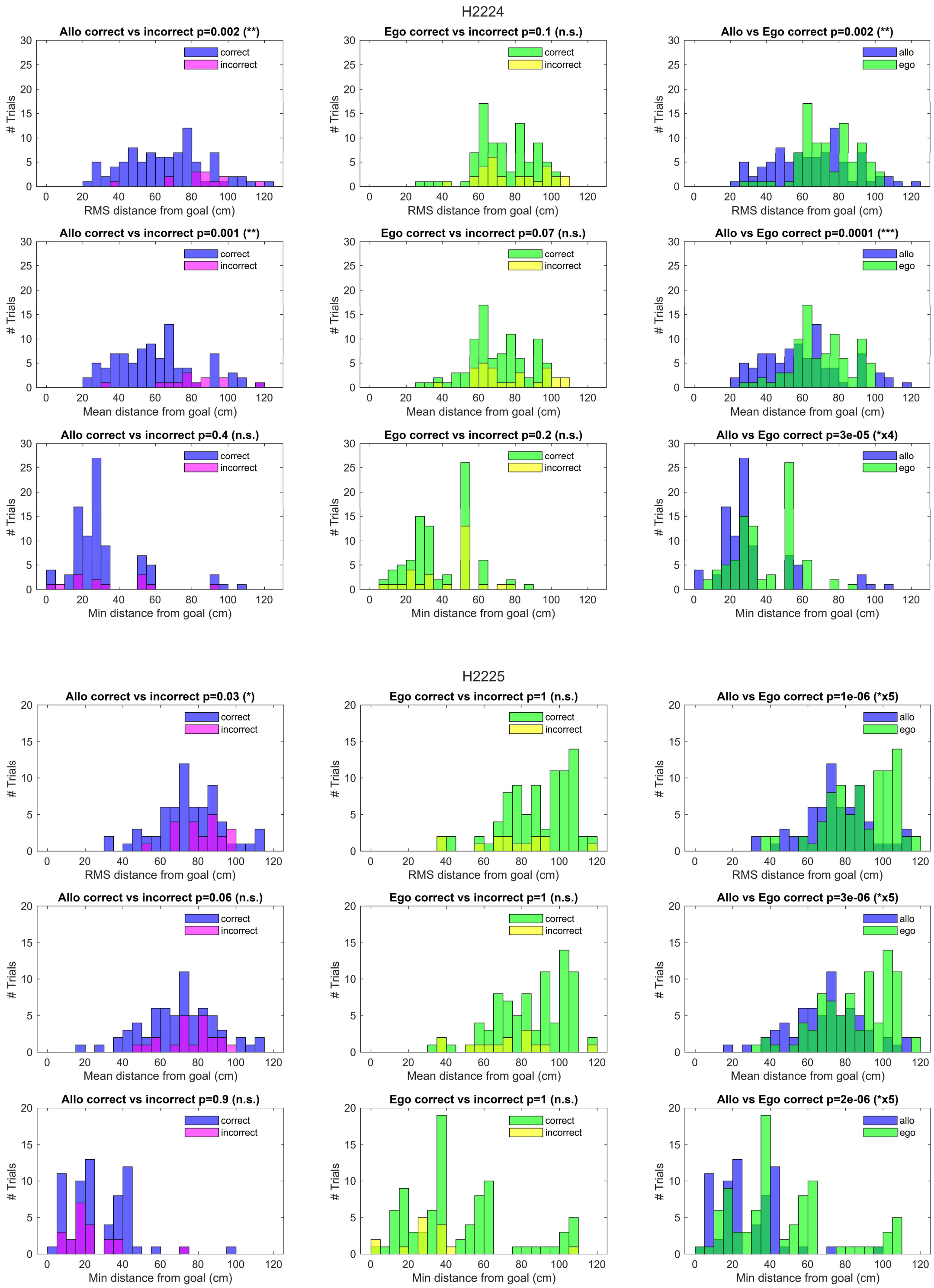

#### Figure S24: Bayesian Decoder (part 1)

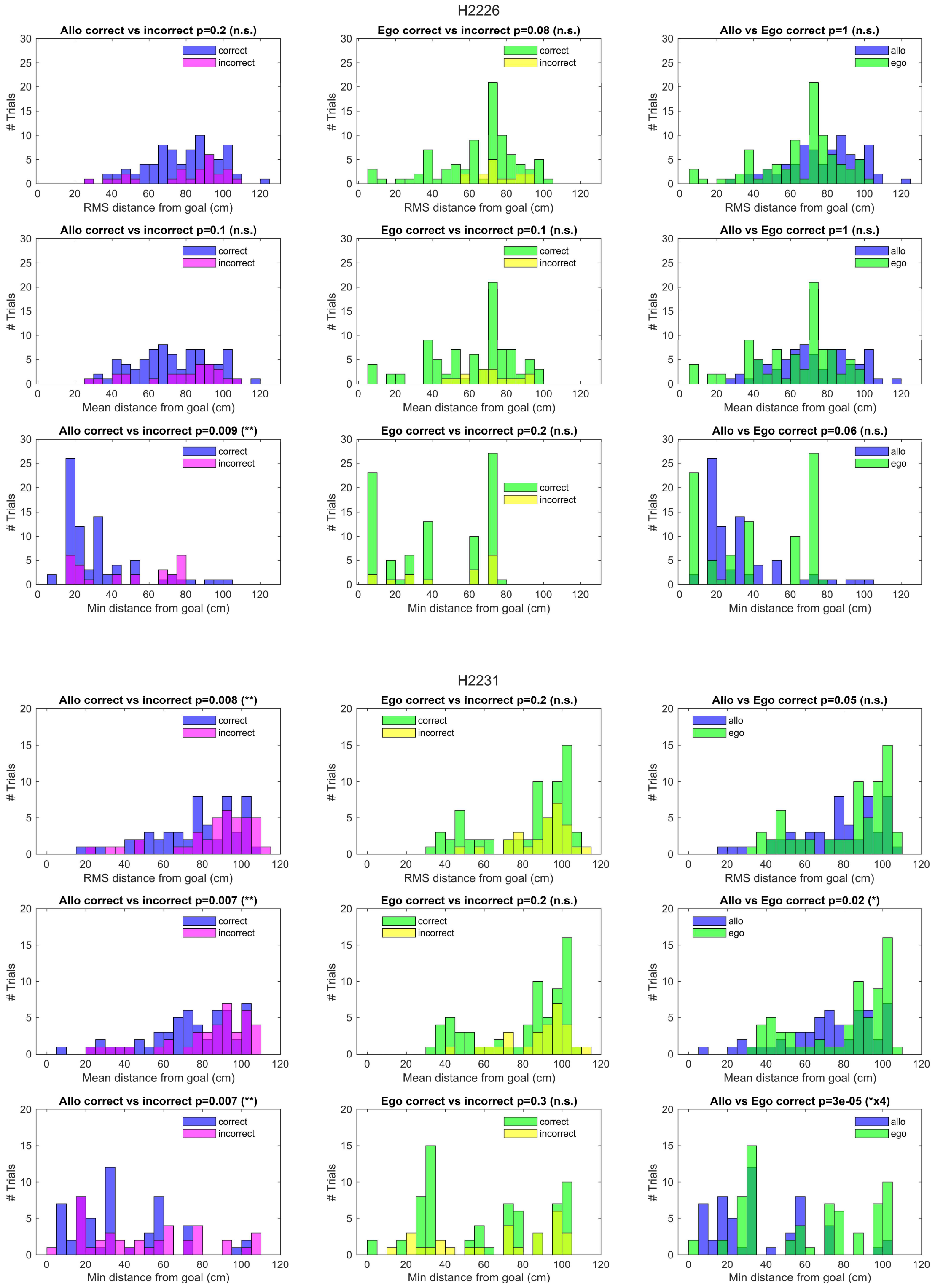

**Figure S24: Bayesian Decoder (part 2).** Goal location decoding error histograms For each rat the figures show overlayed histograms of goal location prediction error of correct vs. incorrect Allo trials (left column), correct vs. incorrect Ego trials (middle column), and correct Allo trials vs. correct Ego trials (right column). The error measure compared is RMS (top row), mean (middle row), and minimum (bottom row). Differences in the medians of the histograms are assessed with a one-tailed Mann-Whitney U test testing for correct < incorrect or Allo < Ego.

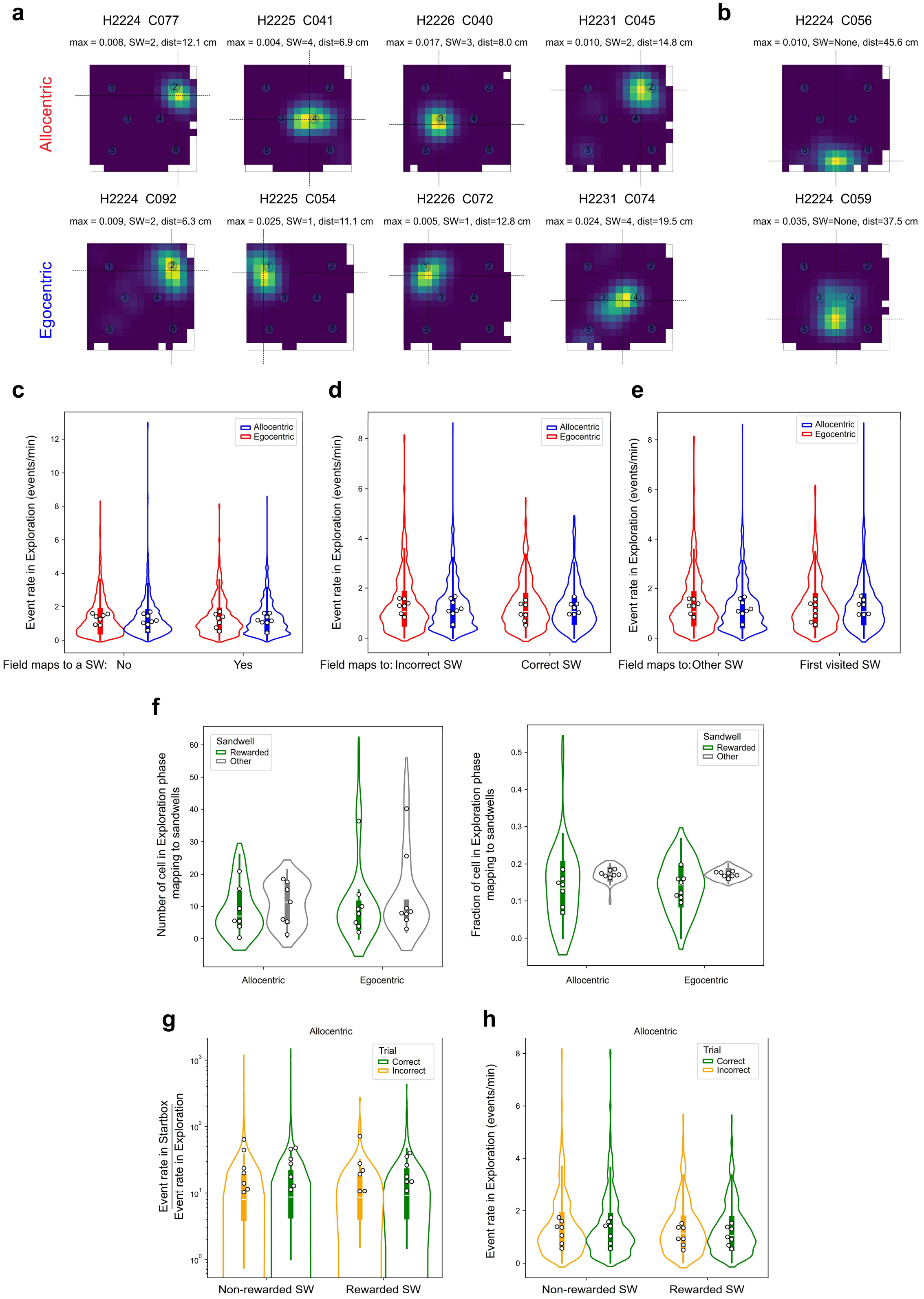
a
b
H2224 C077
max = 0.008, SW=2, dist=12.1 cm
H2225 C041
max = 0.004, SW=4, dist=6.9 cm
H2226 C040
max = 0.017, SW=3, dist=8.0 cm
H2231 C045
max = 0.010, SW=2, dist=14.8 cm
H2224 C056
max = 0.010, SW=None, dist=45.6 cm
Allocentric
H2224 C092
max = 0.009, SW=2, dist=6.3 cm
H2225 C054
max = 0.025, SW=1, dist=11.1 cm
H2226 C072
max = 0.005, SW=1, dist=12.8 cm
H2231 C074
max = 0.024, SW=4, dist=19.5 cm
H2224 C059
max = 0.035, SW=None, dist=37.5 cm
Egocentric
c
d
e
Event rate in Exploration (events/min)
Allocentric
Egocentric
Field maps to a SW: No
Yes
Field maps to: Incorrect SW
Correct SW
Field maps to:Other SW
First visited SW
f
Number of cell in Exploration phase mapping to sandwells
Fraction of cell in Exploration phase mapping to sandwells
Sandwell
Rewarded
Other
Allocentric
Egocentric
g
h
Allocentric
Event rate in Startbox
Event rate in Exploration
Event rate in Exploration (events/min)
Trial
Correct
Incorrect
Non-rewarded SW
Rewarded SW

**Figure S25 (previous page): Event frequency during Exploratory phase.** a,b) Cells with fields mapping to sandwells were determined automatically with the average centroid position. a) Representative place maps for cells mapping to a sandwell (SW) by the algorithm for Allocentric and Egocentric datasets. The maximum of the ratemap, the identity of the classified SW and the centroid distance are indicated above. Crossed lines identify the centroid coordinates. b) Representative maps for cells that were not classified as mapping to any sandwell and classified as 'Other'. c) Event rate (events per minute) during Exploration phase for cells that were active in the 10s time periods before trial starts identified with fields mapping to a Sandwell (Yes) or Other (No) showing no difference in baseline firing rate. LMER Event rate ~ Maps_to + Ego/Allo + (1|animal, cell). Map_to z=0.66 P=0.51; Ego/Allo z=0.85 P=0.40 . d) Event rate (events per minute) during Exploration phase for cells that were active in the 10s time periods before trial starts identified with fields mapping to the correct Sandwell (rewarded SW) or to an incorrect one (incorrect SW) showing no difference in baseline firing rate. LMER Event rate ~ SW + Ego/Allo + (1|animal, cell). SW z=0.52 P=0.61; Ego/Allo z=0.46 P=0.65 . e) Event rate (events per minute) during Exploration phase for cells that were active in the 10s time periods before trial starts identified with fields mapping to the first visited Sandwell (First visited SW) or to one of the other 5 (Other SW) showing no difference in baseline firing rate. LMER Event rate ~ SW + Ego/Allo + (1|animal, cell). SW z=0.56 P=0.58; Ego/Allo z=0.46 P=0.64 . In d,e) we removed cells mapping to Other and considered only whose field mapped to a sandwell. f) Number (left) and Fraction (right) of cells active in the Exploration phase with fields mapping to the rewarded well or one of the other 5 wells. Note the values for other wells are the average of cells per sandwell. LMER N cells ~ SW + Ego/Allo + (1|animal) SW z=1.99 P=0.046, Ego/Allo z=1.42 P=0.16 . g) Ratio of activity per minute in the startbox to activity in the Exploration phase per minute in allocentric animals, if cells mapped to the rewarded sandwell or a non-rewarded one, divided between correct and incorrect trials. Cells with zero activity in the startbox were excluded from this analysis. LMER Normalised_activity ~ SW + Trial_type + (1|animal, cell) SW z=0.23 P=0.82, Trial_type z=2.5 P=0.50. h) Ratio of activity per minute in the startbox to activity in the Exploration phase per minute in allocentric animals, if cells mapped to the rewarded sandwell or a non-rewarded one, divided between correct and incorrect trials. Cells with zero activity in the startbox were excluded from this analysis. LMER Normalised_activity ~ SW + Trial_type + (1|animal, cell) SW z=0.23 P=0.82, Trial_type z=2.5 P=0.50. h) For allocentric animals, Event rate (events per minute) during Exploration phase for cells that were active in the 10s time periods before trial starts identified with fields mapping to the correct Sandwell (rewarded SW) or to an incorrect one (non-rewarded SW), divided between correct and incorrect trials. LMER Event rate ~ SW + Trial_type + (1|animal, cell). SW z=2.11 P=0.034; Ego/Allo z=0.08 P=0.94 .

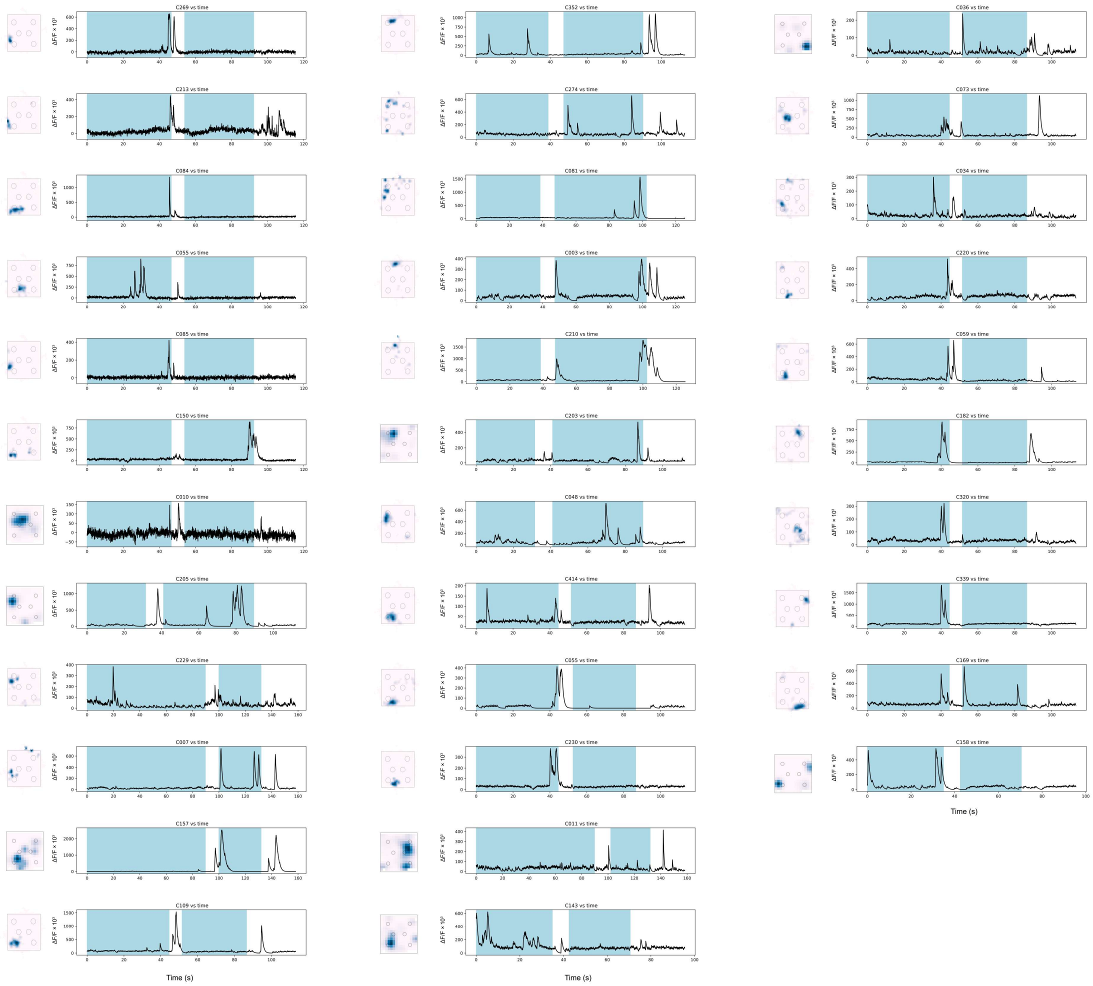

**Figure S26: Calcium traces for cells mapping to future and alternative spaces.** ΔF/F traces (black line) for representative trials for animals performing in the allocentric task. Light blue shades are part of Sample or Choice stages before the beginning of trials when animals were in the startboxes. Each trace represents a pair of trials as described in Figure 1 and Methods. Besides each trace is the place map of the corresponding cell during Exploration in pink-to-blue color map. Note that cells display robust events both in the startbox and at various points in the execution of the trial (white) to the correct sandwell. Some trials where robust events corresponding to alternative locations – not visited in the upcoming trial – are displayed on the rightmost column.

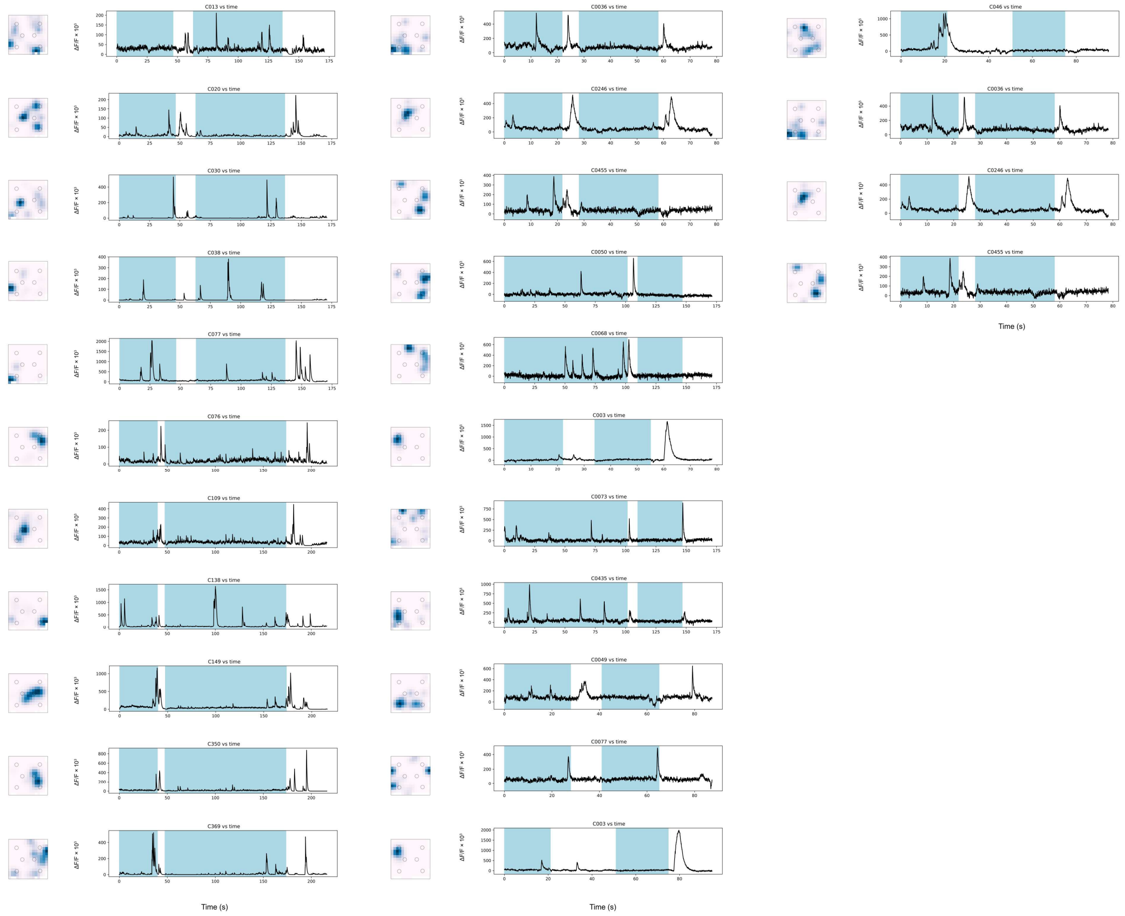

**Figure S27: Calcium traces for cells mapping to future and alternative spaces.** ΔF/F traces (black line) for representative trials for animals performing in the egocentric task. Light blue shades are part of Sample or Choice stages before the beginning of trials when animals were in the startboxes. Each trace represents a pair of trials as described in Figure 1 and Methods. Besides each trace is the place map of the corresponding cell during Exploration in pink-to-blue color map. Note that cells display robust events both in the startbox and at various points in the execution of the trial (white) to the correct sandwell.

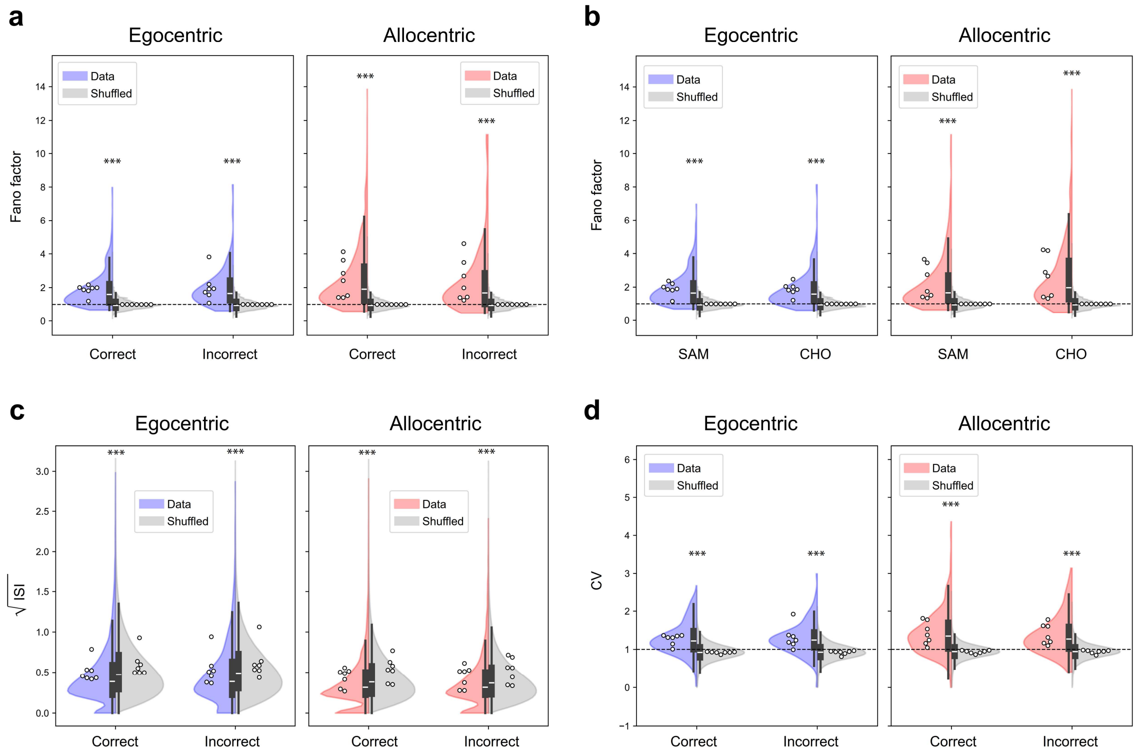

**Figure S28 Temporal distribution of Startbox events.** a) Fano factor distribution calculated for all cells active in the Startbox for Egocentric (blue) and Allocentric (red) trained animals divided by correct and incorrect trials. Fano factor for shuffled data is represented in grey for matching trials. Here and throughout the Figure, violin plots and histogram represent the distribution of trial values, and dots are animal averages. LMER Fano ~ Ego/Allo*Correct/Incorrect + Data/Shuffle + (1|animal); Ego/Allo z= -1.851 P=0.064, Data/Shuffle z= - 135.8 P<0.001; Ego/Allo*Correct/Incorrect z=1.119 P=0.263. b) Same as (a) dividing data between Sample and Choice trials. LMER Fano ~ Ego/Allo*SAM/CHO + Data/Shuffle + (1|animal); Ego/Allo z= -1.349 P=0.177, Data/Shuffle z= - 135.8 P<0.001; Ego/Allo*SAM/CHO z=0.360 P=0.719. Data was significantly different from shuffle: Egocentric z=-0.879 P<0.0001, Allocentric z=-1.475 P<0.0001. c) Inter spike interval (ISI) distribution as in (a). Data are presented as root square of ISI. LMER ISI (mean) ~ Ego/Allo*Correct/Incorrect + Data/Shuffle + (1|animal); Ego/Allo z= 134.041 P<0.0001, Data/Shuffle z= 6.756 P=0.088; Ego/Allo*Correct/Incorrect z=-18.655 P<0.0001. Data vs Shuffle Egocentric z=0.184 P<0.0001, Allocentric z=0.135 P<0.0001. d) CV distribution as in (a). LMER CV ~ Ego/Allo*Correct/Incorrect + Data/Shuffle + (1|animal); Ego/Allo z= -36.831 P<0.0001, Data/Shuffle z= -64.733 P<0.0001; Ego/Allo*Correct/Incorrect z=2.649 P=0.008. Data vs Shuffle Egocentric z=-0.36 P<0.0001, Allocentric z=-0.48 P<0.0001.

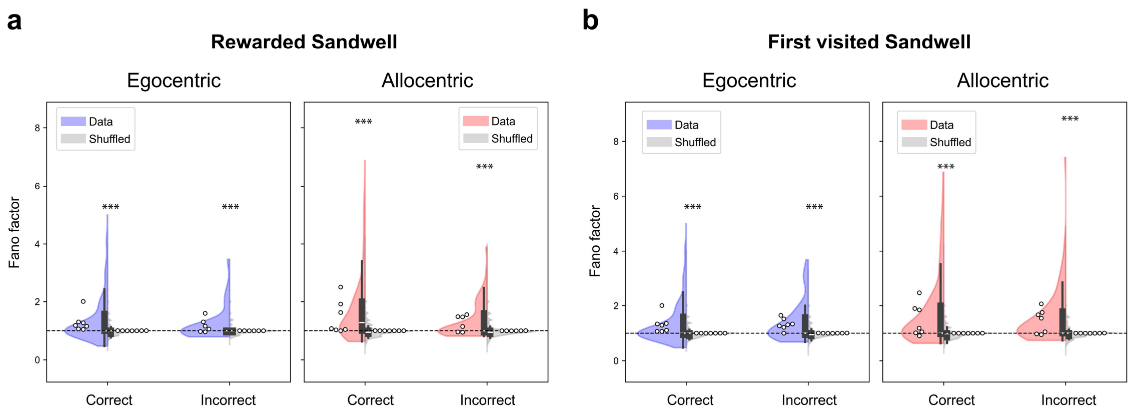

**Figure 29** a) Fano factor distribution calculated for cells active in the startbox representing the rewarded sandwell for Egocentric (blue) and Allocentric (red) trained animals divided by correct and incorrect trials. Fano factor for shuffled data is represented in grey for matching trials. LMER Fano ~ Ego/Allo*Correct/Incorrect + Data/Shuffle + (1|animal); Ego/Allo z= -0.177 P=0.86, Data/Shuffle z= - 44.56 P<0.001; Ego/Allo*Correct/Incorrect z=0.258 P=0.763. Data vs Shuffle Egocentric z=-0.329 P<0.0001, Allocentric z=-0.609 P<0.0001. b) Fano factor distribution calculated for cells active in the startbox representing the first visited sandwell for Egocentric (blue) and Allocentric (red) trained animals divided by correct and incorrect trials. Fano factor for shuffled data is represented in grey for matching trials. LMER Fano ~ Ego/Allo*Correct/Incorrect + Data/Shuffle + (1|animal); Ego/Allo z= -0.452 P=0.651, Data/Shuffle z= - 52.703 P<0.001; Ego/Allo*Correct/Incorrect z=0.963 P=0.349. Data vs Shuffle Egocentric z=-0.368 P<0.0001, Allocentric z=-0.687 P<0.0001.

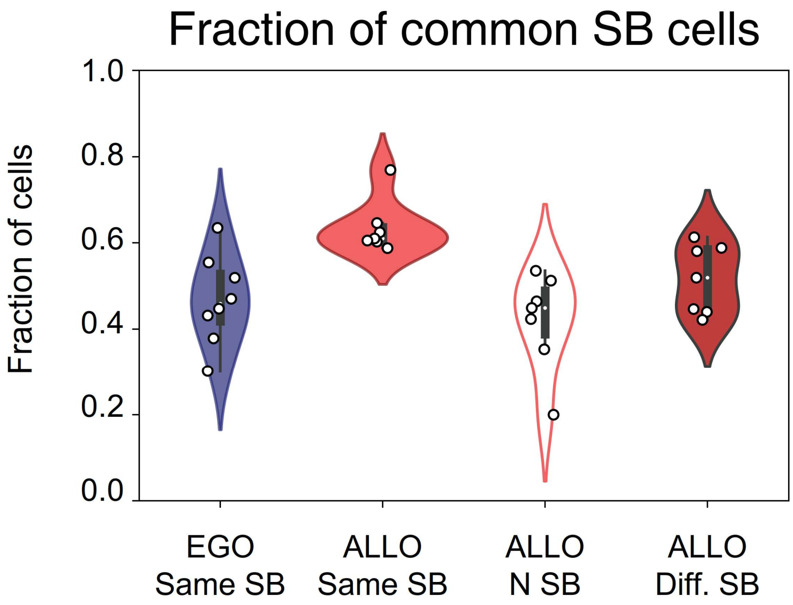

**Figure S30** Common cells between trials in the 10s before trial begin including trials starting from N Goal Box

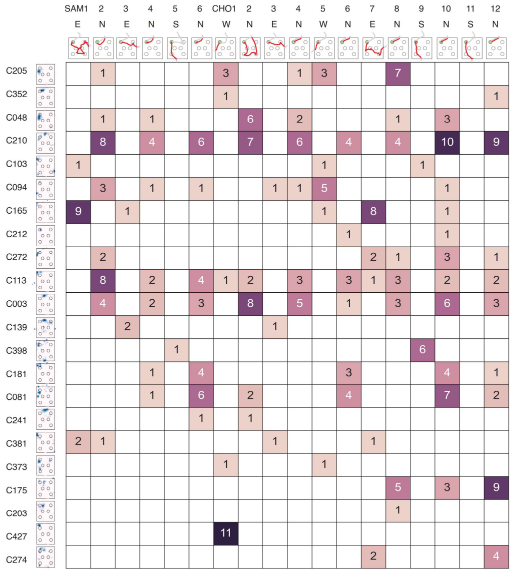
SAM1 2 3 4 5 6 CHO1 2 3 4 5 6 7 8 9 10 11 12
E N E N S N W N E N W N E N S N S N
C205 C352 C048 C210 C103 C094 C165 C212 C272 C113 C003 C139 C398 C181 C081 C241 C381 C373 C175 C203 C427 C274

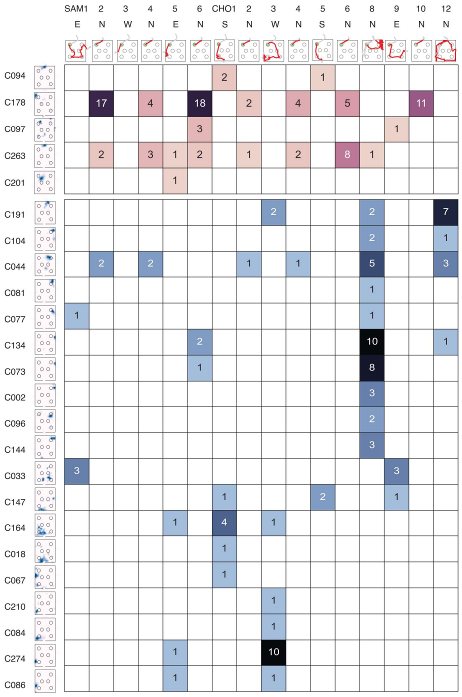
SAM1 2 3 4 5 6 CHO1 2 3 4 5 6 8 9 10 12
E N W N E N S N W N S N N E N N
C094 C178 C097 C263 C201 C191 C104 C044 C081 C077 C134 C073 C002 C096 C144 C033 C147 C164 C018 C067 C210 C084 C274 C086

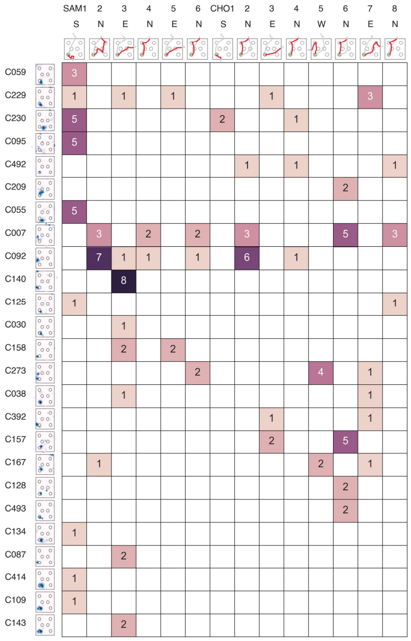
SAM1 2 3 4 5 6 CHO1 2 3 4 5 6 7 8
S N E N E N S N E N W N E N
C059 C229 C230 C095 C492 C209 C055 C007 C092 C140 C125 C030 C158 C273 C038 C392 C157 C167 C128 C493 C134 C087 C414 C109 C143

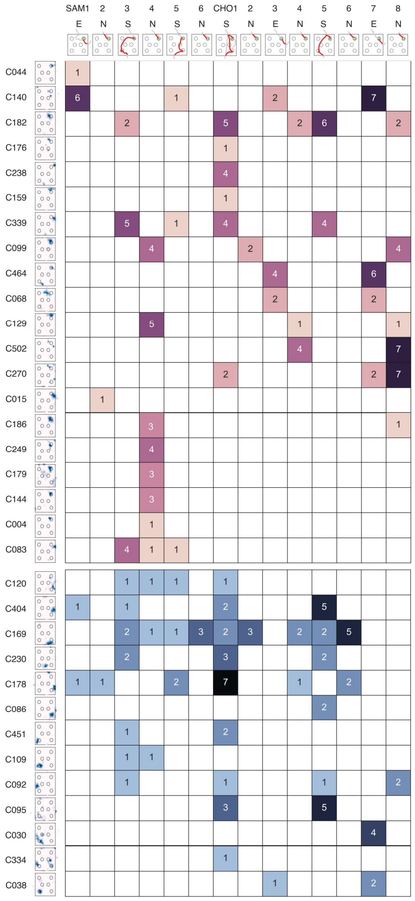
SAM1 2 3 4 5 6 CHO1 2 3 4 5 6 7 8
E N S N S N S N E N S N E N
C044 C140 C182 C176 C238 C159 C339 C099 C464 C068 C129 C502 C270 C015 C186 C249 C179 C144 C004 C083 C120 C404 C169 C230 C178 C086 C451 C109 C092 C095 C030 C334 C038

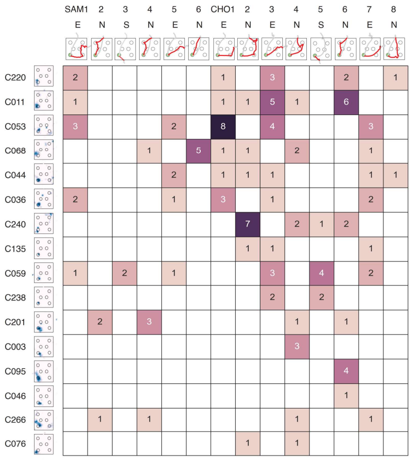
SAM1 2 3 4 5 6 CHO1 2 3 4 5 6 7 8
E N S N E N E N E N S N E N
C220 C011 C053 C068 C044 C036 C240 C135 C059 C238 C201 C003 C095 C046 C266 C076

**Figure S31 (previous page):** Further examples of startbox activity from multiple animals before the beginning of the trial during the allocentric task. For each panel, neurons representing the destination are presented as rows with cell ID and rate map. Session trials are presented as columns with trail ID, Start box and position of the animal during the execution of the trial. Shades of purple indicate the trials where the various cells are active in the startbox; numbers indicate the number of activity events. In the panels on the right, we also present the examples of neurons representing other, non-rewarded, sandwells, using shades of blue to indicate the trials where they were active for clarity.

| | SAM1 E | 2 E | 3 E | 4 E | 5 E | 6 E | CHO1 E | 2 E | 3 E | 4 E | 5 E | 6 E | 7 E | 8 E |
|---|---|---|---|---|---|---|---|---|---|---|---|---|---|---|
| C082 | | | 1 | | | | | | | | | | | |
| C117 | 4 | 2 | 6 | 1 | 5 | 8 | 7 | | 11 | 1 | 5 | | 3 | |
| C107 | 1 | | 1 | | 1 | 2 | 3 | 1 | 2 | 1 | 3 | 1 | 1 | |
| C162 | | 2 | 1 | 1 | 2 | 3 | 1 | 1 | 1 | 3 | 2 | 11 | 2 | |
| C087 | | | 1 | | 3 | | 1 | | 2 | | | | | |
| C125 | | | 12 | | 3 | 22 | 1 | | | | 2 | | | |
| C040 | | | | | 1 | | 4 | | 1 | | 1 | | | |
| C101 | | | | 2 | | | 2 | 1 | | 6 | 4 | 6 | | 5 |
| C137 | | | 1 | 1 | 3 | 8 | | | | | 7 | | | |
| C147 | | | | | | | | 1 | | 1 | 1 | | | |
| C113 | | | 1 | | | | | | 1 | | 1 | | | |
| C047 | 2 | | 2 | | | | | | 1 | | 1 | | | |
| C010 | | 2 | | | | | | | | | | | | |
| C151 | | 1 | | | | | | | | | | | | |
| C077 | | | | | | | | 1 | 1 | | | | | |
| C108 | | | | | 1 | 4 | | | 4 | | | | | |
| C116 | | | 1 | 1 | | | | | | | | 1 | | |
| C103 | | 6 | | | 6 | | | | | | | | | |
| C141 | | | 2 | | | | | | | | | | | |
| C136 | | | 1 | | | | | | | | | | | |
| C065 | 1 | | | 2 | | 2 | | | | | | | | |
| C076 | 14 | | | | | | | | | | | | | |
| C079 | 6 | | | | | | | | | | | | | |

| | SAM1 W | 2 W | 3 W | 4 W | 5 W | 6 W | CHO1 W | 2 W | 3 W | 4 W | 5 W | 6 W | 7 W | 8 W | 9 W | 10 W |
|---|---|---|---|---|---|---|---|---|---|---|---|---|---|---|---|---|
| C043 | 8 | | 13 | | | | 3 | | 1 | | 19 | | | | | |
| C429 | | | | | | | 6 | 5 | 7 | 6 | 6 | 6 | 14 | 5 | 6 | 6 |
| C435 | | | | | | | | | 4 | 1 | | | 2 | 1 | 1 | |
| C215 | | 1 | | | | | | | 3 | | | | 2 | | | |
| C077 | | | | 1 | | 1 | | 3 | | 5 | | 13 | | 10 | | 9 |
| C088 | | | | | 7 | | 1 | | | | 1 | | | | | |
| C036 | 5 | | | | | | | | 3 | 1 | | | 7 | | | |
| C087 | | | | | | | | 1 | | 2 | | | | | | |
| C226 | | | | | | | | 1 | | | | 1 | | | | |
| C053 | 4 | | | | | | | | | | | | | | | |

| | SAM1 W | 2 W | 3 W | 4 W | 5 W | 6 W | CHO1 W | 2 W | 3 W | 4 W | 5 W | 6 W | 7 W | 8 W |
|---|---|---|---|---|---|---|---|---|---|---|---|---|---|---|
| C057 | | | | | | | 12 | | | | 1 | | | 1 |
| C105 | 10 | 23 | 7 | 18 | 21 | 10 | 3 | 4 | 2 | 8 | 14 | 12 | 23 | 8 |
| C072 | 3 | | | | | | 19 | 19 | 18 | 18 | 16 | 16 | 18 | 18 |
| C086 | | 2 | 1 | 2 | | 2 | 5 | | | | 1 | | | |
| C031 | | 1 | 2 | | 2 | | 1 | | | | | | | |
| C002 | | | | | | | | 1 | | | | | | |

| | SAM1 E | 2 E | 3 E | 4 E | 5 E | 6 E | CHO1 E | 2 E | 3 E | 4 E | 5 E | 6 E | 7 E | 8 E |
|---|---|---|---|---|---|---|---|---|---|---|---|---|---|---|
| C234 | 1 | | 2 | | 3 | | 4 | | 1 | | | | | 2 |
| C059 | | | 3 | | 1 | | 2 | | 1 | | | 3 | | |
| C117 | 1 | | | | 2 | | 5 | | 3 | 3 | 6 | 4 | 2 | |
| C036 | | | | | | | 1 | | | | | | | |
| C310 | | | | | 1 | | 4 | | 1 | 1 | 4 | 2 | 1 | |
| C131 | 1 | | | | 2 | | 3 | | 1 | 1 | 4 | 2 | 1 | |
| C040 | 1 | | | | | | 2 | | | | 1 | | | |
| C243 | | 1 | | | | | | 2 | 1 | | | 1 | 1 | |
| C029 | | | | | | | | | 1 | | | | | 1 |
| C141 | 1 | | | | | | | | | | | | | |
| C076 | 12 | | | | | | | | | | | | | |
| C125 | | | | | | | 2 | | 2 | | | 2 | | |
| C107 | | 1 | | | 1 | | | | | | | | | |

| | SAM1 W | 2 W | 3 W | 4 W | 5 W | 6 W | CHO1 W | 2 W | 3 W | 4 W | 5 W | 6 W | 7 W | 8 W |
|---|---|---|---|---|---|---|---|---|---|---|---|---|---|---|
| C031 | 1 | | 1 | | 1 | | | | | | | | | |
| C277 | 6 | | | 1 | 3 | | | | | | | | | |
| C278 | | 9 | 1 | 7 | 1 | 13 | | | | | | 5 | | 5 |
| C106 | | | | | 1 | | 13 | 15 | 12 | 15 | 13 | 16 | 13 | 15 |
| C262 | 1 | | | 1 | | | | | | | | | | |
| C082 | | 1 | | | | | | | | | | | | |
| C086 | | 3 | | 1 | | | 1 | | | | | | | |
| C102 | | | | | | | 1 | | 4 | | | | | |
| C131 | | | | | | | | 1 | 4 | | | | 1 | |
| C436 | 1 | | | | | | | | | | | | | |
| C060 | 1 | | | | | | | | | | | | | |

| | SAM3 S | 4 S | 5 S | 6 S | CHO1 S | 2 S | 3 S | 4 S | 5 S | 6 S | 7 S | 8 S |
|---|---|---|---|---|---|---|---|---|---|---|---|---|
| C026 | 1 | | | | 3 | | | | 1 | | | |
| C147 | | | | | 4 | | 4 | | | | | |
| C137 | 1 | | 1 | 1 | 1 | | 1 | 1 | | 1 | 1 | 1 |
| C156 | | 2 | | | | | | | 7 | | | |
| C064 | | | | | | | | | 2 | | | |
| | | | | | | | | | 5 | 1 | | |
| C170 | 2 | | | | | | | | | | | |
| C148 | | | | | | | | | 2 | | | |
| C023 | | | | | | | | | 8 | | | |

**Figure S32 (previous page)**: Examples of startbox activity from multiple animals before the beginning of the trial during the egocentric task. For each panel, neurons representing the destination are presented as rows with cell ID and rate map. Session trials are presented as columns with trail ID, Start box and position of the animal during the execution of the trial. Shades of teal indicate the trials where the various cells are active in the startbox; numbers indicate the number of activity events.

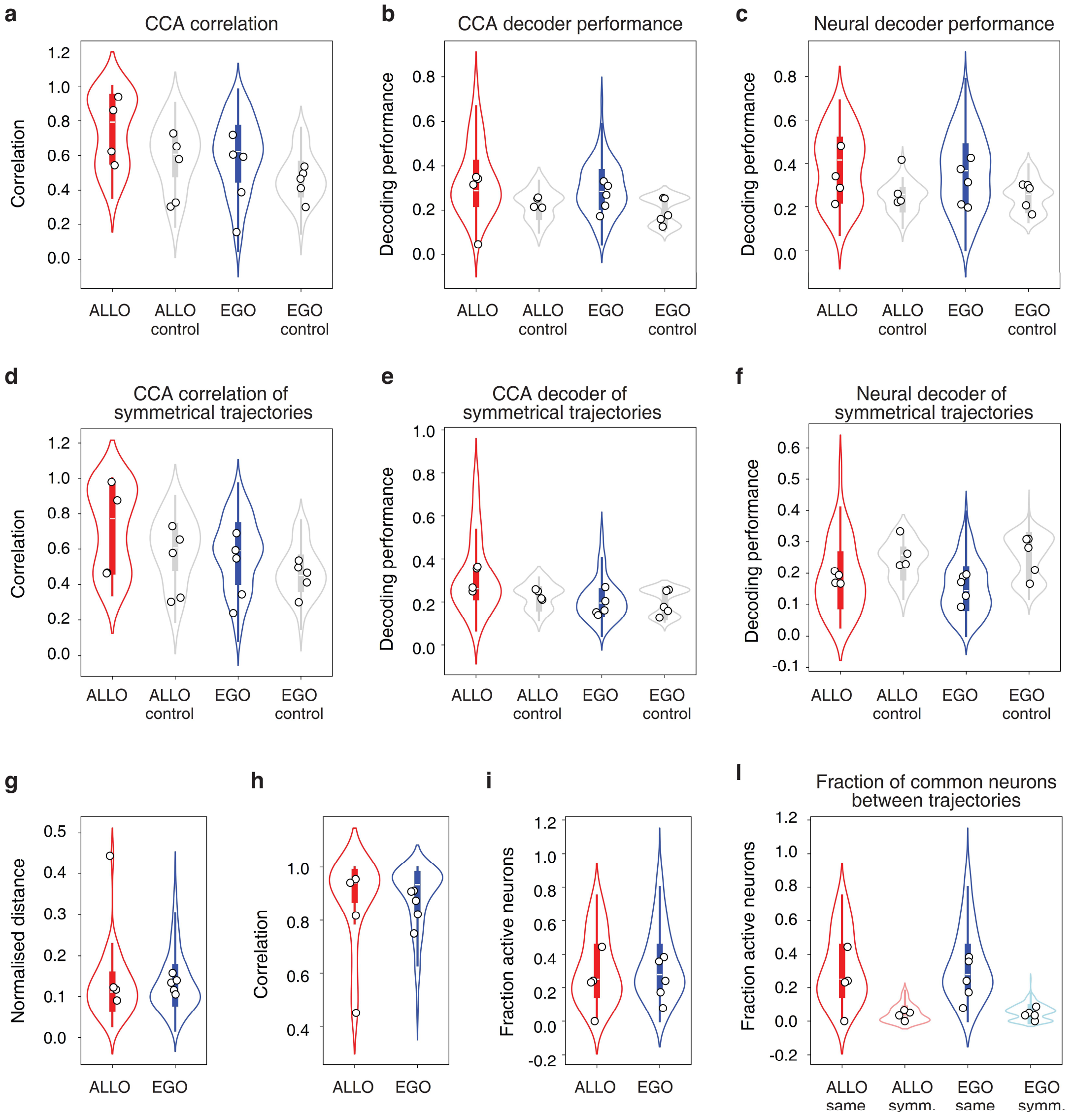

**Figure S33 CCA analysis to quantify the correlation between trials.** Individual points represent the average value for each animal. Controls were done by taking a random samples of neural activities from each animal. a) Average correlation along the first 5 components in CCA space for each pair of same-type trials (from the same startbox to the same reward-well). b-c) Decoders performance in CCA space and neural space when cross-validated on same-type trials. Chance levels given by the classes of the decoder. d) Same analysis as in (a) but on symmetrical trials (where the trials are overlapped if rotated 180°). e-f) Same analysis as in (b) and (c) but on symmetrical trials. g) Distance between the positions of the animal for each pair of trajectories, normalized by their length. h) Correlation between the positions of the animal for each pair of trajectories. i-l) Fraction of common neurons that are active between each pair of trials.

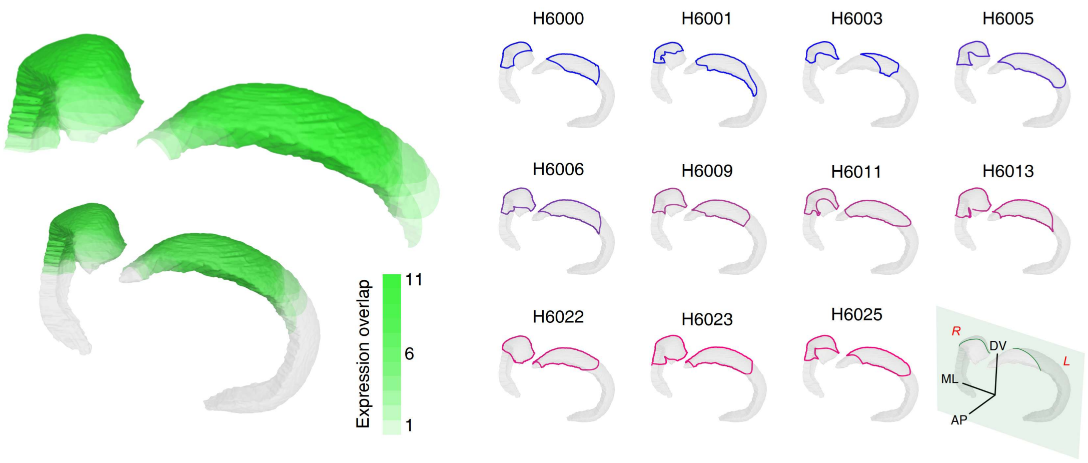

**Figure S34 Histological reconstruction of the extent of JAWS-GFP expression in the dorsal hippocampus.** Random animals infected with JAWS-GFP AAVs were selected and serial coronal sections of the hippocampus were cut every 200 µm and the presence of GFP fluorescence was used as indicator of transgene expression. Sections were matched against the rat atlas to confirm position along the AP axis (Paxinos & Watson, 2014).

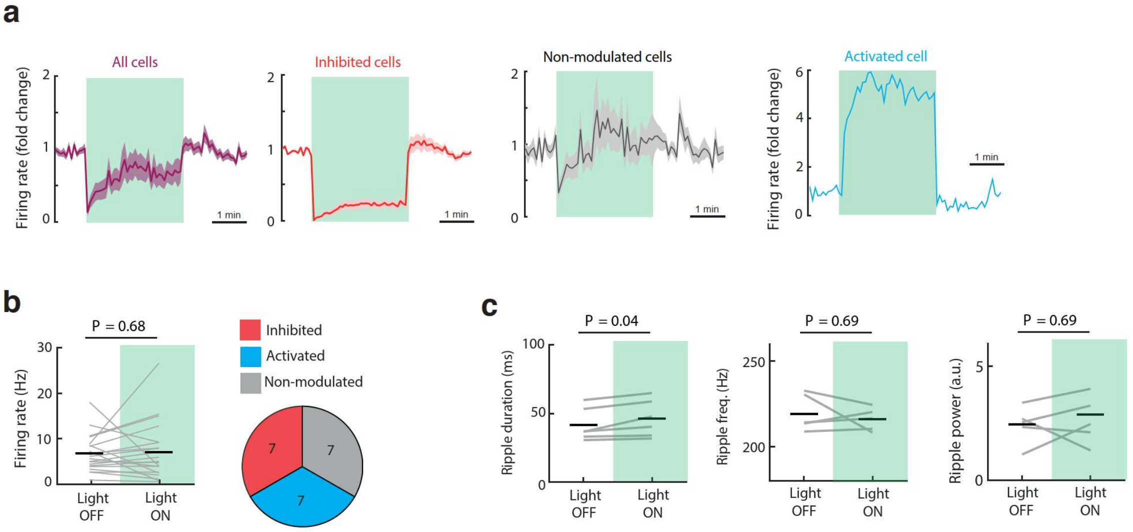

**Figure S35 Optogenetic inhibition of hippocampal activity with JAWS**. a) Firing rate changes of different categories of putative excitatory cells in CA1 before, during and after the light pulse: all recorded putative excitatory cells, significantly inhibited cells, non-modulated cells and one significantly activated cell. b) Left, mean firing rate of putative interneurons (n = 21 cells from 5 rats) did not change during light-ON periods (Wilcoxon Signed Rank Test, Z = 0.40, n = 21 cells, P > 0.05). Right, individually, putative interneurons showed heterogenous responses during light-ON with equal proportions of activated, inhibited and non-modulated cells. c) Properties of sharp wave-ripple (SWRs) events during Light ON and Light OFF periods. Duration of sharp wave-ripples (left) was modestly but significantly increased during light ON periods (Wilcoxon Signed Rank Test, Z = 2.02, n = 5 recordings from 5 animals, P < 0.05). On the other hand, both frequency (middle, Wilcoxon Signed Rank Test, Z = 0.40, n = 5 recordings from 5 animals, P > 0.05) and power (right, Wilcoxon Signed Rank Test, Z = 0.40, n = 5 recordings from 5 animals, P > 0.05) of SWRs did not change during light ON periods.

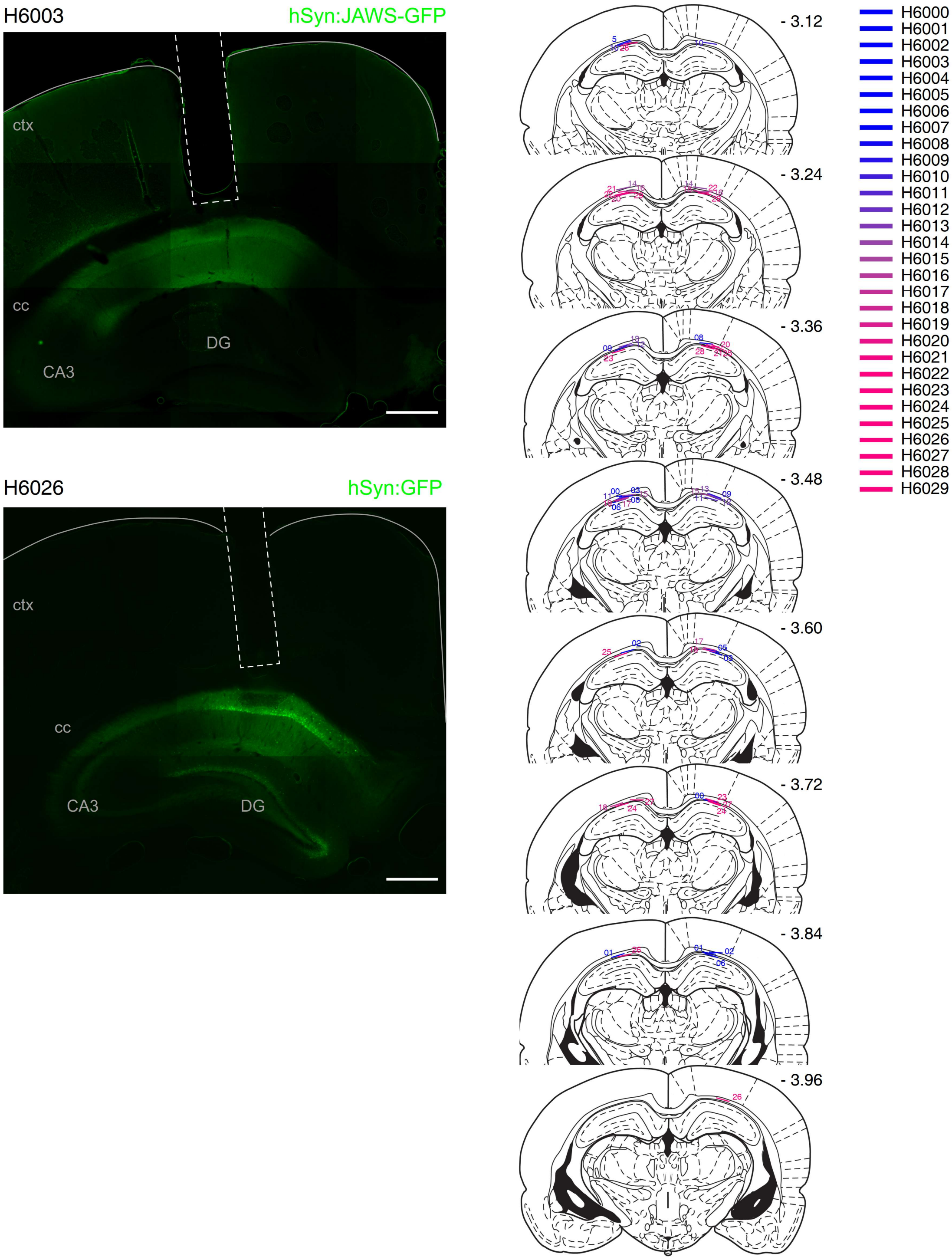

**Figure S36:** Coronal sections of animals expressing JAWS-GFP (top left) or GFP (bottom left). Dashed white lines mark the tract of the optic cannulas. Scale bars 500µm. cc: corpus callosum, ctx: cortex. Right panel, position of bilateral cannulae in animals used in the optogenetic experiments (Fig.5 and related Supplementary Figures) as confirmed by histological analysis. Coronal section derived from (Paxinos & Watson, 2014).

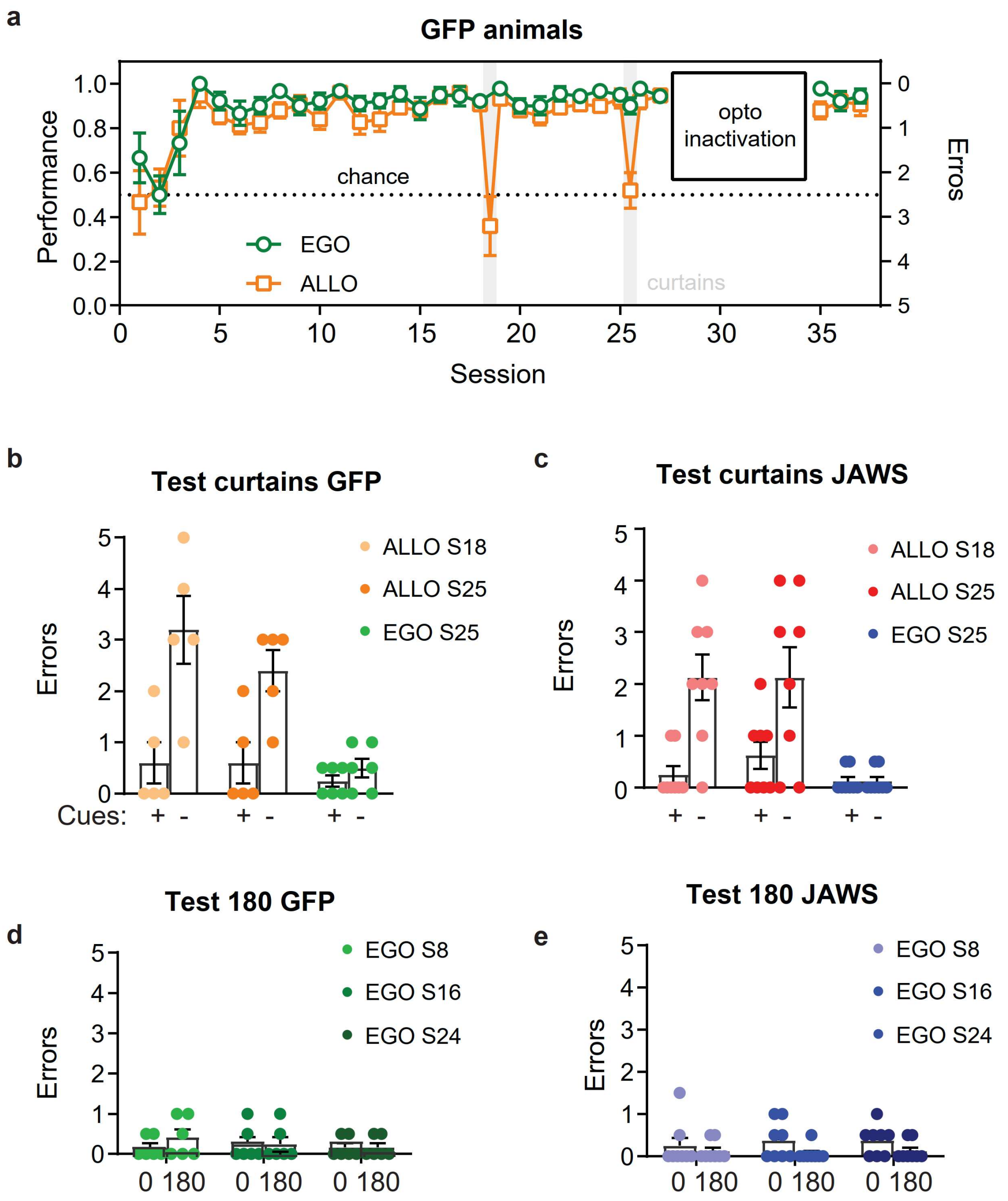

**Figure S37: Behavioral performance of animals used in the optogenetic experiment.** a) Performance index of GFP animals in the allocentric and egocentric groups. b,c) Number of errors for allocentric- and egocentric-trained rats when cues were masked with curtains. Values for the GFP and JAWS groups are presented in (b) and (c) respectively. d,e) Number of errors for egocentrically trained rats when performing in symmetrical trials (see Fig 1g and Methods). Values for the GFP and JAWS groups are presented in (d) and (e) respectively.

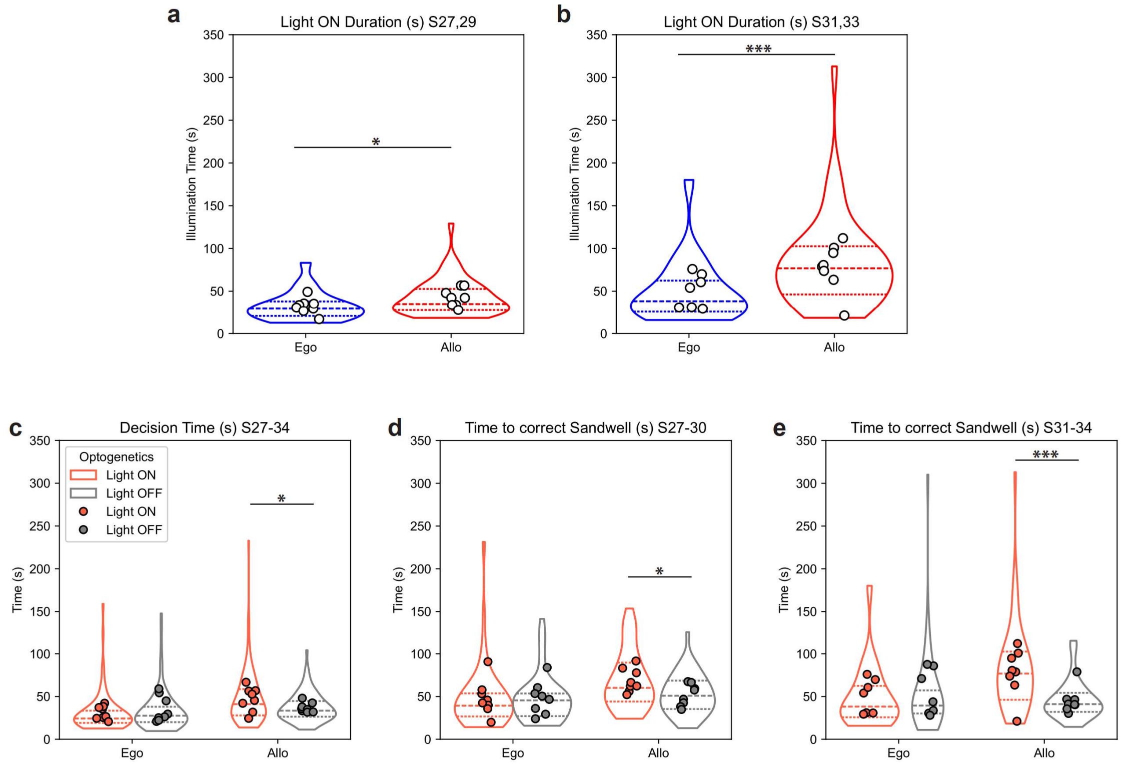

**Figure S38: Optogenetic illumination.** a,b) Duration of optogenetic silencing for animals in the Egocentric and Allocentric group in Light ON trials (see Methods) for Sessions 27 and 29 (a), and Sessions 31 and 33 (b). Note that the duration is longer for animals in the Allocentric group since the illumination was terminated when animals exited from the startbox (a; LMER Duration ~ Ego/Allo + (1|animal), P=0.036) or reached the correct Sandwell (b; LMER Duration ~ Ego/Allo + (1|animal), P=0.004). This is reflected by the longer Decision time (time spent in the startbox before starting the trial – as defined in Figures 3,4 as the moment the animals left the startbox – since the receiving of the cue pellet, and start of the illumination in Light-ON trials) in light ON compared to light OFF trials in the Allocentric, but not the Egocentric, group (c; LMER Time ~ Ego/Allo * light + (1|animal), light z=10.39 P=0.002, Ego/Allo * light z=-12.86 P=0.008; Bonferroni corrected post-hoc comparisons, Allo ON-OFF P=0.034, Ego ON-OFF P=0.99). Consistently, the time to reach the correct Sandwell was longer in both SB illumination trials (d; LMER Time ~ Ego/Allo * light + (1|animal), light z=17.01 P=0.005, Ego/Allo * light z=-15.86 P=0.067; Bonferroni corrected post-hoc comparisons, Allo ON-OFF P=0.04, Ego ON-OFF P=0.99) and Whole-trial illumination (e; LMER Time ~ Ego/Allo * light + (1|animal), light z=35.98 P<0.001, Ego/Allo * light z=-40.11 P=0.001; Bonferroni corrected post-hoc comparisons, Allo ON-OFF P=0.0005, Ego ON-OFF P=0.99 Violin plots are individual trials and filled points are animal averages.

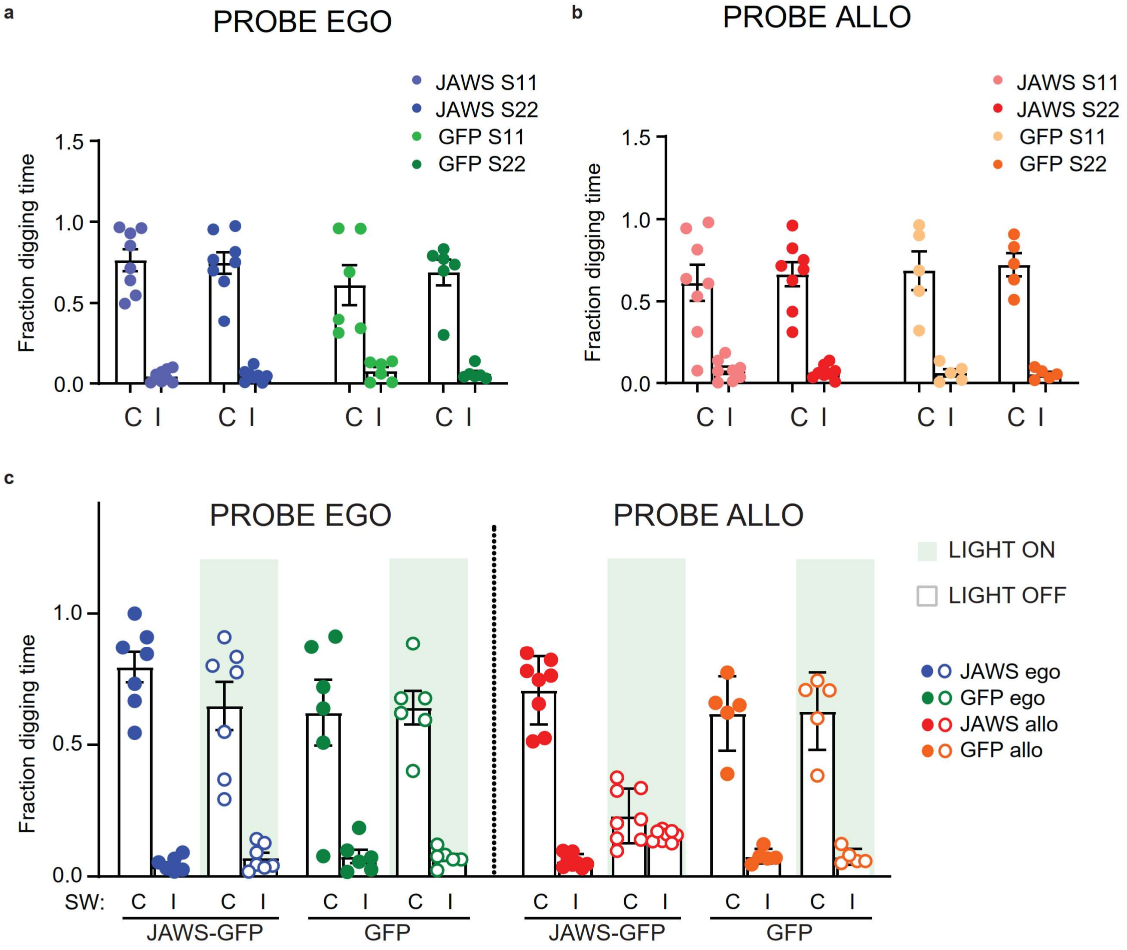

**Figure S39:** Fraction of digging time at correct (C) and incorrect (I) sandwells in probe sessions for egocentric- (a) and allocentric-trained animals (b). c) Probe sessions where the hippocampus was inactivated with red light (light ON) or the animals were handled equivalently but no light was delivered (light OFF).

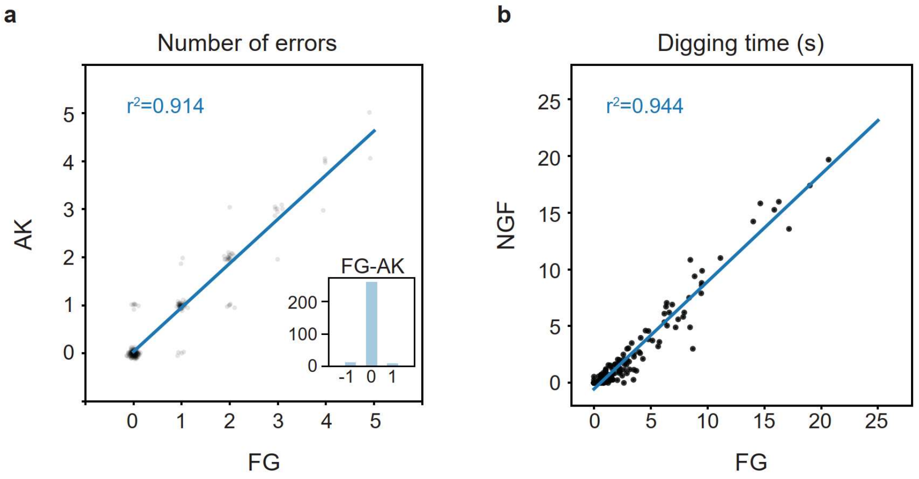

**Figure S40**: Comparison of scores between two experimenters. a) Number of errors calculated for a subset of randomly selected sessions for animals used in the optogenetic experiment. Inset, histogram showing the frequency of the difference between the experimenters' scores. b) Digging time in probe trials in calcium imaging experiments. The first user (FG) ran the experiment blind to virus and the second experimenter (AK, NGF) was blind to all conditions (including rewarded well).

### Supplementary Tables

Sandwells and startboxes combinations used in experiments.

### Supplementary Table 1

| | | | Animal ID | H2222-Group 1 | H2228-Group 1 | H2231-Group 1 | H2226-Group 1 | H2229-Group 1 | H2233-Group 1 | H2224-Group 2 | H2227-Group 2 | H2225-Group 2 | H2230-Group 2 | H2235-Group 3 | H2236-Group 3 | H2234-Group 3 | H2241-Group 3 |
|---|---|---|---|---|---|---|---|---|---|---|---|---|---|---|---|---|---|
| | | | Name | Frédéric | Johann Sebastian | Claude | Amadeus | Georges | Sergei | Johannes | Richard | Ludwig | Antonio | Kurt | Mick | Freddy | Kurt |
| | Habituation | | | X | X | X | X | X | X | X | X | X | X | X | X | X | X |
| | Habituation | | | X | X | X | X | X | X | X | X | X | X | X | X | X | X |
| | Habituation | | | X | X | X | X | X | X | X | X | X | X | X | X | X | X |
| | Habituation | | | X | X | X | X | X | X | X | X | X | X | X | X | X | X |
| | EGO / ALLO | Session | | H2222-Group 1 | H2228-Group 1 | H2231-Group 1 | H2226-Group 1 | H2229-Group 1 | H2233-Group 1 | H2224-Group 2 | H2227-Group 2 | H2225-Group 2 | H2230-Group 2 | H2235-Group 3 | H2236-Group 3 | H2234-Group 3 | H2241-Group 3 |
| Phase 1 | No cues in Ego | 1 | | E - 6 | N - 3 | W - 2 | E - 6 | N - 3 | W - 2 | S - 1 | W - 4 | S - 1 | W - 4 | E - 5 | S - 5 | E - 5 | S - 5 |
| | | 2 | | S - 1 | W - 4 | N - 3 | S - 1 | W - 4 | N - 3 | W - 6 | S - 2 | W - 6 | S - 2 | W - 6 | E - 3 | W - 5 | E - 3 |
| | | 3 | | E - 3 | E - 1 | S - 5 | E - 3 | E - 1 | S - 5 | N - 4 | W - 6 | N - 4 | W - 6 | W - 2 | N - 5 | W - 2 | E - 5 |
| | | 4 | | NNN/NNN - 4 | SSS/SSS - 6 | EEE/EEE - 1 | SWS/WSW - 4 | SWE/SSW - 5 | ESS/WSS - 1 | WWW/WWW - 5 | EEE/EEE - 3 | EWE/SWS - 5 | SWS/WEE - 3 | SSS/SSS - 1 | WWW/WWW - 3 | SWS/ESS - 1 | WSS/ESE - 3 |
| | | 5 | | NNN/NNN - 5 | SSS/SSS - 2 | EEE/EEE - 6 | SSW/ESS - 5 | WEE/WES - 2 | EWE/SSE - 6 | EEE/EEE - 2 | SSS/SSS - 4 | SWS/EWW - 2 | WEW/EES - 4 | NNN/NNN - 4 | EEE/EEE - 1 | ESS/WES - 4 | WSW/EES - 1 |
| | | 6 | | WWW/WWW - 2 | WWW/WWW - 5 | NNN/NNN - 1 | WES/SSE - 2 | WEE/SWS - 6 | SSE/WSE - 2 | SSS/SSS - 3 | EEE/EEE - 6 | WSE/ESW - 3 | SEW/SWS - 6 | SSS/SSS - 3 | WWW/WWW - 6 | ESS/WWS - 3 | SES/WSE - 5 |
| | | 7 | | SSS/SSS - 3 | NNN/NNN - 1 | WWW/WWW - 2 | SSE/WEW - 3 | SSW/ESE - 1 | WEW/SES - 1 | EEE/EEE - 6 | NNN/NNN - 4 | SWS/EWS - 6 | ESE/WSE - 4 | EEE/EEE - 5 | NNN/NNN - 2 | WWS/SSE - 6 | WWS/SSE - 4 |
| | | 8 | TEST 180 | WWW/WWEE - 5(2) | SSS/SSNN - 3(4) | EEE/EEWW - 6(1) | WEW/SWSE - 5 | EEW/SESW - 3 | SSE/WSWE - 6 | SSS/SSNN - 2(5) | EEE/EEWW - 4(3) | EWE/SWSE - 2 | WEW/SESW - 4 | WWW/WWEE - 2(5) | SSS/SSNN - 5(2) | SWS/EWES - 5 | ESE/WSWE - 3 |
| | | 9 | | EEE/EEE - 1 | EEE/EEE - 5 | SSS/SSS - 3 | EES/EWW - 1 | SEE/SWW - 5 | WWE/WSS - 3 | NNN/NNN - 4 | WWW/WWW - 2 | SWE/ESW - 4 | SES/WES - 2 | NNN/NNN - 6 | WWW/WWW - 6 | SWS/ESW - 6 | SSE/WWS - 6 |
| | | 10 | | SSS/SSS - 6 | EEE/EEE - 2 | NNN/NNN - 4 | SEW/WES - 6 | WEE/WEE - 2 | SEE/SWW - 4 | EEE/EEE - 5 | SSS/SSS - 1 | WES/WES - 5 | SWE/SWS - 1 | SSS/SSS - 1 | EEE/EEE - 4 | EES/WES - 1 | WES/SWE - 2 |
| | | 11 | PROBE | SSS/**S**SSS - 4 | WWW/**W**WWW - 6 | EEE/**E**EEE - 5 | WEW/**S**SWE - 4 | SSE/**W**WSS - 6 | SSW/**E**EES - 5 | NNN/**N**NNN - 3 | WWW/**W**WWW - 1 | SWS/**S**SSW - 3 | SEW/**W**WSE - 1 | NNN/NNNN - 2 | EEE/EEE - 4 | WSS/EESW - 2 | SWS/EEES - 3 |
| | | 12 | | NNN/NNN - 3 | WWW/WWW - 1 | EEE/EEE - 2 | EWE/SWS - 3 | WEW/SES - 4 | ESE/WSW - 2 | SSS/SSS - 2 | NNN/NNN - 5 | WSW/ESE - 2 | ESS/SWE - 5 | EEE/EEE - 5 | WWW/WWWW - 6* | SWS/SWE - 5 | SSW/SWE - 6 |
| | | 13 | | EEE/EEE - 2 | NNN/NNN - 4 | SSS/SSS - 3* | WEW/SWW - 2 | ESE/SWS - 4 | WSW/ESE - 3 | SSS/SSS - 1 | WWW/WWW - 6 | ESW/WSE - 1 | WSW/ESW - 6 | WWW/WWW - 4 | SSS/SSS - 1 | WSE/ESE - 4 | EES/EWW - 1 |
| | | 14 | | WWW/WWW - 5 | SSS/SSS - 3 | WWW/WWW - 1 | SWE/WWS - 2 | WWE/SWS - 3 | ESE/WWS - 1 | WWW/WWW - 6 | NNN/NNN - 3 | WSE/WSW - 6 | WSW/ESE - 4 | EEE/EEE - 3 | EEE/EEE - 5 | EES/SWW - 3 | SWS/WSW - 5 |
| | | 15 | | SSS/SSS - 1 | EEE/EEE - 2 | NNN/NNN - 6 | SWS/ESE - 1 | ESW/WEE - 2 | ESS/SEW - 6 | NNN/NNN - 5 | WWW/WWW - 4 | ESE/WSW - 5 | ESE/WSW - 3 | NNN/NNN - 2 | EEE/EEE - 5 | ESE/EWS - 2 | SWE/EWS - 4 |
| | | 16 | TEST 180 | SSS/SSNN - 3(4) | WWW/WWEE - 5(2) | SSS/SSNN - 1(6) | SES/WSWS - 3 | SWS/ESEW - 5 | ESE/WEWS - 1 | NNN/NNSS - 5(2) | EEE/EEWW - 3(4) | WSW/ESEW - 5 | SES/WSWE - 4 | SSS/SSNN - 6(1) | WWW/WWEE - 2(5) | ESE/WSWE - 6 | WSW/ESEW - 2 |
| | | 17 | | NNN/NNN - 6 | NNN/NNN - 1 | EEE/EEE - 5 | ESE/SWS - 6 | SES/WEW - 1 | WEW/SWE - 5 | WWW/WWW - 1 | SSS/SSS - 2 | SES/WSW - 1 | SWS/EES - 2 | EEE/EEE - 5 | NNN/NNN - 1 | SWW/WSE - 5 | EWS/EWS - 3 |
| | Cues in Allo and Ego (with the exception of curtain trials) | 18 | CUES (E) /NO CUES (A) | SSS/SSS - 4 | SSS/SSS - 6 | WWW/WWW - 6 | WEW/SWEs - 4 | SES/SEWe - 6 | SWS/EWSw - 6 | EEE/EEE - 3 | WWW/WWW - 5 | SSW/ESWw - 2 | EES/WSEe - 5 | EEE/EEE - 5 | SSS/SSS - 6 | SSW/ESSe - 3 | SEW/ESSe - 1 |
| | | 19 | | NNN/NNN - 3 | NNN/NNN - 1 | EEE/EEE - 2 | EES/WEW - 3 | WES/EWS - 1 | WWS/SEE - 2 | SSS/SSS - 4 | SSS/SSS - 2 | SEW/WWS - 4 | SWE/SSW - 2 | WWW/WWW - 1 | EEE/EEE - 4 | WES/WEE - 1 | SWE/WES - 4 |
| | | 20 | | EEE/EEE - 1 | WWW/WWW - 2 | NNN/NNN - 3 | SWE/ESW - 2 | SES/WSW - 3 | WES/SEW - 1 | SSS/SSS - 2 | NNN/NNN - 1 | WSE/SEW - 2 | WWE/SSW - 1 | SSS/SSS - 2 | WWW/WWW - 3 | EES/WSE - 3 | SSW/EES - 3 |
| | | 21 | | WWW/WWW - 6 | EEE/EEE - 5 | SSS/SSS - 4 | WES/WES - 6 | EES/WSE - 5 | ESE/WSE - 4 | WWW/WWW - 6 | EEE/EEE - 4 | ESE/SWE - 6 | SSW/ESE - 4 | NNN/NNN - 3 | WWW/WWW - 6 | WWS/WES - 2 | WWE/WSW - 6 |
| | | 22 | PROBE | SSS/**s**SSS - 3 | SSS/**s**SSS - 6 | NNN/**n**NNN - 5 | SSE/WWSE - 3 | SWS/**S**SWE - 6 | EWE/**E**ESW - 5 | EEE/**E**EEE - 1 | WWW/**W**WWW - 2 | ESW/**W**WSE - 2 | WSE/**E**ESW - 1 | SSS/SSSS - 4 | EEE/EEEE - 1 | EWS/SSWE - 3 | WSE/EESW - 1 |
| | | 23 | | NNN/NNN - 4 | WWW/WWW - 1 | EEE/EEE - 2 | ESE/WSS - 4 | WEW/SES - 1 | WWS/ESE - 2 | SSS/SSS - 5 | NNN/NNN- 6 | SSW/WSE - 5 | WES/SSE - 6 | EEE/EEE - 6 | NNN/NNN - 4 | SWE/SWE - 5 | SWW/SWE - 3 |
| | | 24 | TEST 180 | SSS/SSNN - 3(4) | EEE/EEWW - 6(1) | WWW/WWEE - 5(2) | SES/WSWE - 3 | SWS/EWES - 6 | WSW/EWES - 5 | NNN/NNSS - 3(4)) | SSS/SSNN - 1(6) | EWW/SWSE - 3 | WEW/SWSE - 1 | WWW/WWEE - 2(5) | SSS/SSNN - 3(4) | ESE/WSWE - 1 | WSS/ESEW - 5 |

| | | | | | | | | | | | | | | | | | |
|---|---|---|---|---|---|---|---|---|---|---|---|---|---|---|---|---|---|
| | | 25 | CURTAINS | WWW/WWww - 2 | SSS/SSss - 5 | SSS/SSss - 1 | EWE/SWEsw - 2 | WEW/SEWse - 5 | SWW/ESWes - 1 | EEE/EEee - 6 | SSS/SSss - 3 | SWS/ESWew - 6 | ESE/SSWes - 3 | EEE/EEee - 5 | WWW/WWww - 1 | SWW/SWEw - 6 | EES/SESe - 2 |
| | | 26 | | EEE/EEE - 5 | NNN/NNN - 2 | NNN/NNN - 3 | EWS/SWE - 5 | SSW/WSE - 2 | SWE/WSE - 3 | WWW/WWW - 1 | EEE/EEE - 1 | WWS/SWE - 1 | SEE/WSE - 5 | NNN/NNN - 1 | NNN/NNN - 5 | SES/WSE - 2 | SES/SWS - 6 |
| | | 27 | | NNN/NNN - 3 | EEE/EEE - 4 | EEE/EEE - 2 | WSW/WES - 3 | EWS/SWE - 4 | ESE/SWE - 6 | NNN/NNN - 3 | NNN/NNN - 5 | SEE/ESW - 3 | SES/WSW - 2 | SSS/SSS - 3 | EEE/EEE - 3 | WSS/ESW - 4 | WES/SEW - 4 |
| | | 28 | RECORD | EEE/EEEE - 5 | EEE/EEE - 2 | WWW/WWWW - 6 | WEW/WSWE - 6 | SWS/WEE - 2 | EEW/SWS - 5 | SSS/SSSS - 2 | WWW/WWW - 1 | SEE/EEWS - 5 | WSW/WEWW - 1 | EEE/EEEE - 5 | SSS/SSS - 2 | WSW/WEWW - 1 | WSW/EWWS - 5 |
| | | 29 | RECORD | WWW/WWWW - 2 | SSS/SSS - 5 | EEE/EEEE - 1 | EWE/EESS - 1 | WES/EWS - 5 | SWE/WWS - 4 | WWW/WWWW - 1 | SSS/SSS - 4 | WEW/EWWS - 2 | SEE/SWEE - 6 | WWW/WWWW - 2 | WWW/WWW - 1 | SEE/SWEE - 6 | WEW/WSWE - 6 |
| | | 30 | RECORD | SSS/SSSS - 6 | NNN/NNN - 6 | SSS/SSSS - 5 | SWE/ESSW - 5 | SES/SWE - 6 | SWW/WES - 2 | EEE/EEEE - 6 | EEE/EEE - 1 | WSS/SESE - 1 | SEW/WSEW - 2 | SSS/SSSS - 6 | EEE/EEE - 6 | SEW/WSEW - 2 | EWE/EESS - 1 |
| | | 31 | RECORD | EEE/EEEE - 5 | WWW/WWW - 1 | WWW/WWWW - 6 | WSW/WWES - 6 | WWS/SEE - 1 | WSW/WES - 3 | WWW/WWWW - 1 | SSS/SSS - 5 | ESE/EESE - 5 | WEE/WEWE - 6 | EEE/EEEE - 5 | WWW/WWW - 3 | WEE/SEWE - 6 | SEW/ESEW - 5 |
| | | 32 | RECORD | SSS/SSS(nnn) - 6(1) | NNN/NNNsss - 3(4) | SSS/SSSS(nnn) - 3(4) | EES/WEWES S - 1 | SWS/EWES - 6 | SSW/EWES - 1 | SSS/SSSSnn n - 4(3) | NNN/NNSS - 2(5) | SWS/SESEW W - 1 | EWW/SWSW EE - 1 | SSS/SSSS(nn nn) - 6(1) | SSS/SSnn - 2(5) | EWW/SWSW EE - 1 | EES/WEWES S - 1 |
| | | 33 | RECORD | SSS/SSSS - 3 | WWW/WWW - 2 | EEE/EEEE - 1 | SEE/SEWE - 5 | EWE/SEW - 3 | SWS/SWE - 4 | EEE/EEEE - 6 | NNN/NNN - 6 | WES/ESES - 5 | ESE/EWEW - 2 | EEE/EEEE - 1 | SSS/SSS - 4 | WSE/WEWW - 2 | WSW/ESES - 2 |
| | | 34 | RECORD | WWW/WWW(www) - 2 | SSS/SSss - 4 | EEE/EEEE(eeee) - 5 | ESS/SESEs - 2 | SWS/EWSs - 4 | EWS/SWEe - 6 | EEE/EEEEeee - 2 | SSS/SSss - 3 | SWW/WSWE w - 2 | SWE/WWEWs - 1 | WWW/WWW W(wwww) - 2 | EEE/EEee - 6 | SWE/WWSWe - 1 | SWW/WWESe - 6 |
| | | 35 | | NNN/NNN - 1 | SSS/SSS - 3 | NNN/NNN - 6 | WSE/WWS - 3 | SWW/WEW - 1 | EWW/SWW - 2 | SSS/SSS - 3 | NNN/NNN - 5 | ESW/SEW - 4 | SWS/EWS - 5 | NNN/NNN - 3 | SSS/SSS - 4 | SSW/SEW - 3 | WSW/WSE - 4 |
| | | 36 | PROBE | WWW/WWW W - 6 | EEE/EEEE - 2 | SSS/SSSS - 3 | SWS/SESW - 6 | EWE/WWSE - 2 | WSW/SSWE - 3 | NNN/NNNN - 5 | EEE/EEEE - 1 | EWS/EESS - 5 | WSE/SESW - 4 | SSS/SSSS - 1 | WWW/WWW W - 3 | SWS/SSEW - 1 | EWE/EESS - 3 |
| | | **Session** | | **H2222-Group 1** | **H2228-Group 1** | **H2231-Group 1** | **H2226-Group 1** | **H2229-Group 1** | **H2233-Group 1** | **H2224-Group 2** | **H2227-Group 2** | **H2225-Group 2** | **H2230-Group 2** | **H2235-Group 3** | **H2236-Group 3** | **H2234-Group 3** | **H2241-Group 3** |
| **Phase 2** | No cues in Ego | 1 | | ESE/WSW - 6 | SWS/ESE - 3 | ESE/WSW - 2 | EEE/EEE - 6 | NNN/NNN - 3 | WWW/WWW - 2 | EWS/SSE - 1 | ESE/WEW - 4 | SSS/SSS - 1 | WWW/WWW - 4 | SWS/EWE - 5 | ESS/WEW - 2 | EEE/EEE - 5 | SSS/SSS - 4 |
| | | 2 | | SWS/ESE - 1 | WSW/ESE - 4 | SWE/ESW - 3 | SSS/SSS - 1 | WWW/WWW - 4 | NNN/NNN - 3 | SWE/WEW - 6 | WEW/ESS - 2 | WWW/WWW - 6 | SSS/SSS - 2 | EEE/WSW - 2 | SSW/ESW - 1 | WWW/WWW - 4 | EEE/EEE - 1 |
| | | 3 | | ESS/WES - 3 | EWE/SES - 1 | SWS/SEW - 5 | EEE/EEE - 3 | EEE/EEE - 1 | SSS/SSS - 5 | EWE/WSS - 4 | ESS/SEW - 6 | NNN/NNN - 4 | WWW/WWW - 6 | SSW/ESE - 1 | WSW/ESW - 3 | SSS/SSS - 2 | NNN/NNN - 6 |
| | | 4 | | SWS/WSW - 4 | SWE/SSW - 5 | ESS/WSS - 1 | NNN/NNN - 4 | SSS/SSS - 6 | EEE/EEE - 1 | EWE/SWS - 5 | SWS/WEE - 3 | WWW/WWW - 5 | EEE/EEE - 3 | WEE/SWW - 6 | EWW/SEE - 5 | NNN/NNN - 6 | EEE/EEE - 5 |
| | | 5 | | SSW/ESS - 5 | WEE/WES - 2 | EWE/SSE - 6 | NNN/NNN - 5 | SSS/SSS - 2 | EEE/EEE - 6 | SWS/ESW - 3 | EWE/SWE - 2 | EEE/EEE - 3 | NNN/NNN - 2 | SWW/EWS - 2 | ESE/WSE - 3 | EEE/EEE - 3 | SSS/SSS - 2 |
| | | 6 | | WES/SSE - 2 | WEE/SWS - 6 | SSE/WSE - 2 | WWW/WWW - 2 | EEE/EEE - 5 | NNN/NNN - 1 | EWE/SWS - 2 | SES/WSW - 5 | EEE/EEE - 2 | SSS/SSS - 2 | WWS/ESS - 5 | SSW/WSS - 1 | SSS/SSS - 5 | NNN/NNN - 1 |
| | | 7 | | SSE/WEW - 3 | SSW/ESE - 1 | WEW/SES - 1 | SSS/SSS - 3 | NNN/NNN - 1 | WWW/WWW - 2 | WWS/ESW - 4 | WSE/WEE - 1 | WWW/WWW - 5 | NNN/NNN - 4 | EES/WSE - 4 | WEE/SEW - 6 | WWW/WWW - 4 | EEE/EEE - 6 |
| | | 8 | TEST 180 | WEW/SWSE - 5 | EEW/SESW - 3 | SSE/WSWE - 6 | WWW/WWEE - 5(2) | SSS/SSNN - 3(4) | EEE/EEWW - 6(1) | SWS/EWES - 2 | ESS/WSWE - 3 | EEE/EEWW - 2(5) | SSS/SSNN - 3(4) | SWS/ESEW - 6 | SES/WSWE - 2 | SSS/SSNN - 6(1) | WWW/WWEE - 2(5) |
| | | 9 | | EES/EWW - 1 | SEE/SWW - 5 | WWE/WSS - 3 | NNN/NNN - 1 | WWW/WWW - 6 | SSS/SSS - 3 | EWE/SWS - 1 | WES/SWS - 4 | SSS/SSS - 4 | EEE/EEE - 1 | EWW/ESW - 1 | EWW/ESW - 4 | WWW/WWW - 1 | SSS/SSS - 4 |
| | | 10 | | SEE/WEW - 6 | ESE/WES - 4 | SWS/WES - 4 | EEE/EEE - 5 | EEE/EEE - 2 | SSS/SSS - 4 | SWE/SWE - 6 | SEW/SWE - 5 | NNN/NNN - 6 | WWW/WWW - 5 | SWW/SWE - 2 | SWS/WSE - 5 | NNN/NNN - 2 | WWW/WWW - 6 |
| | | 11 | PROBE | WEW/WSWE - 4 | SSE/WWSE - 6 | SSW/SEES - 5 | SSS/SSSS - 4 | WWW/WWW W - 6 | EEE/EEEE - 5 | SWS/SSSW - 3 | SEW/WWSE - 1 | EEE/EEEE - 1 | WWW/WWW W - 2 | EEW/SSWW - 2 | WSS/WEWS - 4 | EEE/EEEE - 2 | EEE/EEEE - 3 |
| | | 12 | | ESW/EWS - 2 | EWS/SEW - 6 | SSW/ESE - 2 | EEE/EEE - 2 | SSS/SSS - 1 | NNN/NNN - 4 | EWS/SWE - 2 | SSE/WES - 3 | WWW/WWW - 2 | EEE/EEE - 3 | SSW/ESE - 3 | SSW/ESW - 1 | SSS/SSS - 3 | SSS/SSS - 1 |
| | | 13 | | SWE/SEW - 2 | SEW/ESE - 1 | ESW/SWS - 6 | WWW/WWW - 5 | NNN/NNN - 6 | SSS/SSS - 3 | SWE/WSE - 5 | EWS/SWE - 2 | WWW/WWW - 6 | NNN/NNN - 6 | WEE/SSW - 5 | EWS/SEW - 3 | WWW/WWW - 5 | EEE/EEE - 3 |
| | | 14 | | WES/SEW - 5 | ESE/WSW - 3 | ESW/SWW - 6 | SSS/SSS - 1 | EEE/EEE - 3 | WWW/WWW - 2 | EWS/WES - 4 | SSW/SEW - 1 | SSS/SSS - 3 | NNN/NNN - 1 | SWW/ESW - 6 | EES/EWS - 6 | NNN/NNN - 6 | SSS/SSS - 6 |
| | | 15 | | EWS/EWS - 6 | SWE/SSW - 5 | WSE/SEW - 5 | WWW/WWW - 6 | SSS/SSS - 4 | NNN/NNN - 1 | WES/ESW - 1 | EWS/SEW - 6 | EEE/EEE - 1 | SSS/SSS - 5 | EES/WWS - 4 | SWE/WEW - 1 | EEE/EEE - 3 | WWW/WWW - 1 |
| | | 16 | TEST 180 | SES/WSWS - 3 | SWS/ESEW - 5 | ESE/WEWS - 1 | SSS/SSNN - 3(4) | EEE/EEWW - 6(1) | WWW/WWEE - 5(2) | ESE/WEWS - 6 | WSW/ESEW - 2 | NNN/NNSS - 6(1) | EEE/EEWW - 3(4) | ESE/WSWE - 1 | WSS/ESEW - 5 | WWW/WWEE - 2(5) | SSS/SSNN - 3(4) |
| | | 17 | | EWS/SEW - 1 | ESW/SSW - 4 | EWS/SWE - 2 | NNN/NNN - 6 | NNN/NNN - 1 | EEE/EEE - 5 | SEW/EWS - 3 | SSW/SEW - 4 | SSS/SSS - 3 | EEE/EEE - 6 | SWW/ESE - 1 | WWS/ESS - 1 | SSS/SSS - 1 | EEE/EEE - 1 |
| | Cues in Allo and Ego (with the exception of curtain trials) | 18 | CUES (E) /NO CUES (A) | WEW/SWEs - 4 | SES/SEWe - 6 | SWS/EWSw - 6 | SSS/SSS - 4 | SSS/SSS - 6 | WWW/WWW - 6 | ESW/SWEw - 2 | SWE/WESw - 6 | NNN/NNN - 4 | EEE/EEE - 2 | SSW/WSWw - 6 | ESE/SEWw - 2 | NNN/NNN - 6 | EEE/EEE - 1 |
| | | 19 | | EWS/EWS - 1 | WES/EWS - 3 | SWE/ESE - 4 | EEE/EEE - 2 | WWW/WWW - 4 | NNN/NNN - 3 | SWE/WSS - 5 | WWS/WEE - 2 | WWW/WWW - 5 | WWW/WWW - 5 | EWS/SWE - 3 | SWW/WEW - 1 | WWW/WWW - 4 | SSS/SSS - 2 |
| | | 20 | | SEW/ESW - 6 | ESW/ESE - 2 | WES/EWS - 3 | EEE/EEE - 1 | SSS/SSS - 5 | SSS/SSS - 4 | ESW/WES - 4 | WES/EEW - 3 | NNN/NNN - 2 | NNN/NNN - 3 | WWS/WSE - 2 | SWW/EWS - 5 | EEE/EEE - 2 | NNN/NNN - 5 |
| | | 21 | | EES/WSW - 5 | WSS/ESW - 1 | SSW/WES - 6 | WWW/WWW - 5 | NNN/NNN - 3 | EEE/EEE - 1 | SEW/ESW - 5 | ESW/WES - 6 | EEE/EEE - 1 | SSS/SSS - 6 | WWE/SWE - 5 | WWS/SWE - 6 | WWW/WWW - 5 | WWW/WWW - 6 |

| | | | | | | | | | | | | | | | |
|---|---|---|---|---|---|---|---|---|---|---|---|---|---|---|---|
| 22 | PROBE | SSE/WWSE - 3 | SWS/SSWE - 6 | EWE/EESW - 5 | SSS/sSSS - 3 | SSS/sSSS - 6 | NNN/nNNN - 5 | ESW/WWSE - 2 | WSE/EESW - 1 | EEE/EEEE - 1 | WWW/WWWW - 2 | EWW/SSES - 4 | SWS/EESS - 1 | SSS/SSSS - 3 | EEE/EEEE - 1 |
| 23 | | WSE/SWW - 1 | ESE/WSE - 2 | SSE/WSW - 3 | NNN/NNN - 4 | WWW/WWW - 2 | EEE/EEE - 2 | SWE/WSW - 2 | ESE/WES - 6 | SSS/SSS - 2 | EEE/EEE - 5 | SWW/EES - 3 | ESS/WES - 3 | NNN/NNN - 1 | WWW/WWW - 4 |
| 24 | TEST 180 | SES/WSEW - 3 | SWS/EWES - 6 | WSW/EWES - 5 | SSS/SSNN - 3(4) | EEE/EEWW - 6(1) | WWW/WWEE - 5(2) | EWW/SWSE - 3 | WEW/SWSE - 1 | NNN/NNSS - 3(4) | SSS/SSNN - 1(6) | SWS/ESEW - 2 | EWE/SWSE - 1 | EEE/EEWW - 5(2) | SSS/SSNN - 6(1) |
| 25 | CURTAINS | EWE/WSEwe - 2 | WEW/SEWes - 5 | SWE/EWSes - 1 | WWW/WWww - 2 | SSS/SSss - 5 | SSS/SSss - 1 | SWS/ESW*ew* - 6 | ESE/WSS*ns* - 3 | EEE/EEee - 6 | SSS/SSss - 3 | ESW/EWSsw - 1 | SWW/EWSse - 4 | SSS/SSss - 2 | SSS/SSss - 3 |
| 26 | | SWE/SEW - 4 | SEW/ESE - 3 | SWE/SEW - 2 | NNN/NNN - 5 | NNN/NNN - 2 | EEE/EEE - 6 | SWE/SEW - 1 | SWE/SSW - 2 | SSS/SSS - 1 | NNN/NNN - 4 | WWS/WES - 5 | SWS/SEW - 1 | WWW/WWW - 6 | EEE/EEE - 2 |
| 27 | | WSE/SWE - 6 | WES/WSE - 2 | ESW/SEW - 4 | SSS/SSS - 2 | WWW/WWW - 1 | NNN/NNN - 4 | WES/WSE - 1 | WWS/EWS - 5 | WWW/WWW - 2 | WWW/WWW - 5 | SEW/SEW - 4 | SEE/SWE - 4 | SSS/SSS - 4 | NNN/NNN - 4 |
| 28 | RECORD | SEE/EEW - 5 | WSE/SWS - 1 | WEW/WSWE - 6 | WWW/WWWW - 6 | EEE/EEE - 6 | SSS/SSS - 3 | WSW/WEWW - 1 | EWS/SWE - 4 | EEE/EEEE - 5 | SSS/SSSS - 2 | SEE/WES - 5 | SEW/ESW - 5 | WWW/WWWW - 5 | SSS/SSSS - 5 |
| 29 | RECORD | WEW/WWS - 2 | EWE/SES - 5 | EWE/EESS - 1 | EEE/EEEE - 1 | SSS/SSS - 6 | WWW/WWW - 5 | SEE/SWEE - 6 | WES/SWW - 5 | WWW/WWWW - 2 | WWW/WWWW - 1 | SWW/EWS - 2 | ESS/WES - 6 | WWW/WWWW - 1 | WWW/WWWW - 6 |
| 30 | RECORD | W**SS/S***E***S***E* - 1 | SWE/WES - 4 | SW*E*/*E*S*E*W - 5 | SSS/SSSS - 5 | NNN/NNN - 5 | EEE/EEE - 2 | S*E***W/WS***E***W** - 2 | EWS/EWE - 1 | SSS/SSSS - 6 | EEE/EEEE - 6 | EWE/ESE - 1 | EWS/SEW - 3 | EEE/EEEE - 6 | EEE/EEEE - 1 |
| 31 | RECORD | ESE/ESEE - 5 | EWE/SWS - 6 | WSW/WWES - 6 | WWW/WWWW - 6 | EEE/EEE - 2 | WWW/WWW - 1 | WEE/WEWE - 6 | SES/WSW - 2 | EEE/EEEE - 5 | WWW/WWWW - 1 | ESS/SWS - 6 | EWS/SSW - 2 | WWW/WWWW - 5 | WWW/WWWW - 6 |
| 32 | RECORD | **SWS/S***E***S***E*WW - 1 | SWS/EWSE - 2 | EES/WEWESS - 1 | SSS/SSSS(nnnn) - 3(4) | WWW/WWEE - 5(2) | EEE/EEWW - 6(1) | EWW/SWSWEE - 1 | WEW/SWSE - 3 | SSS/SSSS(nnnn) - 6(1) | SSS/SSSSnnnn - 2(5) | WSW/EWES - 2 | WSS/ESEW - 5 | SSS/SSSSnnnn - 3 | SSS/SSSSnnnn - 5(2) |
| 33 | RECORD | WE**S**/E**S**E - 5 | ESW/SSE - 3 | S*EE*/S*E*W*E* - 5 | EEE/EEEE - 1 | SSS/SSS - 1 | NNN/NNN - 2 | *E***S***E*/*E***W***E***W** - 2 | WEW/SEW - 5 | EEE/EEEE - 1 | EEE/EEEE - 6 | SEE/WES - 5 | SSW/ESS - 1 | WWW/WWWW - 1 | EEE/EEEE - 5 |
| 34 | RECORD | SWW/WSWEw - 2 | EWS/SEWe - 6 | ESS/SESEs - 2 | EEE/EEEE(eeee) - 5 | NNN/NNnn - 4 | SSS/SSss - 5 | SWE/WWEWe - 1 | SEW/WSEw - 2 | WWW/WWWW(wwww) - 2 | EEE/EEEEeeee - 2 | SWS/SWEe - 6 | SWW/WSE - 1 | EEE/EEEEeeee - 6 | EEE/EEEEeeee - 1 |
| 35 | | WES/SWE - 3 | WSE/SWW - 5 | ESS/WES - 3 | WWW/WWW - 2 | SSS/SSS - 6 | EEE/EEE - 4 | WES/WES - 3 | SSW/SSE - 5 | SSS/SSS - 3 | NNN/NNN - 5 | SEW/EWS - 3 | WSW/SES - 2 | SSS/SSS - 2 | SSS/SSS - 3 |
| 36 | PROBE | SWE/SSWE - 6 | EWE/WWSE - 2 | WSW/SSWE - 3 | WWW/WWWW - 6 | EEE/EEEE - 2 | SSS/SSSS - 3 | EWS/ESEW - 5 | EWS/SESW - 4 | NNN/NNNN - 5 | EEE/EEEE - 1 | SWS/SSEW - 1 | EWE/EESS - 3 | WWW/WWWW - 3 | SSS/SSSS - 1 |
| | non cued | E - 4 | N - 6 | S - 2 | E - 4 | N - 6 | S - 2 | W - 4 | E - 5 | W - 4 | E - 6 | E - 5 | W - 1 | E - 1 | W - 5 |

### Supplementary Table 2

| | 2300hZ, DELAY 5S, 0.5 S DURATION | Animal ID | H6000-Group 1 | H6002-Group 1 | H6004-Group 1 | H6006-Group 1 | H6001-Group 1 | H6003-Group 1 | H6005-Group 1 | H26010-Group 1 | H6009-Group 2 | H6011-Group 2 | H6012 - Group 2 | H6008-Group 2 | H6013-Group 2 |
|---|---|---|---|---|---|---|---|---|---|---|---|---|---|---|---|
| | | Name | Johannes | Francis | Wassily | Éduard | Gustav | Claude | René | Pablo | Edvard | Diego | Andy | Sandro | Vincent |
| **H1** | Explore arena + Startbox | | X | X | X | X | X | X | X | X | X | X | X | X | X |
| **H2** | Explore arena + Startbox (1' sb + 2' arena) | | X | X | X | X | X | X | X | X | X | X | X | X | X |
| **H3** | GO OUT, DIG AND EAT IN STARTBOX | | X | X | X | X | X | X | X | X | X | X | X | X | X |
| **H4** | GO OUT, DIG AND EAT IN STARTBOX | | X | X | X | X | X | X | X | X | X | X | X | X | X |
| **EGO NO CUES** | | | | | | | | | | | | | | | |
| **Session** | | | | | | | | | | | | | | | |
| **1** | 1 OPEN sw IN SAMPLE | | E - 5 | S - 3 | W - 3 | S - 1 | W - 4 | S - 3 | W - 2 | S - 1 | W - 6 | E - 6 | E - 5 | E - 6 | S - 2 |
| **2** | | | S - 1 | W - 4 | N - 2 | E - 6 | S/E - 1 | W/S - 4 | W/E - 6 | S/W - 2 | S - 4 | S - 2 | S - 6 | E/S - 2 | W/S - 6 |
| **3** | | | E - 3 | E - 1 | S - 5 | S - 4 | E/S - 3 | S/E - 1 | S/E - 6 | W/S - 4 | E - 1 | W - 5 | E - 5 | W/E - 6 | S/E - 3 |
| **4** | 3 TRIALS IN SAMPLE AND CHOICE (FROM NOW ON) | | NNN/NNN - 4 | SSS/SSS - 6 | EEE/EEE - 1 | WWW/WWW - 5 | SWS/WSW - 4 | SWE/SSW - 5 | ESS/SWS - 5 | EWE/SWS - 1 | EEE/EEE - 1 | EEE/EEE - 3 | SSS/SSS - 2 | SWS/WEE - 3 | WES/EWS - 1 |
| **5** | | | NNN/NNN - 5 | SSS/SSS - 2 | EEE/EEE - 6 | EEE/EEE - 2 | SSW/ESS - 5 | WEE/WES - 2 | EWE/SSE - 6 | SWS/EWW - 2 | EEE/EEE - 6 | NNN/NNN - 4 | SSS/SSS - 1 | WEW/EES - 4 | SES/WSE - 3 |
| **6** | | | WWW/WWW - 2 | WWW/WWW - 5 | NNN/NNN - 1 | SSS/SSS - 3 | WES/SSE - 2 | WEE/SWS - 3 | SSE/WSE - 2 | WSE/ESW - 6 | NNN/NNN - 1 | EEE/EEE - 6 | SSS/SSS - 5 | SEW/SWS - 6 | EES/WSE - 5 |
| **7** | | | SSS/SSS - 3 | NNN/NNN - 1 | *Terminated for health reasons* | EEE/EEE - 6 | SSE/WEW - 3 | SSW/ESE - 1 | WEW/SES - 1 | SWS/EWS - 6 | WWW/WWW - 2 | NNN/NNN - 1 | WWW/WWW - 4 | ESE/WSE - 4 | SWE/EWS - 2 |
| **8** | TEST | TEST | WWW/WWEE - 5(2) | SSS/SSNN - 3(4) | | SSS/SSNN - 2(5) | WEW/SWSE - 5 | EEW/SESW - 3 | SSE/SWEW - 6 | EWE/WSES - 2 | EEE/EEWW - 6(1) | EEE/EEWW - 4(3) | SSS/SSNN - 5(2) | WEW/SESW - 4 | SWS/ESWE - 6 |
| **9** | | | EEE/EEE - 1 | EEE/EEE - 5 | | NNN/NNN - 4 | EES/EWW - 1 | SEE/SWW - 3 | SWE/WSS - 5 | SWE/ESW - 4 | SSS/SSS - 3 | WWW/WWW - 2 | WWW/WWW - 6 | SES/EWS - 2 | WSW/ESW - 6 |
| **10** | | | SSS/SSS - 6 | EEE/EEE - 2 | | EEE/EEE - 5 | SEW/WES - 6 | WEE/WEE - 2 | WSE/SWW - 4 | SEW/WES - 5 | NNN/NNN - 4 | SSS/SSS - 1 | NNN/NNN - 5 | SWE/WSW - 1 | WSW/ESE - 4 |
| **11** | | PROBE | SSS/**S**SSS - 4 | WWW/**W**WWW - 6 | | NNN/**N**NNN - 3 | WEW/**S**SWE - 4 | SES/**E**EWS - 6 | SSW/**S**SES - 5 | SWS/**W**WSE - 3 | EEE/**E**EEE - 5 | WWW/**W**WWW - 1 | SSS/**S**SSS - 2 | SEW/**W**WSE - 1 | ESE/WWSW - 3 |
| **12** | | | NNN/NNN - 3 | WWW/WWW - 1 | | SSS/SSS - 2 | WEW/SWW - 2 | ESE/SES - 4 | WSW/ESE - 3 | ESW/WSE - 1 | EEE/EEE - 2 | NNN/NNN - 5 | EEE/EEE - 1 | ESS/SWE - 5 | SEE/SEE - 2 |
| **13** | | | SSS/SSS - 4 | EEE/EEE - 3 | | EEE/EEE - 2 | EWE/SWS - 3 | WEW/SES - 4 | ESE/WSW - 5 | SSW/ESE - 2 | SSS/SSS - 3 | WWW/WWW - 6 | WWW/WWW - 2 | WSW/ESW - 6 | WSS/SEW - 5 |
| **14** | | | WWW/WWW - 5 | SSS/SSS - 3 | | WWW/WWW - 6 | SWE/WWS - 2 | WWE/SWS - 3 | ESE/WWS - 1 | WSE/WSW - 6 | WWW/WWW - 1 | NNN/NNN - 2 | EEE/EEE - 3 | WSW/ESE - 4 | WES/EWS - 1 |
| **15** | | | SSS/SSS - 1 | EEE/EEE - 2 | | NNN/NNN - 5 | SWS/ESE - 1 | ESW/WEE - 2 | ESS/SEW - 6 | ESE/WSW - 5 | NNN/NNN - 6 | WWW/WWW - 3 | NNN/NNN - 4 | ESE/WSW - 3 | WWS/SWE - 1 |
| **16** | TEST | TEST | SSS/SSNN - 3(4) | WWW/WWEE - 5(2) | | NNN/NNSS - 5(2) | SES/WSWS - 3 | SWS/ESEW - 5 | ESE/WEWS - 1 | WSW/ESEW - 5 | SSS/SSNN - 1(6) | EEE/EEWW - 4(3) | EEE/EEWW - 2(5) | SES/WSWE - 4 | SWS/SEEW - 6 |
| **17** | | | NNN/NNN - 6 | NNN/NNN - 1 | | WWW/WWW - 1 | ESE/EWS - 6 | SES/WEW - 1 | WEW/SWE - 2 | SES/WSW - 1 | EEE/EEE - 5 | SSS/SSS - 2 | NNN/NNN - 6 | SWS/EES - 2 | SSW/WSE - 2 |
| **18** | CUES FOR EGO/CURTAINS FOR ALLO IN CHOICE | CUES | SSS/SSS - 4 | SSS/SSS - 6 | | EEE/EEE - 3 | WES/SWEs - 4 | SES/SSWe - 6 | SWS/EWSw - 6 | SSW/WESw - 2 | WWW/WWW - 6 | WWW/WWW - 5 | NNN/NNN - 4 | EES/WSEw - 5 | EWS/SEWs - 4 |
| **19** | | | NNN/NNN - 3 | NNN/NNN - 1 | | SSS/SSS - 4 | EES/SEW - 3 | WES/EWS - 1 | WWS/SEE - 2 | SEW/WWS - 4 | EEE/EEE - 2 | SSS/SSS - 2 | EEE/EEE - 2 | SWE/SSW - 2 | ESS/SWE - 5 |

| | | | | | | | | | | | | | | | |
|---|---|---|---|---|---|---|---|---|---|---|---|---|---|---|---|
| 20 | | | EEE/EEE - 1 | WWW/WWW - 2 | | SSS/SSS - 2 | SWE/ESW - 2 | SES/WSW - 3 | WES/SEW - 1 | WSE/SEW - 2 | NNN/NNN - 4 | NNN/NNN - 1 | WWW/WWW - 5 | WWE/SSW - 1 | WSW/ESW - 6 |
| 21 | | | WWW/WWW - 6 | EEE/EEE - 5 | | WWW/WWW - 6 | WES/WES - 6 | EES/WSE - 5 | ESE/WSE - 4 | ESE/SWE - 6 | SSS/SSS - 3 | EEE/EEE - 4 | SSS/SSS - 1 | SSW/ESE - 4 | WSW/ESE - 4 |
| 22 | | PROBE | SSS/**s**SSS - 3 | SSS/**s**SSS - 6 | | EEE/**E**EEE - 1 | SSW/SWSE - 3 | SWS/**S**SWE - 6 | EWE/**E**ESW - 5 | ESW/**W**WSE - 2 | NNN/**n**NNN - 5 | WWW/**W**WWW - 2 | NNN/**N**NNN - 4 | WSE/**E**ESW - 1 | ESE/WWSW - 3 |
| 23 | | | NNN/NNN - 4 | WWW/WWW - 1 | | SSS/SSS - 5 | ESE/SWS - 4 | WEW/SES - 1 | WWS/ESE - 2 | SSW/WSE - 5 | EEE/EEE - 2 | NNN/NNN - 6 | EEE/EEE - 3 | WES/SSE - 6 | SWW/WSE - 1 |
| 24 | TEST | TEST | SSS/SSNN - 3(4) | EEE/EEWW - 5(2) | | NNN/NNSS - 2(5) | SES/WSWE - 3 | SWS/EWES - 6 | WSW/EWES - 5 | EWW/SWSE - 3 | WWW/WWEE - 5(2) | SSS/SSNN - 1(6) | EEE/EEWW - 6(1) | WEW/SWSE - 1 | SWS/ESEW - 2 |
| 25 | CURTAINS | CURTAINS | WWW/WWww - 2 | SSS/SSss - 5 | | EEE/EEee - 6 | EWE/SWEs - 2 | WEW/SEWe - 5 | SWW/SWWe - 1 | SWS/ESEw - 2 | SSS/SSss - 1 | SSS/SSss - 3 | SSS/SSss - 3 | ESE/WSEw - 4 | WWE/SSWe - 1 |
| 26 | | | EEE/EEE - 6 | NNN/NNN - 2 | | WWW/WWW - 1 | EWS/SWE - 5 | SSW/WSE - 2 | SWE/WSE - 3 | WWS/SWE - 6 | NNN/NNN - 3 | EEE/EEE - 1 | NNN/NNN - 4 | SEE/WSE - 5 | SWE/ESS - 4 |
| | HABITUATION TO FIBERS AND TEST OPTOGENETIC SETUP | | | | | | | | | | | | | | |
| 27 | LIGHT ON | inactivation SB | EEE/EEE - 5 | EEE/EEE - 2 | | SSS/SSS - 2 | WEE/WSE - 6 | SWS/WES - 2 | EWW/EWS - 5 | SEE/WWS - 5 | EEE/EEE - 5 | NNN/NNN - 2 | SSS/SSS - 3 | SES/WSW - 2 | ESE/SEW - 3 |
| 28 | LIGHT OFF | | WWW/WWW - 2 | SSS/SSS - 5 | | WWW/WWW - 1 | ESE/ESS - 1 | WES/EWS - 5 | SWE/SES - 4 | SSW/WEE - 2 | WWW/WWW - 6 | WWW/WWW - 1 | WWW/WWW - 1 | WSW/SEE - 1 | SWW/WSE - 1 |
| 29 | LIGHT ON | inactivation SB | NNN/NNN - 1 | NNN/NNN - 6 | | EEE/EEE - 6 | SWS/EWS - 5 | SEE/SWE - 6 | SWW/ESW - 2 | WSW/ESE - 1 | EEE/EEE - 1 | SSS/SSS - 4 | EEE/EEE - 6 | SEE/SEW - 5 | ESE/SWE - 2 |
| 30 | LIGHT OFF | | *Terminated for health reasons* | SSS/SSS - 5 | | SSS/SSS - 2 | SEE/SEE - 6 | EWS/SEW - 2 | SWW/EES - 2 | ESE/WES - 5 | SSS/SSS - 5 | EEE/EEE - 1 | WWW/WWW - 2 | SEW/SEW - 6 | WWS/WSW - 3 |
| 31 | LIGHT ON | inactivation ARENA | | EEE/EEE - 2 | | WWW/WWW - 5 | ESW/SES - 3 | SWW/ESE - 1 | SES/SWE - 4 | SEE/SWS - 2 | WWW/WWW - 6 | NNN/NNN - 2 | WWW/WWW - 1 | SWE/WSE - 1 | SWE/SWE - 1 |
| 32 | LIGHT OFF | | | SSS/SSS - 1 | | WWW/WWW - 1 | EWS/ESE - 5 | SWS/SES - 6 | ESE/SES - 5 | SWW/SES - 1 | EEE/EEE - 1 | EEE/EEE - 1 | SSS/SSS - 3 | EWW/SSW - 5 | WSE/ESE - 2 |
| 33 | LIGHT ON | inactivation ARENA | | SSS/SSS - 1 | | EEE/EEE - 6 | SEE/WEW - 6 | SWS/EWS - 5 | EWS/WES - 1 | ESE/WSW - 4 | SSS/SSS - 5 | WWW/WWW - 1 | WWW/WWW - 2 | ESE/WEW - 2 | SEE/WSW - 4 |
| 34 | LIGHT OFF | | | NNN/NNN - 6 | | WWW/WWW - 5 | WSE/WSS - 3 | SWW/WEW - 1 | WSE/ESW - 2 | ESW/SES - 4 | EEE/EEE - 5 | SSS/SSS - 4 | EEE/EEE - 6 | WEE/ESS - 6 | SEE/EES - 4 |
| 35 | LIGHT OFF | PROBE | | NNN/**N**NNN - 5 | | WWW/**W**WWW - 6 | SWS/**S**ESW - 6 | EWE/**W**WSE - 2 | WSW/SSWE - 3 | EWS/EESW - 5 | SSS/**S**SSS - 3 | EEE/**E**EEE - 2 | EEE/**E**EEE - 1 | WSE/**S**ESW - 4 | WEW/**S**SES - 1 |
| 36 | | | | WWW/WWW - 3 | | NNN/NNN - 3 | WSS/ESE - 2 | WSE/WSW - 3 | WES/SWS - 5 | WWS/ESW - 1 | NNN/NNN - 2 | NNN/NNN - 5 | WWW/WWW - 3 | SES/WSW - 1 | WSE/SEE - 5 |
| 37 | LIGHT ON | PROBE INACTIVATIO N | | EEE/**E**EEE - 2 | | NNN/**N**NNN - 5 | ESE/**W**WSE - 4 | WES/**E**ESW - 5 | ESE/WWSW - 6 | SEE/WWSE - 6 | EEE/**E**EEE - 1 | EEE/**E**EEE - 6 | WWW/**W**WWW - 3 | SWS/**E**SWE - 3 | SSE/WSSE - 2 |

#### Supplementary Table 2 (continued)

| | 2300hZ, DELAY 5S, 0.5 S DURATION | | H6016 - group 3 | H6017 - group 3 | H6024 - group 3 | H6029 - group 3 | H6014 - group 3 | H6015 - group 3 | H6018 - group 3 | H6020 - group 3 | H6019 - group 4 | H6021 - group 4 | H6022 - group 4 | H6023 - group 4 | H6026 - group 4 | H6027 - group 4 | H6025 - group 4 | H6028 - group 4 |
|---|---|---|---|---|---|---|---|---|---|---|---|---|---|---|---|---|---|---|
| | | | Salvador | Camille | Marc | Jan | Piet | Henri | Roy | Pierre-Auguste | Jean Auguste | Raffaello | Caspar | Edgar | Amedeo | Peter | Michelangelo | Leonardo |
| **H1** | Explore arena + Startbox | | X | X | X | X | X | X | X | X | X | X | X | X | X | X | X | X |
| **H2** | Explore arena + Startbox (1' sb + 2' arena) | | x | x | X | X | X | X | X | X | X | X | X | X | X | X | X | X |
| **H3** | GO OUT, DIG AND EAT IN STARTBOX | | x | x | x | X | X | X | X | X | X | X | X | X | X | X | X | X |
| **H4** | GO OUT, DIG AND EAT IN STARTBOX | | x | x | x | X | X | X | X | X | X | X | X | X | X | X | X | X |
| **EGO NO CUES** | | | | | | | | | | | | | | | | | | |
| **Session** | | | | | | | | | | | | | | | | | | |
| **1** | 1 OPEN sw IN SAMPLE | | E - 6 | N - 3 | W - 2 | S - 1 | E - 6 | E - 3 | W - 2 | S - 1 | W - 5 | N - 1 | E - 6 | W - 2 | W - 5 | E - 6 | S - 2 | S - 1 |
| **2** | | | S - 1 | W - 4 | N - 3 | W - 6 | S - 1 | W - 4 | W - 3 | W - 6 | S - 2 | S - 4 | S - 1 | W - 3 | S - 2 | S - 4 | W - 6 | W - 6 |
| **3** | | | E - 3 | E - 1 | S - 5 | N - 4 | E - 3 | E - 1 | S - 5 | E - 5 | W - 6 | E - 2 | E - 3 | N - 5 | W - 6 | E - 6 | S - 3 | W - 2 |
| **4** | 3 TRIALS IN SAMPLE AND CHOICE (FROM NOW ON) | | NNN/NNN - 4 | SSS/SSS - 6 | EEE/EEE - 1 | WWW/W WW - 5 | SWS/WS W - 4 | SWE/SSW - 5 | ESS/WS S - 1 | EWE/SW S - 5 | EEE/EEE - 3 | NNN/NNN - 2 | NNN/NNN - 4 | EEE/EEE - 1 | SWS/WE E - 3 | ESS/SW E - 5 | WES/EW S - 1 | EWE/SW S - 5 |
| **5** | | | NNN/NNN - 5 | SSS/SSS - 2 | EEE/EEE - 6 | EEE/EEE - 2 | SSW/ES S - 5 | terminated for health reasons | EWE/SS E - 6 | SWS/EW W - 2 | WWW/W WW - 4 | EEE/EEE - 3 | EEE/EEE - 5 | EEE/EEE - 6 | WEW/EE S - 4 | WSW/W ES - 6 | SES/WS W - 3 | SWS/EW W - 2 |
| **6** | | | WWW/W WW - 2 | WWW/WW W - 5 | WWW/WW W - 5 | EEE/EEE - 6 | WES/SS E - 2 | | SSE/WS E - 2 | WSE/ES W - 3 | EEE/EEE - 6 | SSS/SSS - 5 | WWW/WW W - 2 | NNN/NNN - 1 | SEW/SW S - 6 | WSW/ES E - 4 | EES/WS W - 5 | WSW/W SW - 3 |
| **7** | | | SSS/SSS - 3 | NNN/NNN - 1 | WWW/WW W - 2 | SSS/SSS - 4 | SSE/WE W - 3 | | WEW/SE S - 1 | EWS/EW S - 6 | NNN/NNN - 4 | WWW/WW W - 4 | SSS/SSS - 3 | WWW/WW W - 2 | ESE/WS E - 4 | ESE/WS W - 3 | SEW/EW S - 2 | SWS/EW S - 6 |
| **8** | TEST | TEST | EEE/EEW W - 6(1) | SSS/SSN N - 5(2) | WWW/WW EE - 2(5) | NNN/NNN - 3(4) | WEW/S WSE - 5 | | SSE/WS WE - 6 | EWE/SW SE - 2 | EEE/EEW W - 4(3) | SSS/SSN N - 5(2) | WWW/WW EE - 2(5) | SSS/SSN N - 3(4) | WEW/SE SW - 4 | WWE/SE SW - 1 | SWS/ES WE - 6 | EWE/SW SE - 2 |
| **9** | | | WWW/W WW - 1 | EEE/EEE - 5 | SSS/SSS - 3 | WWW/W WW - 6 | EES/EW W - 1 | | WWE/W SS - 3 | SWE/ES W - 4 | WWW/W WW - 2 | WWW/WW W - 6 | EEE/EEE - 1 | EEE/EEE - 5 | SES/WE S - 2 | SSW/S WW - 5 | WSW/ES W - 6 | SWE/ES W - 4 |
| **10** | | | SSS/SSS - 6 | EEE/EEE - 2 | NNN/NNN - 4 | EEE/EEE - 5 | SEW/WE S - 6 | | SEE/SW W - 4 | WES/WE S - 5 | SSS/SSS - 1 | NNN/NNN - 5 | SSS/SSS - 6 | EEE/EEE - 2 | SWE/SW S - 1 | EEW/SW W - 2 | WSS/ES E - 4 | EES/EW W - 1 |
| **11** | | PROBE | SSS/**S**SS S - 4 | WWW/**W** WWW - 6 | EEE/**E**EEE - 5 | NNN/**N**N NN - 3 | WWE/**S**S WE - 4 | | SSW/**E**E ES - 5 | SWS/**S**S SW - 3 | WWW/**W** WWW - 1 | SSS/**S**SS S - 2 | SSS/**S**SS S - 4 | WWW/**W** WWW - 6 | SEW/**W** WSE - 1 | SSE/SS EE - 4 | ESW/WS WS - 1 | SWE/ES ES - 2 |
| **12** | | | NNN/NNN - 3 | WWW/WW W - 1 | EEE/EEE - 2 | SSS/SSS - 2 | EEW/SE S - 3 | | ESE/WS W - 2 | WES/SE E - 2 | NNN/NNN - 5 | EEE/EEE - 1 | NNN/NNN - 3 | WWW/WW W - 1 | ESS/SW E - 5 | SSE/WS E - 1 | SEE/SEE - 2 | WEW/SE S - 4 |
| **13** | | | EEE/EEE - 2 | NNN/NNN - 4 | SSS/SSS - 3 | SSS/SSS - 1 | WEW/S WW - 2 | | WSW/ES W - 3 | ESW/WS E - 1 | WWW/W WW - 6 | WWW/WW W - 2 | EEE/EEE - 2 | NNN/NNN - 4 | WSE/ES W - 6 | SES/WS E - 2 | WSS/SE W - 5 | ESE/SW E - 4 |
| **14** | | | WWW/W WW - 5 | SSS/SSS - 3 | WWW/WW W - 1 | WWW/W WW - 6 | SWE/W WS - 2 | | ESW/WS E - 1 | WSE/WE S - 6 | NNN/NNN - 3 | EEE/EEE - 3 | WWW/WW W - 5 | SSS/SSS - 3 | WSW/ES E - 4 | EWS/SW E - 4 | WES/EW S - 1 | WWE/S WS - 3 |
| **15** | | | SSS/SSS - 1 | EEE/EEE - 2 | NNN/NNN - 6 | NNN/NNN - 5 | SWS/SS E - 1 | | ESS/SE W - 6 | ESE/WS W - 5 | WWW/W WW - 4 | NNN/NNN - 4 | SSS/SSS - 1 | EEE/EEE - 2 | ESE/WE S - 3 | SSW/SS W - 4 | WWS/S WE - 1 | ESW/WE E - 2 |
| **16** | TEST | TEST | SSS/SSN N - 4(3) | WWW/WW EE - 2(5) | SSS/SSN N - 1(6) | NNN/NNS S - 5(2) | SES/WS WS - 3 | | ESE/WE WS - 1 | WSW/ES EW - 5 | EEE/EEW W - 3(4) | WWW/WW EE - 5(2) | SSS/SSN N - 3(4) | WWW/WW EE - 5(2) | SES/WS WE - 4 | WSS/ES ES - 3 | SWS/ES EW - 6 | SWS/ES EW - 5 |
| **17** | | | NNN/NNN - 6 | NNN/NNN - 1 | EEE/EEE - 5 | WWW/W WW - 1 | ESE/SW S - 6 | | WEW/S WE - 5 | SES/ES W - 1 | SSS/SSS - 2 | SSS/SSS - 6 | SSS/SSS - 2 | NNN/NNN - 6 | SWS/EE S - 2 | EWW/S WS - 2 | SSW/W SE - 2 | SES/WE W - 1 |
| **18** | CUES FOR EGO/CURTAINS FOR ALLO IN CHOICE | CUES | SSS/SSS - 4 | SSS/SSS - 6 | WWW/WW W - 6 | EEE/EEE - 3 | WEW/S WEs - 4 | | SWS/EW Sw - 6 | SSW/W SEw - 2 | WWW/W WW - 5 | NNN/NNN - 4 | WWW/WW W - 5 | SSS/SSS - 4 | EES/WS Ee - 5 | SWW/W SSe - 1 | EWS/SW Es - 4 | SES/SE We - 6 |

| | | | | | | | | | | | | | | | | | |
|---|---|---|---|---|---|---|---|---|---|---|---|---|---|---|---|---|---|
| **19** | | | NNN/NNN<br>- 3 | NNN/NNN<br>- 1 | EEE/EEE -<br>2 | SSS/SSS<br>- 4 | EES/WE<br>W - 5 | | WWS/SE<br>E - 2 | SEW/SW<br>S - 4 | SSS/SSS<br>- 2 | EEE/EEE -<br>2 | SSS/SSS -<br>2 | EEE/EEE -<br>2 | SWE/SS<br>W - 2 | WES/EW<br>E - 1 | ESS/SW<br>E - 5 | EES/WE<br>W - 3 |
| **20** | | | EEE/EEE -<br>1 | WWW/WW<br>W - 2 | NNN/NNN<br>- 3 | SSS/SSS<br>- 2 | SWE/ES<br>W - 2 | | WES/SE<br>W - 1 | WSE/SE<br>W - 2 | NNN/NNN<br>- 1 | WWW/WW<br>W - 5 | NNN/NNN<br>- 1 | WWW/WW<br>W - 5 | WWE/SS<br>W - 1 | EES/WS<br>W - 3 | WSW/ES<br>W - 6 | SWE/ES<br>W - 2 |
| **21** | | | WWW/W<br>WW - 6 | EEE/EEE -<br>5 | SSS/SSS -<br>4 | WWW/W<br>WW - 6 | WES/SE<br>S - 6 | | ESE/WS<br>E - 4 | ESE/EW<br>E - 6 | EEE/EEE -<br>4 | SSS/SSS -<br>1 | EEE/EEE -<br>4 | SSS/SSS -<br>1 | SSW/ES<br>E - 4 | EES/WS<br>E - 5 | WSW/ES<br>E - 4 | WES/WE<br>S - 6 |
| **22** | | PROBE | SSS/**s**SS<br>S - 3 | SSS/**s**SSS<br>- 6 | NNN/**n**NN<br>N - 5 | EEE/**E**EEE<br>- 1 | SSW/W<br>WSE - 3 | | EWE/**E**E<br>SW - 5 | ESW/**W**<br>WSE - 2 | WWW/**W**<br>WWW - 2 | NNN/**N**NN<br>N - 4 | NNN/**N**NN<br>N - 4 | WWW/**W**<br>WWW - 2 | WSE/**E**E<br>SW - 6 | SSE/SS<br>SE - 2 | ESE/WW<br>SW - 3 | SSE/WW<br>SE - 5 |
| **23** | | | NNN/NNN<br>- 4 | WWW/WW<br>W - 1 | EEE/EEE -<br>2 | SSS/SSS<br>- 5 | ESE/ESS<br>- 4 | | WWS/ES<br>E - 2 | SSW/W<br>SE - 5 | NNN/NNN<br>- 6 | EEE/EEE -<br>3 | EEE/EEE -<br>3 | NNN/NNN<br>- 6 | WES/SS<br>E - 2 | SWW/W<br>EE - 6 | SWW/W<br>SE - 1 | ESE/WS<br>S - 4 |
| **24** | TEST | TEST | SSS/SSN<br>N - 3(4) | EEE/EEW<br>W - 5(2) | WWW/WW<br>EE - 5(2) | NNN/NNS<br>S - 1(6) | SES/WS<br>WE - 3 | | WSW/E<br>WES - 5 | EWW/S<br>WSE - 3 | SSS/SSN<br>N - 1(6) | EEE/EEW<br>W - 6(1) | EEE/EEW<br>W - 6(1) | SSS/SSN<br>N - 1(6) | WEW/S<br>WSE - 1 | SWS/EW<br>ES - 4 | SWS/ES<br>EW - 2 | SES/WS<br>WE - 3 |
| **25** | CURTAINS | CURTAINS | WWW/W<br>Www - 2 | SSS/SSss<br>- 5 | SSS/SSss<br>- 1 | EEE/EEee<br>- 6 | EWE/SW<br>Es - 2 | | SWW/ES<br>We - 1 | SWS/SS<br>We - 6 | SSS/SSss<br>- 3 | SSS/SSss<br>- 3 | SSS/SSss<br>- 3 | SSS/SSss<br>- 5 | ESE/SS<br>Ws - 3 | WSE/SW<br>Sw - 2 | WWE/SS<br>We - 1 | EWE/SW<br>Ew - 2 |
| **26** | | | EEE/EEE -<br>5 | NNN/NNN<br>- 2 | NNN/NNN<br>- 4 | WWW/W<br>WW - 1 | EWS/SW<br>E - 5 | | SWE/WS<br>E - 3 | WWS/ES<br>W - 1 | EEE/EEE -<br>1 | NNN/NNN<br>- 4 | NNN/NNN<br>- 4 | EEE/EEE -<br>1 | SEE/WS<br>E - 5 | WWE/W<br>SS - 3 | SWE/ES<br>W - 4 | EWS/SW<br>E - 5 |
| | HABITUATION TO FIBERS AND TEST OPTOGENETIC SETUP | | | | | | | | | | | | | | | | | |
| **27** | LIGHT ON | inactivation SB | SSS/SSS<br>- 2 | EEE/EEE -<br>5 | NNN/NNN<br>- 2 | WWW/W<br>WW - 2 | SWS/WE<br>S - 2 | | SEW/W<br>WS - 5 | WSE/SE<br>E - 1 | WWW/W<br>WW - 1 | WWW/WW<br>W - 3 | WWW/WW<br>W - 1 | WWW/WW<br>W - 2 | SWW/W<br>SW - 1 | ESE/WS<br>E - 1 | SEE/WS<br>E - 1 | WSW/ES<br>W - 2 |
| **28** | LIGHT OFF | | WWW/W<br>WW - 1 | WWW/WW<br>W - 6 | WWW/WW<br>W - 1 | SSS/SSS<br>- 3 | SWS/EW<br>S - 5 | | SSW/WE<br>E - 2 | ESE/WE<br>W - 2 | EEE/EEE -<br>2 | EEE/EEE -<br>5 | EEE/EEE -<br>2 | EEE/EEE -<br>5 | ESE/SW<br>S - 2 | EWW/W<br>SS - 3 | EWW/W<br>SS - 3 | WSE/ES<br>E - 2 |
| **29** | LIGHT ON | inactivation SB | EEE/EEE -<br>6 | EEE/EEE -<br>1 | SSS/SSS -<br>4 | EEE/EEE -<br>6 | WSW/W<br>EW - 1 | | WSW/ES<br>E - 1 | EWS/SS<br>W - 5 | SSS/SSS<br>- 5 | WWW/WW<br>W - 2 | SSS/SSS -<br>5 | WWW/WW<br>W - 3 | ESE/SE<br>W - 3 | SWS/SS<br>E - 4 | SWS/ES<br>E - 5 | SSW/ES<br>E - 3 |
| **30** | LIGHT OFF | | SSS/SSS<br>- 2 | SSS/SSS -<br>5 | EEE/EEE -<br>1 | WWW/W<br>WW - 2 | EWS/SE<br>W - 2 | | ESE/WE<br>S - 5 | SWE/WE<br>S - 1 | WWW/W<br>WW - 1 | SSS/SSS -<br>6 | WWW/WW<br>W - 1 | SSS/SSS -<br>6 | SWE/SW<br>E - 1 | SWS/ES<br>E - 5 | ESW/WE<br>S - 1 | EWE/SW<br>S - 4 |
| **31** | LIGHT ON | inactivation ARENA | WWW/W<br>WW - 5 | WWW/WW<br>W - 6 | NNN/NNN<br>- 2 | WWW/W<br>WW - 1 | SEE/SW<br>E - 6 | | SES/SE<br>S - 2 | WEE/ES<br>S - 6 | EEE/EEE -<br>6 | EEE/EEE -<br>5 | EEE/EEE -<br>2 | EEE/EEE -<br>5 | SEE/EES<br>- 4 | WES/ES<br>E - 5 | SES/WS<br>W - 4 | WEE/SE<br>S - 1 |
| **32** | LIGHT OFF | | WWW/W<br>WW - 1 | EEE/EEE -<br>1 | EEE/EEE -<br>1 | WWW/W<br>WW - 1 | SWW/SE<br>W - 1 | | ESE/ES<br>W - 4 | SEE/ES<br>S - 5 | EEE/EEE -<br>6 | WWW/WW<br>W - 2 | EEE/EEE -<br>6 | WWW/WW<br>W - 2 | EES/SS<br>W - 3 | SES/SW<br>S - 4 | SES/SW<br>S - 4 | SWW/W<br>SE - 3 |
| **33** | LIGHT ON | inactivation ARENA | EEE/EEE -<br>6 | SSS/SSS -<br>5 | WWW/WW<br>W - 1 | SSS/SSS<br>- 3 | WES/EW<br>S - 5 | | EWS/SS<br>E - 4 | SES/WS<br>W - 2 | EEE/EEE -<br>2 | SSS/SSS -<br>6 | EEE/EEE -<br>6 | SSS/SSS -<br>6 | WSE/ES<br>E - 2 | EWS/SW<br>S - 3 | EWS/EW<br>S - 3 | EWS/SW<br>S - 1 |
| **34** | LIGHT OFF | | WWW/W<br>WW - 5 | EEE/EEE -<br>5 | SSS/SSS -<br>4 | EEE/EEE -<br>6 | SWS/SE<br>S - 6 | | SWW/SE<br>S - 1 | SEW/SE<br>W - 6 | SSS/SSS<br>- 5 | WWW/WW<br>W - 3 | SSS/SSS -<br>5 | WWW/WW<br>W - 3 | SEE/SW<br>E - 3 | ESW/EW<br>S - 1 | WES/ES<br>E - 5 | ESW/SE<br>S - 4 |
| **35** | LIGHT OFF | PROBE | WWW/**W**<br>WWW - 6 | SSS/**S**SS<br>S - 3 | EEE/**E**EEE<br>- 2 | EEE/**E**EEE<br>- 1 | EWE/**W**<br>WSE - 2 | | EWS/EE<br>SW - 5 | WSS/**S**S<br>EW - 4 | WWW/W<br>WW - 1 | SSS/SSS -<br>4 | WWW/WW<br>W - 1 | SSS/SSS -<br>4 | SEW/**S**S<br>ES - 1 | SWS/SS<br>EE - 6 | SWS/SS<br>EE - 6 | WSE/SS<br>EW - 5 |
| 36 | | | NNN/NNN<br>- 3 | NNN/NNN<br>- 1 | NNN/NNN<br>- 5 | WWW/W<br>WW - 3 | WSE/WS<br>W - 3 | | WWS/ES<br>W - 1 | SES/WS<br>W - 6 | EEE/EEE -<br>6 | EEE/EEE -<br>2 | EEE/EEE -<br>6 | EEE/EEE -<br>2 | ESE/SEE<br>- 5 | SWS/ES<br>E - 2 | SWS/ES<br>E - 2 | SWS/SE<br>E - 6 |
| **37** | LIGHT ON | PROBE INACTIVATI ON | NNN/**N**N<br>NN - 5 | EEE/**E**EEE<br>- 2 | EEE/**E**EEE<br>- 6 | WWW/**W**<br>WWW - 3 | WES/**E**E<br>SW - 5 | | SEE/WW<br>SE - 6 | SWS/**S**S<br>WE - 3 | SSS/SSS<br>- 4 | SSS/SSS -<br>1 | SSS/SSS -<br>4 | SSS/SSS -<br>1 | SSE/WW<br>SE - 2 | WSW/W<br>WSE - 1 | WSW/EE<br>SE - 1 | SWS/SS<br>EE - 4 |
